# Inhibition of the ubiquitin-proteasome system by a bioactivatable compound

**DOI:** 10.1101/2021.03.16.435589

**Authors:** Tatiana A. Giovannucci, Florian A. Salomons, Martin Haraldsson, Lotta H. M. Elfman, Malin Wickström, Patrick Young, Thomas Lundbäck, Jürgen Eirich, Mikael Altun, Rozbeh Jafari, Anna-Lena Gustavsson, John Inge Johnsen, Nico P. Dantuma

**Affiliations:** Department of Cell and Molecular Biology (CMB), Karolinska Institutet, Stockholm, Sweden; Chemical Biology Consortium Sweden (CBCS), Science for Life Laboratory, Division of Translational Medicine and Chemical Biology, Department of Medical Biochemistry and Biophysics, Karolinska Institutet, Solna, Stockholm, Sweden; Childhood Cancer Research Unit, Department of Women’s and Children’s Health, Karolinska Institutet, Stockholm, Sweden; Science for Life Laboratory, Department of Oncology-Pathology, Clinical Proteomics Mass Spectrometry, Karolinska Institutet, Solna, Stockholm, Sweden; Science for Life Laboratory, Department of Medical Biochemistry and Biophysics (MBB), Karolinska Institutet, Solna, Stockholm, Sweden; Science for Life Laboratory, Department of Laboratory Medicine, Karolinska Institutet, Solna, Stockholm, Sweden; Mechanistic Biology & Profiling, Discovery Sciences, R&D, AstraZeneca, Gothenburg, Sweden; Institute of Plant Biology and Biotechnology, University of Muenster 48143 Muenster, Germany

**Author notes:** Corresponding Author: Nico P. Dantuma, Department of Cell and Molecular Biology, Karolinska Institutet, Biomedicum, 7A, Solnavägen 9, SE-171 77 Stockholm, Sweden.

**Keywords:** ubiquitin-proteasome system, inhibitor, prodrug, drug development, bioactivation, NQO1

## Abstract

Malignant cells display an increased sensitivity towards drugs that reduce the function of the ubiquitin-proteasome system (UPS), which is the primary proteolytic system for destruction of aberrant proteins. Here, we report on the discovery of the bioactivatable compound CBK77, which causes an irreversible collapse of the UPS, accompanied by a general accumulation of ubiquitylated proteins and caspase-dependent cell death. CBK77 caused accumulation of ubiquitin-dependent, but not ubiquitin-independent, reporter substrates of the UPS, suggesting a selective effect on ubiquitin-dependent proteolysis. In a genome-wide CRISPR interference screen, we identified the redox enzyme NAD(P)H:quinone oxidoreductase 1 (NQO1) as a critical mediator of CBK77 activity, and further demonstrated its role as the compound bioactivator. Through affinity-based proteomics, we found that CBK77 covalently interacts with ubiquitin. In vitro experiments showed that CBK77-treated ubiquitin conjugates were less susceptible to disassembly by deubiquitylating enzymes. *In vivo* efficacy of CBK77 was validated by reduced growth of NQO1-proficient human adenocarcinoma cells in nude mice treated with CBK77. This first-in-class NQO1-activatable UPS inhibitor suggests that it may be possible to exploit the intracellular environment in malignant cells for leveraging the impact of compounds that impair the UPS.

## Introduction

Proteotoxic stress is a condition that is observed in many cancer types and can be capitalized in anti-cancer regimens^1^. The underlying causes of proteotoxic stress in malignant cells are multifactorial and inherently linked to their hyperactive state, such as an overall increase in protein synthesis and accumulation of aberrant and orphan proteins due to the loss of genome integrity^2^. Moreover, additional protein damaging factors, like elevated levels of radical oxygen species, increase the intracellular levels of damaged and misfolded proteins in malignant cells^3^. The ubiquitin-proteasome system (UPS) is the primary proteolytic system in charge of the timely and efficient degradation of aberrant and excessive proteins that are at risk of polluting the intracellular milieu^4^. Under physiological conditions, cells are endowed with a large excess of proteasomal activity, but the UPS can become rate limiting under proteotoxic stress conditions when the levels of substrates exceed the proteolytic capacity of the UPS^5^.

In malignant cells, the UPS is operating closer to the limits of its capacity and hence its operational efficiency is of critical importance for their survival^6^. The increased demand for clearance of misfolded proteins in malignant cells generates a therapeutic window at which reduction of UPS activity is lethal for cancer cells without severely affecting protein homeostasis in healthy cells^7^. To date, three proteasome inhibitors of the UPS have been clinically approved for treatment of multiple myeloma and mantle cell lymphoma^8^. Despite the overall success of proteasome inhibitors in the treatment of hematological malignancies, several remaining challenges, such as adverse side effects, drug resistance and poor activity towards solid tumors, motivate the exploration of alternative approaches for inhibiting UPS activity.

Proper functioning of the UPS involves hundreds of proteins, many of which display apparent druggable enzymatic activities^4^. Recent efforts have focused on the development of compounds that inhibit deubiquitylating (DUB) enzymes^9^ as well as the ubiquitin-selective unfoldase p97/valosin containing protein (VCP)^10^. Other experimental compounds target the process of ubiquitylation^11^ or ubiquitin itself^12, 13^. However, the complex and integrated nature of the UPS makes it hard to predict alternative strategies for obstructing this complex proteolytic pathway in cancer cells. Therefore, we opted for an unbiased, high-content phenotypic screen in a pursuit for small molecules that cause a general inhibition of the UPS by novel molecular mechanisms. This screen led to the discovery of the bioactivatable UPS inhibitor CBK77 that requires for its biological activity the presence of NAD(P)H:quinone oxidoreductase 1 (NQO1), a redox enzyme that is often upregulated in malignant cells^14^. We show that this first-in-class compound modifies ubiquitin conjugates, causes an irreversible global inhibition of ubiquitin-dependent proteasomal degradation, and induces caspase-dependent death of malignant cells. We propose that the elevated levels of NQO1 in malignant cells can be exploited for enhancing the selectivity of UPS inhibitors towards cancer cells.

## Results

### Cell-based, high-content screen for small molecule inhibitors of the UPS

With the aim of finding mechanistically novel UPS inhibitors, we performed a phenotypic screen for small molecules that cause a general accumulation of a UPS reporter substrate (**Fig. 1A**; **Suppl. Table S1**). For this purpose, we used a human melanoma cell line, MelJuSo, that stably expresses ubiquitin^G76V^-yellow fluorescent protein (Ub-YFP), a readily detectable fluorescent protein that is constitutively targeted for proteasomal degradation by the ubiquitin fusion degradation (UFD) pathway^15^. The Ub-YFP-expressing MelJuSo cells were treated for 16 hours with 10 µM solutions of a diverse set of 5720 small molecules from the Chemical Biology Consortium Sweden at the Karolinska Institute (www.cbcs.se). The set can be described as a chemically diverse representation of a larger collection of lead- and drug-like compounds, designed to afford straightforward screen follow-up through testing of close analogs to hit compounds. The screen resulted in the identification of the 1,3-thiazol compound CBK092352 (**Fig. 1B, Suppl. Fig. S1A**), which caused strong accumulation of the Ub-YFP substrate as well as an increase in apoptotic nuclei (**Suppl. Fig. S1B,C**).

**Figure 1.**
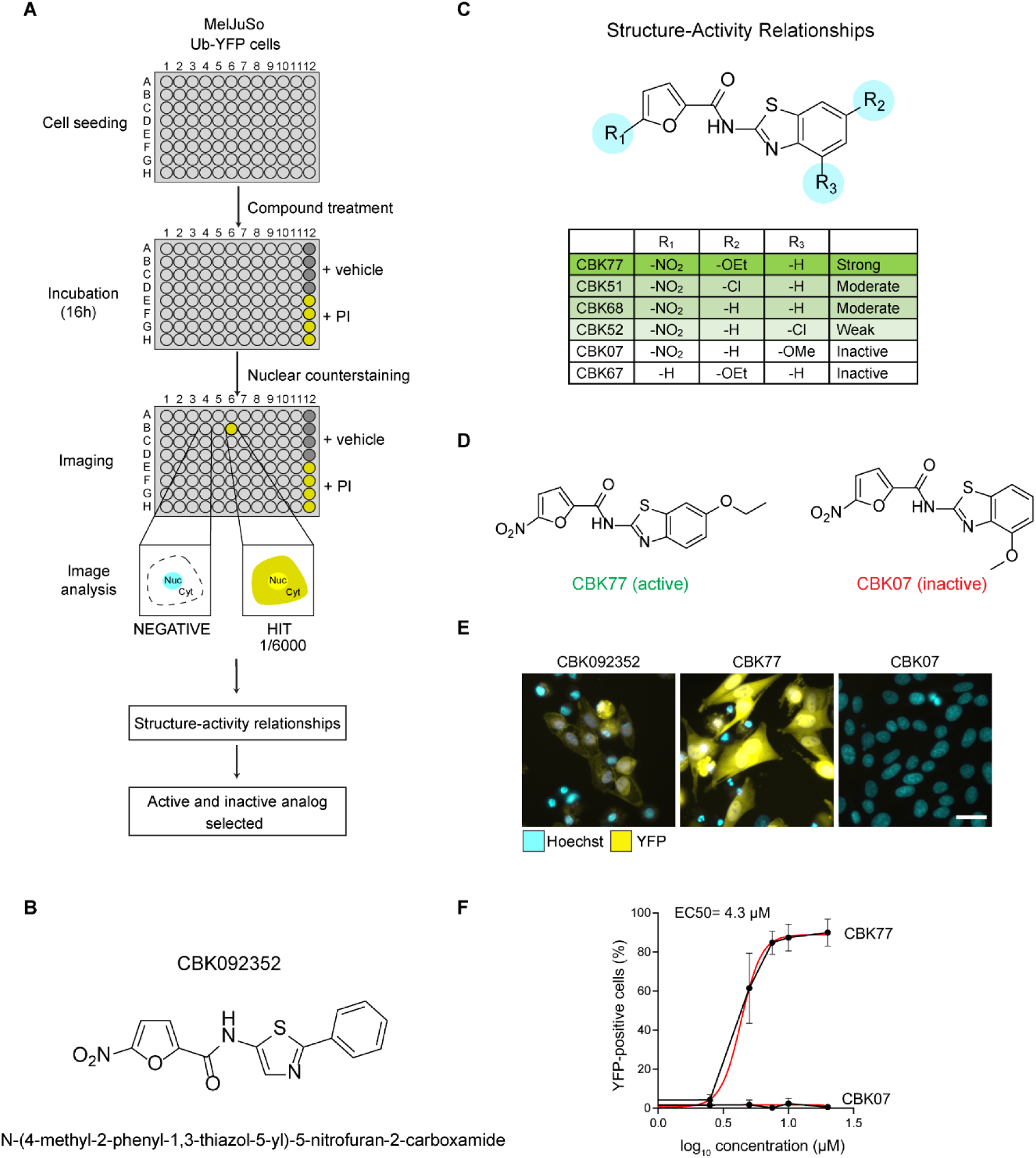
A cell-based phenotypic screen for inhibitors of the UPS. **A)** Representation of the screen workflow. Positive hits were defined as compounds that accumulated the UFD reporter in the nucleus above basal levels, defined by the signal in the DMSO-treated wells (negative control). Positive control wells treated with epoxomicin (proteasome inhibitor = PI, 100 nM) were also included in each plate of the screen. One positive hit was found in the initial screen, which formed the basis for a structure-activity relationship study (SAR). An active and an inactive structurally related compounds were selected for further characterization. **B)** Chemical structure of the hit compound CBK092352. **C)** Summary of structure activity relationships (SAR). Attenuation of activity is observed when R_3_ is substituted, whereas substituents in R_2_ are accepted. Removal of the nitro group results in an inactive compound. **D)** Structures of the analogs selected for further characterization, the active compound CBK77 and the inactive compound CBK07. **E)** Representative images of MelJuSo Ub-YFP cells treated for 16 hours with the indicated compounds (10 µM). DMSO at 0.1% was used as negative control. The nuclei were counterstained with Hoechst and cells imaged live with an automated widefield microscope. Scale bar = 20 µm. **F)** Concentration-response experiments performed with MelJuSo Ub-YFP cells. Cells were treated for 6 hours with a range of compound concentrations. Nuclei were stained with Hoechst and cells were directly imaged live with an automated widefield microscope. Data are represented as mean ± SD of three independent experiments. Non-linear curve fitting is depicted in red. The half-maximal effective concentration (EC_50_) upon CBK77 treatment is shown (4.3 µM, 95% confidence interval 3.8 – 5.0).

From the initial hit, we selected 41 analogs based on substructure and fingerprint similarity searches, as well as manual searches of the larger compound collection, to investigate a possible structure-activity relationship (SAR). This set of analogs was also further expanded through the synthesis of 19 structurally related compounds (**Suppl. Fig. S2A**; **Suppl. Table S2**). Most notably, SAR analysis showed that the 5-nitrofuran group is critical for biological activity of these compounds. Moreover, the O-ethoxy group at the 6 position (R_2_) improved the inhibitory effect of the compound as compared to the unsubstituted compound, whereas an O-methoxy group at the 4 position (R_3_) abrogated the effect (**Fig. 1C**). The importance of R_3_ was further strengthened by methyl and chlorine substitutions at this position, which also strongly reduced UPS inhibition and induction of cell death.

We selected a pair of closely related analogues of which one compound, CBK006377 (referred to as CBK77; N-[6-ethoxy-1,3-benzothiazol-2-yl]-5-nitrofuran-2- carboxamide), displayed profound UPS impairment and cellular toxicity, while the second compound, CBK085907 (referred to as CBK07; N-(4-methoxy-1,3-benzothiazol-2-yl)-5-nitrofuran-2-carboxamide), lacked these activities (**Fig. 1D,E**). The EC_50_ of CBK77 was determined as 4.3 µM (6 hour-treatment, 95% C.I 3.8 – 5.0 µM) with no detectable inhibition for CBK07 in the tested concentration range (**Fig. 1F**). It should, however, be mentioned that CBK07 is not completely inert as we observed modest UPS impairment and toxicity at high concentrations (>50 µM) over longer incubations (24 hours) (**Suppl. Fig. S2B**). The uptake of CBK77 and CBK07 in cells was comparable, eliminating the possibility that the strongly reduced activity of CBK07 could be attributed to a loss of cell permeability (**Suppl. Fig. S2C**). Together these data show that CBK77 blocks degradation of a reporter substrate of the UPS and induces cell death.

### CBK77 causes global impairment of the UPS

Following on the hit validation with the UFD substrate, we analyzed the effect of CBK77 on additional reporters that represent different classes of proteasome substrates: ubiquitin-arginine-GFP (Ub-R-GFP), a soluble reporter targeted for proteasomal degradation by the N-end rule pathway; YFP-CL1, which carries a C-terminal degradation signal that renders the reporter aggregation-prone and lastly, a reporter based on the T cell receptor subunit CD3δ (CD3δ-YFP), which is degraded by ER-associated degradation (ERAD)^16^. Administration of CBK77, but not CBK07, to three MelJuSo cell lines stably expressing these UPS reporters resulted in accumulation of each of these substrates, suggesting that CBK77 has a general inhibitory effect on the UPS (**Fig. 2A,B**).

**Figure 2.**
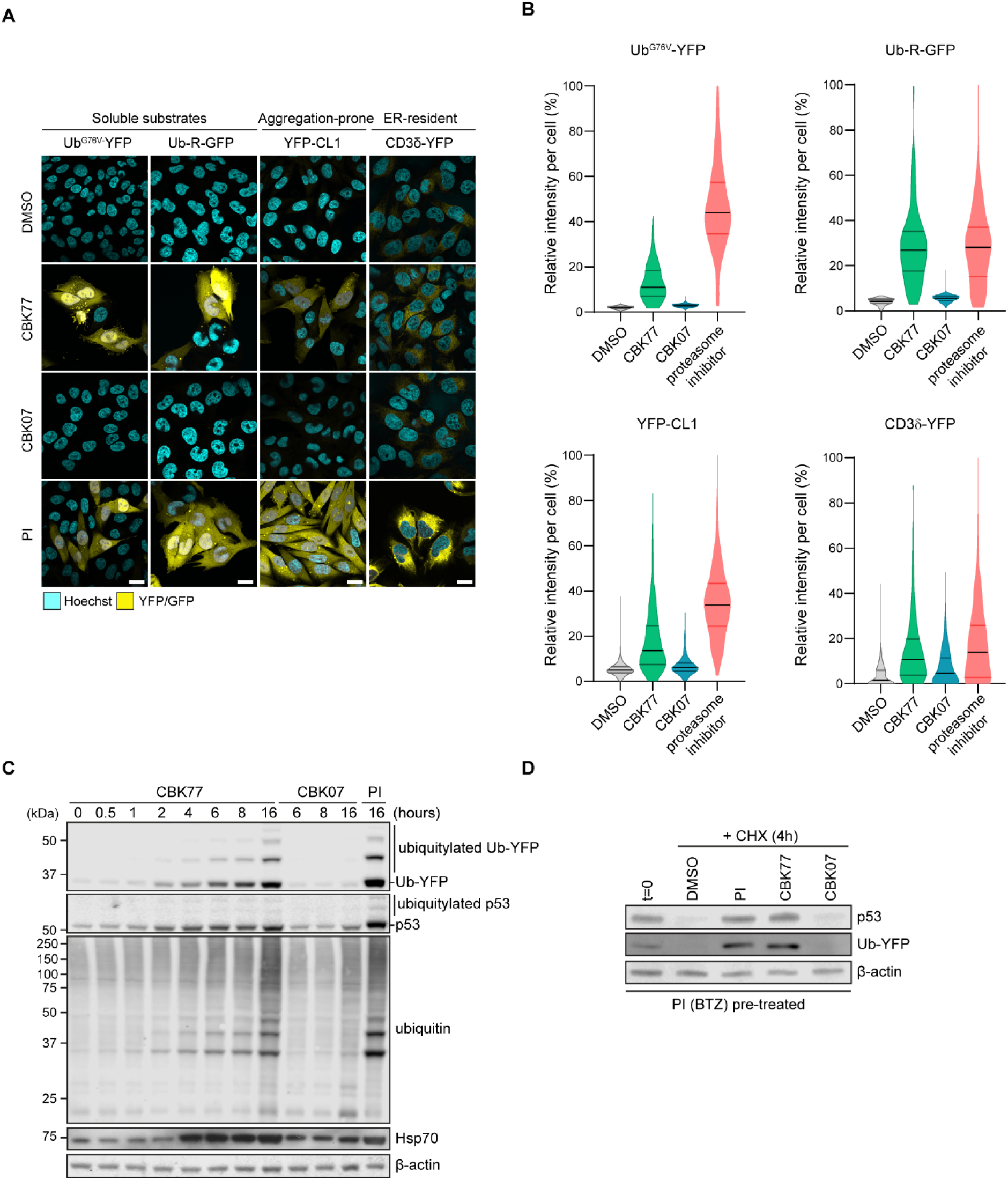
The UPS inhibitor CBK77 causes global impairment of the UPS. **A)** Representative confocal images of MelJuSo cells stably expressing the indicated fluorescent UPS reporters. Cells were treated for 16 hours with either CBK77 or CBK07 (10 µM) or bortezomib (proteasome inhibitor = PI, 25 nM) as a positive control. DMSO at 0.1% was used as negative control. Cells were fixed and nuclei were counterstained with Hoechst. Scale bars = 20 µm. **B)** Cells were treated for 16 hours with CBK77 or CBK07 (10 µM for all reporter cell lines besides YFP-CL1, treated at 5 µM). The proteasome inhibitor bortezomib (25 nM) and DMSO 0.1% were included as positive and negative controls, respectively. After nuclei counterstaining with Hoechst, cells were imaged in an automated manner with a widefield fluorescent microscope. Frequency and distribution of the reporter (YFP) intensity per cell are shown as violin plots from a representative experiment. Data are normalized to percentages were 0% = minimum value in the DMSO sample; 100% = maximum value in the proteasome inhibitor sample. Black lines within each distribution represent the median; colored lines represent the upper and lower interquartile range limits. Ub-YFP: n= 591, n= 283, n= 514, n= 337; Ub-R-GFP: n= 391, n=190, n= 201, n= 208; YFP-CL1: n= 666, n= 470, n= 623, n= 397; CD3δ-YFP: n= 717, n= 335, n= 505, n= 447 in DMSO, CBK77, CBK07 and bortezomib-treated cells, respectively, from a representative experiment. **C)** MelJuSo Ub-YFP cells were treated with either CBK77, CBK07 (10 µM) or epoxomicin (PI = proteasome inhibitor, 100 nM) and harvested at the indicated timepoints. Cell lysates were analyzed by immunoblotting with the indicated antibodies. **D)** MelJuSo Ub-YFP cells were pre-treated for 3 hours with the reversible proteasome inhibitor bortezomib (BTZ, 25 nM) to increase the levels of YFP substrate before the chase. Samples were taken directly after pretreatment (t=0). The remaining wells were co-treated with cycloheximide (CHX, 50 µg/ml) and the indicated compounds and harvested after 4 hours (CHX 4h). Cell lysates were analyzed by immunoblotting with the indicated antibodies.

Western blot analysis showed an increase in the levels of Ub-YFP as well as the endogenous UPS substrate p53 already 2 hours after administration of CBK77, while the steady-state levels of these proteins remained unaltered in CBK07-treated cells, even after 16 hours (**Fig. 2C**). Several high-molecular weight products were detected in both the Ub-YFP and p53 immunoblots, which gave rise to a ladder pattern that is characteristic for ubiquitin-modified proteins (**Fig. 2C**). Moreover, CBK77 caused a general accumulation of ubiquitin conjugates with similar kinetics as observed for polyubiquitylated Ub-YFP and p53 (**Fig. 2C**). In line with a general inhibition of ubiquitin-dependent degradation, CBK77 treatment increased the levels of Hsp70 (**Fig. 2C**). The chaperone Hsp70 is a general marker for activation of the heat-shock response, which is a common stress response elicited by inhibition of protein degradation^17^. Analysis of the turnover of Ub-YFP and p53 upon inhibition of protein synthesis with cycloheximide showed that the accumulation of these substrates was due to a delay in their clearance by proteasomal degradation (**Fig. 2D**). Together, these data show that CBK77 inhibits proteasomal degradation resulting in accumulation of polyubiquitylated substrates.

### CBK77 causes irreversible UPS impairment followed by caspase-dependent cell death

The UPS is critical for cell viability and blockade of the pathway typically results in induction of apoptosis^18^. Consistent with apoptotic cell death, administration of the pan-caspase inhibitor Q-VD-OPh significantly reduced cell death of CBK77-treated cells, in contrast to the compound necrostatin, which prevents necroptosis, a form of nonapoptotic cell death (**Fig. 3A**). It is noteworthy that the pan-caspase inhibitor did not prevent accumulation of the reporter substrate, in line with the model that induction of apoptosis is a consequence and not a cause of UPS impairment (**Fig. 3B**).

**Figure 3.**
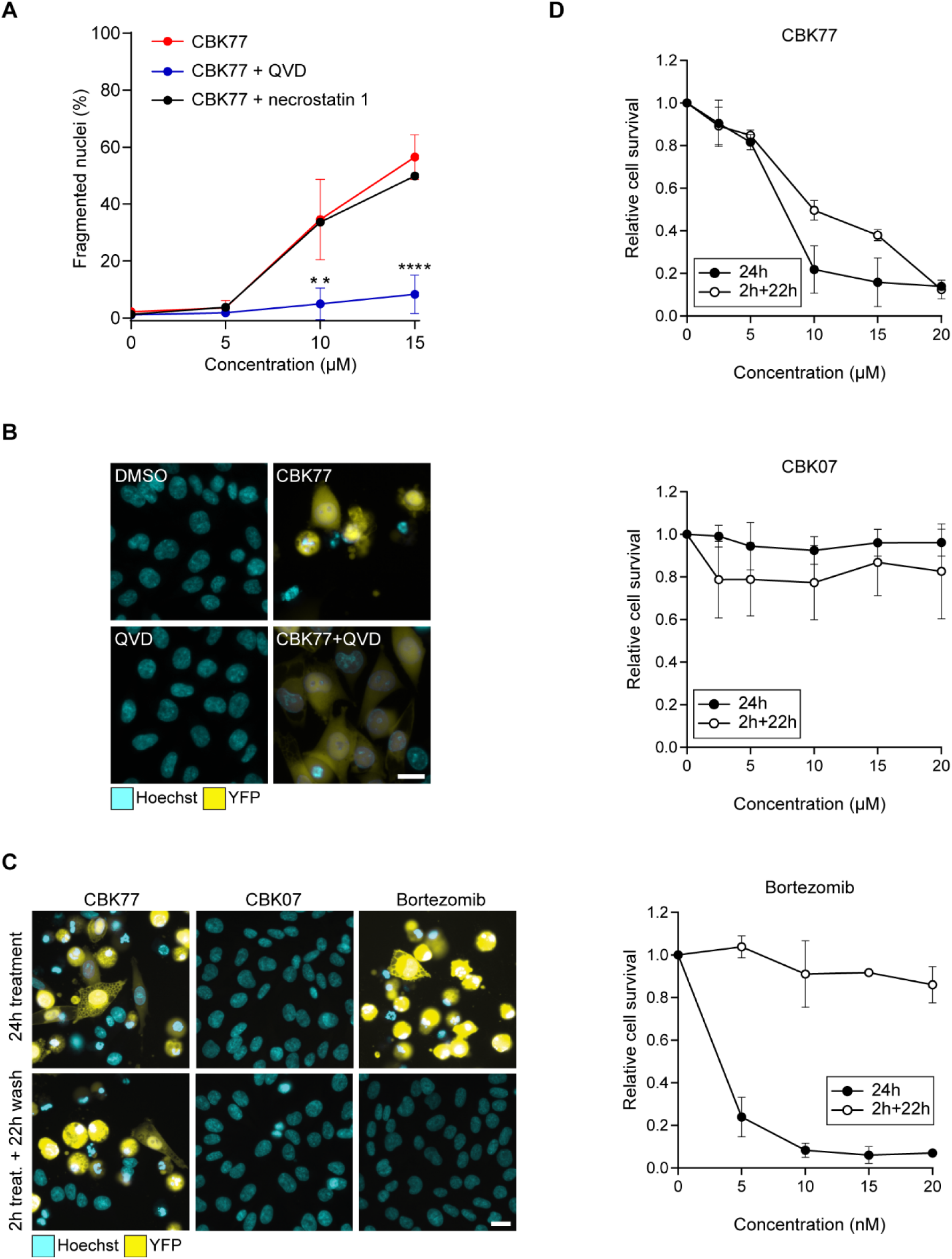
CBK77 causes irreversible UPS impairment followed by caspase-dependent cell death. **A)** MelJuSo Ub-YFP cells were treated with either 5, 10 or 15 µM of CBK77 with or without co-treatment with the pan-caspase inhibitor Q-VD-OPh (QVD, 20 µM) or necrostatin 1 (30 µM) for 24 hours. DMSO-treated cells (0.2 or 0.3%, respectively) were used as negative control. After incubation cells were fixed, counterstained with Hoechst and imaged with an automated widefield microscope. The number of apoptotic cells was determined in an automated manner based on nuclear staining intensity and fragmentation. Data are shown as mean percentage of apoptotic cells ± SD from three independent experiments. ** adjusted p ≤ 0.01, **** adjusted p ≤ 0.0001 (two-way ANOVA with Dunnett’s multiple comparisons test). **B)** Representative images from (**A**). Scale bar = 20 µm. **C)** Representative images from wash-out experiments performed in MelJuSo Ub-YFP cells. Cells were either continuously treated for 24 hours (“24 treatment”), or treated for 2 hours with DMSO, CBK77 (10 µM) or the reversible proteasome inhibitor bortezomib (25 nM) and further incubated with inhibitor-free medium up to 24 hours (“2h treat. + 22h wash”). Nuclei were stained with Hoechst. Imaging was then performed live with an automated widefield microscope. Scale bar = 20 µm. **D)** Quantification of (**C**). The cell count was obtained based on the nuclei staining. Data are shown as mean relative cell count to DMSO-treated cells ± SD from three technical replicates of a representative experiment.

Interestingly, a 2-hour treatment of cells with CBK77 followed by a washout and 22-hour incubation in the absence of the compound still caused a level of UPS inhibition that matched the inhibition that was observed in cells that had been continuously incubated with CBK77 for 24 hours, in contrast to results for the reversible proteasome inhibitor bortezomib, where a washout resulted in restoration of UPS activity (**Fig. 3C**). This finding shows that CBK77 has an irreversible effect on the functionality of the UPS. Accordingly, transient exposure to CBK77 was sufficient to induce cell death whereas cells were able to recover from a short incubation with bortezomib (**Fig. 3D**). Altogether, this shows that CBK77 induces irreversible inhibition of proteasomal degradation followed by caspase-dependent apoptosis.

### CBK77 does not cause global inhibition of proteasomal or DUB activity

Proteasome activity and DUB activity are two primary targets of compounds that have been reported to cause a general impairment of the UPS^8, 19^. Therefore, we explored if CBK77 executed its inhibitory effect by interfering with these enzymes.

Using an *in vitro* enzymatic assay, we analyzed the effect of CBK77 treatment on the chymotrypsin-like activity of the proteasome, which is the primary catalytic site and the main target of clinically used proteasome inhibitors^8^. *In vitro* analysis of the chymotrypsin-like activity in lysates of CBK77-treated cells showed that CBK77, in contrast to proteasome inhibitor, did not inhibit this activity (**Fig. 4A**). Moreover, contrary to the proteasome inhibitor epoxomicin, CBK77 did not impair the degradation of the ubiquitin-independent reporter ZsGreen ornithine decarboxylase (ODC)^20^ (**Fig. 4B,C**), further excluding that CBK77 impairs UPS activity at the level of the proteasome and suggesting that the effect of CBK77 is confined to ubiquitin-dependent proteasomal degradation.

**Figure 4.**
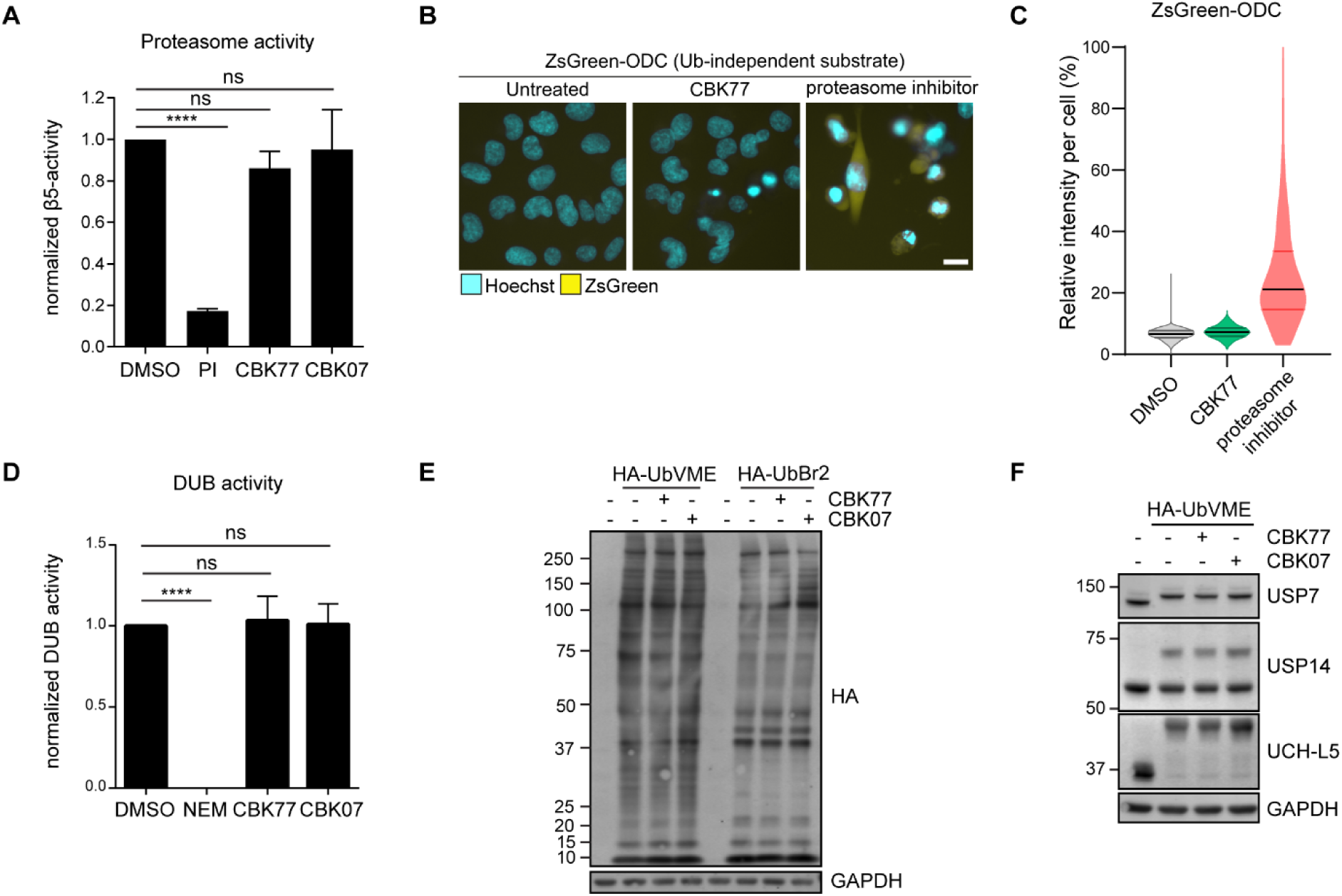
CBK77 does not cause global inhibition of proteasomal or DUB activity. **A)** MelJuSo parental cells were treated with DMSO 0.1%, CBK77 10 µM, CBK07 10 µM or epoxomicin (proteasome inhibitor = PI, 200 nM) for 4 hours. The chymotrypsin activity (β5 subunit) of the proteasome was assessed in the corresponding lysates by following conversion of the fluorogenic Suc-LLVY-AMC substrate over 1 hour. Data are shown as mean ± SD of three independent experiments. ns = non-significant. **** adjusted p-value ≤ 0.0001 (One-way ANOVA with Dunnett’s multiple comparisons test). Suc-LLVY-AMC = Suc-Leu-Leu-Val-Tyr-7-amido-4-methylcoumarin. **B)** Representative images of MelJuSo ZsGreen-ODC cells. Cells were treated for 16 hours with CBK77 (5 µM). The proteasome inhibitor epoxomicin (100 nM) was included as positive control. Nuclei were counterstained with Hoechst and the cells imaged live with an automated widefield microscope. Scale bar = 20 µm. **C)** Quantification of (**B**). The nuclear YFP intensity per cell was quantified using Cell Profiler. Frequency and distribution of the measured YFP intensity per cell are shown as violin plots. Data are normalized to percentages were 0% = minimum value in the DMSO sample; 100% = maximum value in the proteasome inhibitor sample. n > 500 cells (DMSO); n = 250 cells (CBK77) and n = 298 cells (proteasome inhibitor) from a representative experiment. Black lines within each distribution represent the median; colored lines represent the upper and lower interquartile range limits. **D)** MelJuSo Ub-YFP cells were treated with DMSO 0.1%, CBK77 10 µM or CBK07 10 µM for 4 hours. The global deubiquitylating activity in the cells was assessed in the corresponding lysates by following the conversion of the fluorogenic ubiquitin-AMC substrate over 30 minutes. A sample containing DMSO-treated cell lysate + 15 mM NEM was included as a negative control. Data are shown as mean ± SD of three independent experiments. ns = non-significant. **** adjusted p-value ≤ 0.0001 (One-way ANOVA with Dunnett’s multiple comparisons test). AMC = 7-amido-4-methylcoumarin; DUB = deubiquitylating enzyme; NEM = N-ethylmaleimide. **E)** MelJuSo parental cells were treated with DMSO 0.1%, CBK77 10 µM or CBK07 10 µM for 6 hours. The enzymatic activity of DUBs was assessed by incubating the cell extracts with the DUB-specific probes HA-UbVME and HA-UbBr2. A sample was incubated without probe as a technical control for DUB labeling. DUB activity was assessed by SDS-PAGE and western blotting with an anti-HA antibody. **F)** The same samples displayed in (**E**) were probed with antibodies against the specified enzymes. Active DUBs shift upwards in the gel due to the incorporation of the probe.

The defect in ubiquitin-dependent degradation and accumulation of ubiquitylated proteins could be due to inhibition of deubiquitylating (DUB) enzymes, as removal of ubiquitin chains is a prerequisite for proteasomal degradation of ubiquitylated substrates^21^. However, a role for inhibition of DUBs in the mode of action of CBK77 is unlikely as our analysis showed that CBK77 did not cause an overall reduction in DUB activity in cell lysates (**Fig. 4D**). Moreover, profiling of DUB activity with activity probes^22^ did not reveal apparent differences in the levels or activity of DUBs in lysates from cells treated with CBK77 (**Fig. 4E**). There was also no effect of CBK77 on the activity on USP7^23, 24^, USP14^25^ and UCH-L5^9^, three DUBs that we considered of specific interest as these have been found to be targeted by other experimental compounds (**Fig. 4F**). Thus, CBK77 causes a global impairment of ubiquitin-dependent proteasomal degradation without global inhibition of proteasome or DUB activity.

### NAD(P)H:quinone oxidoreductase 1 (NQO1) is critical for the CBK77-mediated UPS impairment

To gain further insight in the actual mode of action of CBK77, we performed an unbiased, genome-wide CRISPR interference screen to identify mechanisms of resistance. For this purpose, we transduced Cas9-expressing MelJuSo cells with a lentiviral library containing four targeting single guide RNA (sgRNA) per gene (**Fig. 5A**) and exposed them three times during a total period of five days to the IC_90_ concentration of CBK77 (25 µM) (**Fig. 5B**). After five days, cells were harvested, and samples subjected to barcode sequencing to identify sgRNAs that were significantly enriched in the CBK77-treated populations. Interestingly, a clear enrichment of guides targeting the oxidoreductase NQO1 was found in both replicates (**Fig. 5C, Suppl. Table S3**). Moreover, the gene encoding for Nrf2 (NFE2L2), a transcriptional stimulator of NQO1^26^, as well as the gene encoding for flavin adenine dinucleotide synthase 1 (FLAD1), which is responsible for synthesis of the NQO1 coenzyme flavin adenine dinucleotide (FAD)^27^, were highly enriched in the CBK77-resistant cell population (**Fig. 5C, Suppl. Table S3**).

**Figure 5.**
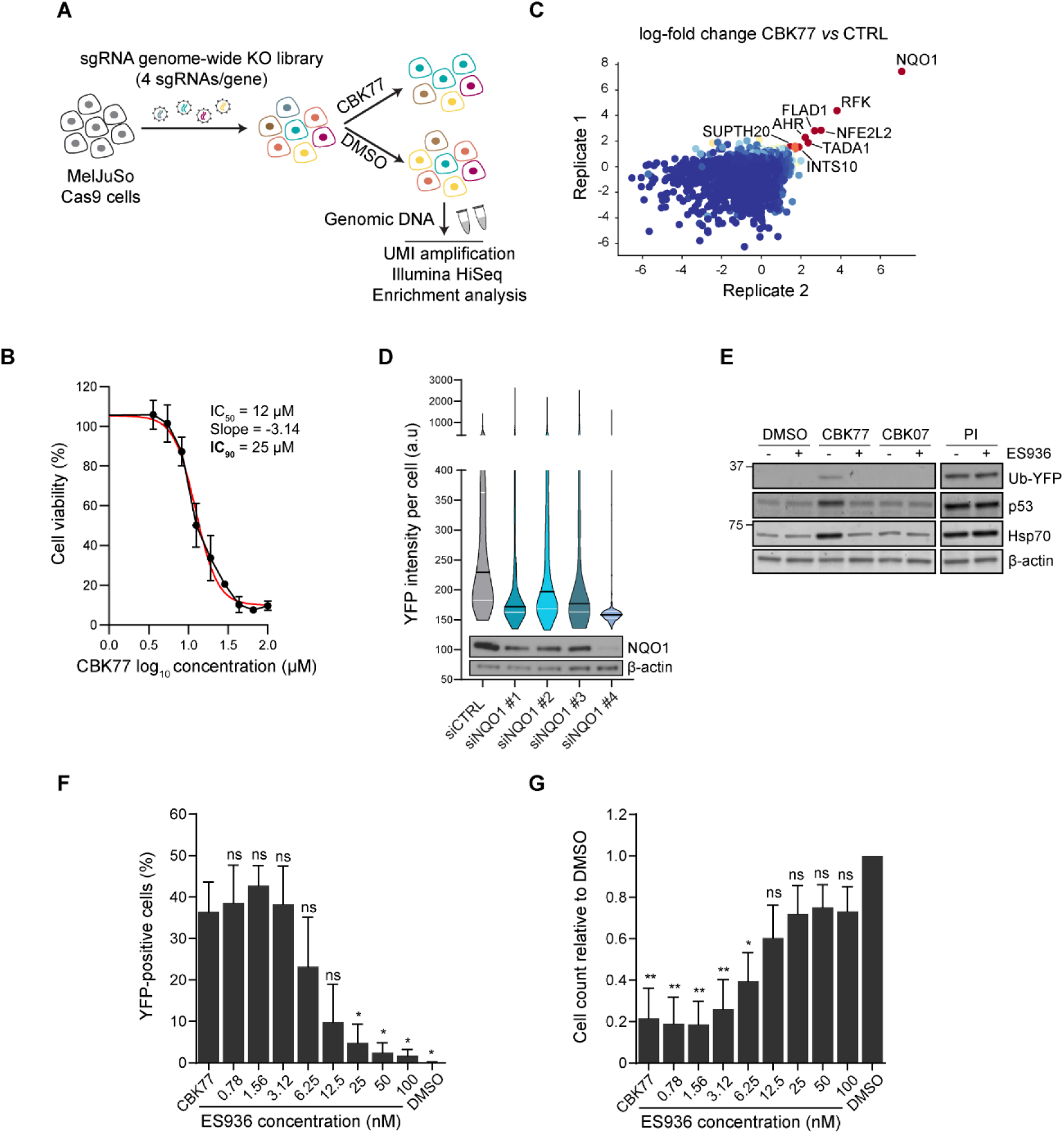
NAD(P)H:quinone oxidoreductase 1 (NQO1) is a critical factor for CBK77-mediated UPS impairment. **A)** Schematic drawing depicting the pooled CRISPR–Cas9 screening paradigm. MelJuSo cells expressing Cas9 were infected with a lentiviral sgRNA library (four sgRNAs per gene). Cells were treated with DMSO (0.25%) or CBK77 25 µM thrice for a total of five days. The resulting cell populations were harvested and subjected to barcode sequencing and analysis. Two technical replicates were performed. **B)** Concentration-response experiments performed with MelJuSo Ub-YFP cells. Cell viability was assessed after 72 hours. Data are represented as mean ± SD of three independent experiments. Non-linear curve fitting is depicted in red. The half-maximal inhibitory concentration (IC_50_) and slope of the curve are used to calculate the theoretical IC_90_ concentration. **C)** Scatter plot of the average gene log-fold change (LFC) in CBK77-treated cells versus control (CTRL)-treated cells in the two replicates as calculated using MaGeCK^54^. Each dot represents a gene and is colored from lower (blue) to higher (red) LFC values. The names of genes with an average LFC > 1.5 are shown. **D)** MelJuSo Ub-YFP cells were depleted of NQO1 for 48 hours with the indicated siRNAs (10 nM) and then treated for 24 hours with CBK77 (5 µM). Non-targeting siRNA (siCTRL) was used as control. After the treatment, nuclei were counterstained with Hoechst and cells imaged live with a widefield automated microscope. Frequency and distribution of the nuclear YFP intensity per cell are shown as violin plots from one experiment. Black lines within each distribution represent the median; white lines represent the upper and lower interquartile range limits. The knockdown efficiencies are shown in the blot below. **E)** MelJuSo Ub-YFP cells were co-treated with either DMSO, CBK77, CBK07 (10 µM) or bortezomib (proteasome inhibitor = PI, 25 nM) with or without the NQO1 inhibitor ES936 (100 nM) for 6 hours. Cell lysates were analyzed by immunoblotting with the indicated antibodies. **F)** MelJuSo Ub-YFP cells were treated with CBK77 (10 µM) alone or in combination with the NQO1 inhibitor ES936 at the indicated concentrations for 24 hours. After the treatment, cells were fixed and nuclei counterstained with Hoechst. Cells were imaged with an automated widefield microscope. The percentage of YFP-positive cells are shown as mean of three independent experiments. Error bars depict the SEM. ns non-significant, * adjusted p ≤ 0.05, ** adjusted p ≤ 0.01, **** adjusted p ≤ 0.0001 (one-way ANOVA with Dunnett’s multiple comparisons test). **G)** The cell count from (**F**) was obtained based on the nuclei staining. Data are shown as mean of three independent experiments. Error bars depict the SEM. ns non-significant, * adjusted p ≤ 0.05, ** adjusted p ≤ 0.01, **** adjusted p ≤ 0.0001 (one-way ANOVA with Dunnett’s multiple comparisons test).

The enrichment of these candidates suggested that the presence of NQO1 may be critical for the mode of action of CBK77. In line with this hypothesis, we found that cellular depletion of NQO1 using siRNAs reduced the accumulation of the reporter substrate Ub-YFP in response to CBK77 treatment (**Fig. 5D**). Notably, the rescue of UPS functionality correlated with the level of NQO1 depletion obtained by four independent siRNAs (**Fig. 5D**). To determine if the catalytic activity of NQO1 was required for the inhibitory effect of CBK77, we took advantage of ES936, a potent irreversible inhibitor of NQO1 activity^28^. Consistent with a role for the catalytic activity of NQO1, ES936 abrogated the CBK77-induced accumulation of Ub-YFP, p53 and Hsp70 (**Fig. 5E**). Moreover, administration of ES936 caused a concentration-dependent decrease in CBK77-induced UPS impairment (**Fig. 5F**) and improved cell survival (**Fig. 5G**). An inverse correlation between the level of UPS activity and cell death was evident from these experiments, supporting the model that UPS impairment and cell viability are mechanistically linked. It has been reported that some 5-nitrofurans can cause a general cellular toxicity upon reduction by aldehyde dehydrogenase (ALDH)-2 activity^29^. However, inhibition of ALDH-2 did not alleviate the effects of CBK77, indicating that the UPS inhibition and toxicity caused by CBK77 differs mechanistically from the generic toxicity caused by other 5-nitrofurans (**Suppl. Fig. 3A,B**).

### CBK77 is an NQO1 substrate

Even though the natural substrates of NQO1 are quinones, NQO1 can also reduce certain nitroaromatic compounds^30, 31^. To test if CBK77 is metabolized by NQO1, we performed an *in vitro* enzymatic assay that measures NAD(P)H consumption by NQO1 when reducing substrates. Strikingly, CBK77 induced the oxidation of NAD(P)H comparably to the genuine quinone substrate menadione, which suggests that CBK77 is efficiently metabolized by NQO1 (**Fig. 6A,B**). Accordingly, the enzymatic reaction towards CBK77 and menadione was blocked by the NQO1 inhibitor dicoumarol. On the contrary, NQO1 showed very limited enzymatic activity towards the inactive CBK07 (**Fig. 6A,B**).

**Figure 6.**
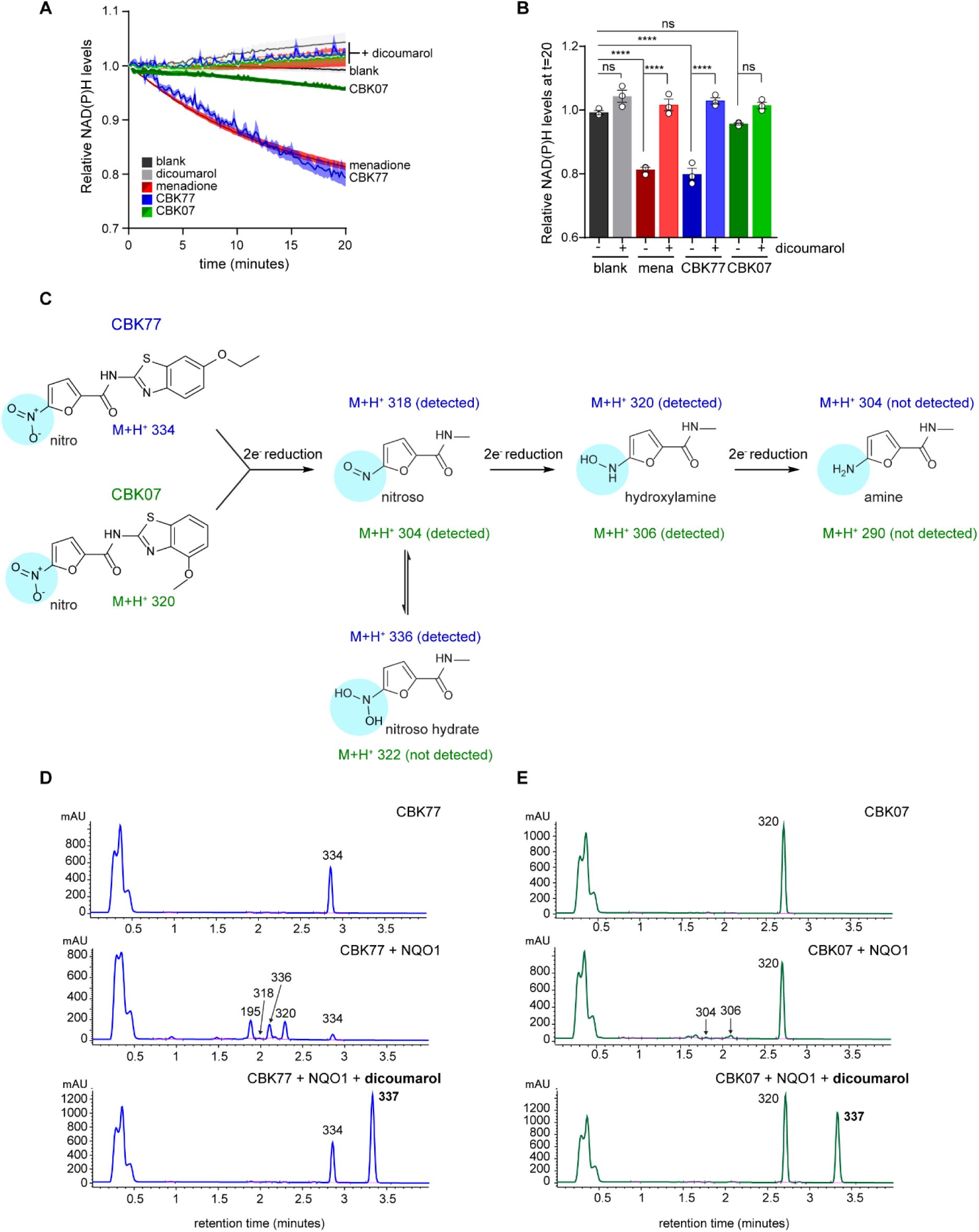
CBK77 is a substrate of NQO1. **A)** Recombinant NQO1 was incubated with CBK77 or CBK07 (100 µM) in the presence or absence of the NQO1 inhibitor dicoumarol (100 µM). The NQO1 substrate menadione (40 μM) with or without dicoumarol was used as control. Plot showing the NAD(P)H levels over time, which are reduced due to the conversion of NAD(P)H to NADP+ in the presence of NQO1 substrates. Data are shown as average NAD(P)H levels ± SEM (shadowed area) of three independent experiments. **B)** Bar plot of the values displayed in the last timepoint (t = 20 minutes) in (**H**). Datapoints are shown as aligned dots ± SEM. ns non-significant, * adjusted p ≤ 0.05, ** adjusted p ≤ 0.01, **** adjusted p ≤ 0.0001 (one-way ANOVA with Tukey’s multiple comparisons test). **C)** Schematic drawing depicting sequential two-electron reductions of the nitro group present in CBK77 and CBK07. These expected sequential reductions produce amines via nitroso and hydroxylamine intermediates. The impact of these reductions on the expected M+H^+^ after electrospray ionization (ESI+) of the compounds are shown below each structure in a color-coded manner (blue = related to CBK77; green = related to CBK07). **D)** HPLC-MS DAD (305 ± 90 nm) chromatograms of *in vitro* reactions incubating CBK77 (100 µM) with or without recombinant NQO1. A sample with dicoumarol (100 µM) was included to assess dependency on NQO1’s activity. The M+H^+^ of expected metabolites found after electrospray ionization (ESI+) in the MSD1 report are annotated above each peak. **E)** HPLC-MS DAD (305 ± 90 nm) chromatograms of *in vitro* reactions incubating CBK07 (100 µM) with or without recombinant NQO1. A sample with dicoumarol (100 µM) was included to assess dependency on NQO1’s activity. The M+H^+^ of expected metabolites found after electrospray ionization (ESI+) in the MSD1 report are annotated above each peak.

The 2-electron reduction of nitroaromatic compounds, which has been previously described^32^, is expected to proceed stepwise involving the formation of three major species: the nitroso, hydroxylamine and amine derivates (**Fig. 6C**). Analytical HPLC-MS confirmed that the level of CBK77 (M+H^+^ 334) in the *in vitro* reaction rapidly decreased in the presence of NQO1 (**Fig. 6D**). The rapid decline in CBK77 was accompanied by the appearance of the 5-nitrosamine derivative (M+H^+^ 318) and the hydroxylamine derivative (M+H^+^ 320), corresponding with two- and four-electron reductions of the parental compound, respectively. Other species detected corresponded with the hydrate form of the 5-nitrosamine derivative (M+H^+^ 336) and the 6-ethoxy-1,3-benzothiazol-2-amine (M+H^+^ 195), which appears to be the product from hydrolysing the amide of the parent compound (**Fig. 6D**). Consistent with the enzymatic assay, CBK07 was poorly metabolized, giving rise to trace amounts of the 5-nitrosamine derivative (M+H^+^ 304) and hydroxylamine derivative (M+H^+^ 306) (**Fig. 6E**). Importantly, the appearance of the CBK77 and CBK07 derivatives could be blocked by the addition of the NQO1 inhibitor dicoumarol (**Fig. 6D,E**). Overall, these data confirm that CBK77, in contrast to CBK07, is efficiently metabolized by NQO1 resulting in the production of metabolites that are in line with the anticipated stepwise two-electron reductions of the parental compound.

### CBK77 interacts with ubiquitin in an NQO1-dependent fashion

To identify molecular targets of CBK77, we developed an activity-based probe that allowed labelling of interacting molecules^33^. For this purpose, we took advantage of our SAR analysis that showed that modifications in the 6 position of the 1,3-benzothiazol group did not affect the activity of CBK77. By introducing an alkyne group at this position, we generated the probe CBK77^CLICK^ for activity-based target identification (**Fig. 7A**). We confirmed that the introduction of the terminal alkyne group did not affect CBK77 activity, demonstrated by a similar EC_50_ in MelJuSo Ub-YFP cells (**Suppl. Fig. 4A**) and comparable kinetics of UPS impairment to CBK77 (**Suppl. Fig. 4B**). Thus, CBK77^CLICK^ was cell permeable and retained the UPS inhibitory activity of the parental compound, which encouraged us to use this probe for target identification. The introduced alkyne moiety allows covalent linkage of a tag-of-choice via copper(I)-catalyzed azide alkyne cycloaddition (CuAAC)^34^. For identification of the CBK77-interactome, we prepared cell lysates from vehicle or CBK77^CLICK^-treated cells and introduced the trifunctional linker (TFL) TAMRA-azide-biotin, which allows fluorescent visualization by the TAMRA fluorophore and biochemical enrichment through the biotin group (**Fig. 7B**). In-gel detection of streptavidin-bead eluates confirmed efficient enrichment of CBK77^CLICK^ conjugates (**Suppl. Fig. 4C**). Mass spectrometry-based analysis of three independent pulldown experiments identified 40 proteins to be enriched in the CBK77^CLICK^-treated samples (**Fig. 7C**). Gene ontology enrichment analysis showed that most of these gene products were involved in cytoskeletal protein binding, nucleobase-containing compound kinases and, interestingly, several proteins were associated with oxidoreductase activities, recapitulating the involvement of a redox-related toxicity mechanism as found in the CRISPR/Cas9 screen (**Suppl. Table S4**). NQO1 was identified in the mass spectrometry experiment but was not significantly enriched in CBK77^CLICK^-treated cells, consistent with the notion that this enzyme is important for processing of the compound but not a primary target of activated CBK77. Most notably, ubiquitin (UBC) was identified as one of the most enriched proteins in the pulldown with the CBK77^CLICK^ probe (**Fig. 7C**).

**Figure 7.**
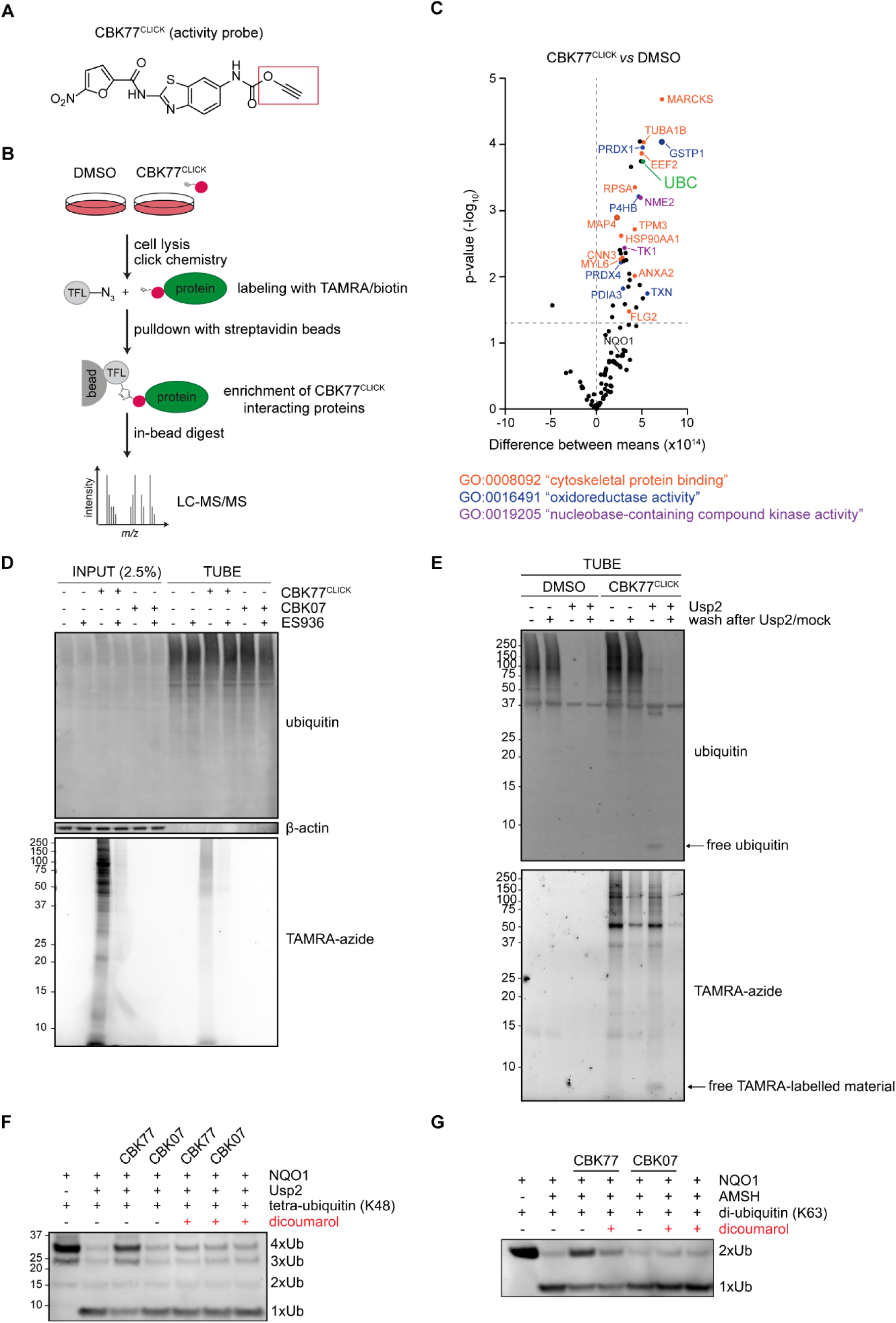
CBK77 interacts with ubiquitin in an NQO1-dependent fashion. **A)** Chemical structure of activity probe CBK77^CLICK^. Boxed in red is the alkyne moiety, which can be used for covalently linking tags via *click* chemistry. **B)** Schematic drawing representing the chemical proteomics workflow. **C)** Volcano plot depicting the results of multiple t-testing in the protein abundance in CBK77^CLICK^-treated cells versus DMSO. Abundance is defined by the average peak area of the three most intense peptides, including unique and non-unique peptides. The x-axis represents the difference between means; the y-axis represents the statistical significance as the -log_10_ p-value. Dashed lines indicate the threshold for significance (p-value ≤0.05) and the point with no difference between the groups (x=0). No correction for multiple comparisons was carried out to have a less stringent, hypothesis-generating approach. The names of the proteins involved in significantly enriched molecular functions are colored according to the legend. Ubiquitin (UBC) was the only enriched protein with a direct role in the UPS. **D)** MelJuSo cells were treated with either DMSO, CBK77, CBK77^CLICK^ (100 µM) or CBK77^CLICK^ in combination with the NQO1 inhibitor ES936 (100 nM) for 45 minutes. After cell lysis and labelling of the compound with a TAMRA-azide fluorophore, ubiquitin was pulled down with TUBE-agarose beads. Binding of the compound to ubiquitin was assessed via TAMRA detection and immunoblotting with a ubiquitin antibody. **E)** MelJuSo Ub-YFP cells were treated with CBK77^CLICK^ (10 µM) for 1 hour. After cell lysis and labelling of the compound with a TAMRA-azide fluorophore, ubiquitin was pulled down with TUBE-agarose beads. After incubation, the beads were treated with or without Usp2 for 1 hour. A set of samples were stopped directly with addition of LDS-sample buffer, while a second set was further washed to remove unbound proteins. Binding of the compound to ubiquitin was assessed via TAMRA detection and immunoblotting with a ubiquitin antibody. **F)** Recombinant NQO1 was incubated with CBK77 or CBK07 (100 µM) in the presence or absence of the NQO1 inhibitor dicoumarol (100 µM) in the presence of K48-linked tetraubiquitin chains. Samples were then treated with the recombinant catalytic domain of the DUB Usp2 and cleavage of the ubiquitin moieties was assessed via SDS-PAGE and immunoblotting with a ubiquitin antibody. **G)** Recombinant NQO1 was incubated with CBK77 or CBK07 (100 µM) with K63-linked di-ubiquitin chains in the presence or absence of the NQO1 inhibitor dicoumarol (100 µM). Samples were then treated with 233 nM of recombinant AMSH and cleavage of the ubiquitin moieties was assessed via SDS-PAGE and immunoblotting with a ubiquitin antibody.

In an siRNA-mediated depletion of the proteins that were enriched in the CBK77 interactome, we did not observe accumulation of Ub-YFP for any of the candidates (**Suppl. Fig. 4D**). The effect of ubiquitin depletion on Ub-YFP degradation could, however, not be tested as the ubiquitin-targeting siRNA would also reduce the levels of Ub-YFP due to its N-terminal ubiquitin moiety. Therefore, we turned to the YFP-CL1 reporter cell line and observed, as expected, that depletion of ubiquitin resulted in stabilization of YFP-CL1 (**Suppl. Fig. 4E**). Thus, the only CBK77-interacting protein that upon depletion phenocopied CBK77-induced UPS impairment was ubiquitin itself. To validate the covalent modification of endogenous ubiquitin by CBK77, we performed ubiquitin pulldowns with tandem-repeated ubiquitin-binding entities (TUBEs), which are recombinant proteins composed of naturally occurring ubiquitin binding domains^35^. A specific smear of high-molecular weight CBK77^CLICK^-labelled proteins was detected that was reminiscent to the signal observed for endogenous ubiquitin conjugates (**Fig. 7D**). The NQO1 inhibitor ES936 strongly reduced the TAMRA labelling (**Fig. 7D, Suppl. Fig. 4F**), suggesting that binding to ubiquitin conjugates, like UPS impairment and cell death, is dependent on NQO1 activity.

The labelling of ubiquitin conjugates on the TUBEs, as well as the enrichment of ubiquitin in the CBK77 interactome, could be due to covalent binding of CBK77 to ubiquitylated substrate(s) or, alternatively, to ubiquitin itself. To address this question, we performed again enrichment of ubiquitin conjugates with TUBEs, but this time we treated the enriched fractions with the recombinant catalytic domain of ubiquitin specific protease 2 (Usp2). Usp2 is a protease that specifically disassembles ubiquitin chains of various linkages. We reasoned that if CBK77^CLICK^ binds to ubiquitin directly, treatment with Usp2 should lead to TAMRA-labelled ubiquitin monomers. Our results are in favour of direct binding of CBK77^CLICK^ to ubiquitin moieties, as treatment with Usp2 resulted in the appearance of a TAMRA-labelled ubiquitin monomers (**Fig. 7E**). Notably, while the appearance of the labelled ubiquitin monomers occurred at the expense of high-molecular weight species labelled with TAMRA, consistent with the mono-ubiquitin being derived from disassembled polyubiquitin chains, a substantial signal of the TAMRA-labelled smear remained. We argued that this could be due to CBK77^CLICK^ interacting with deubiquitylated substrates and/or CBK77^CLICK^-modified ubiquitin conjugates that had not been disassembled by Usp2. Washing of the beads after Usp2 treatment resulted in the disappearance of the ubiquitin monomers, in line with the specificity of the TUBEs for polyubiquitin chains, while some of the TAMRA-labelled smear remained, which may suggest that the remaining high molecular weight signal is, at least in part, due to TAMRA-labelled polyubiquitylated substrates that were not deconjugated by Usp2. We conclude that NQO1-activated CBK77 modifies polyubiquitin chains.

### CBK77-modified ubiquitin conjugates resist deubiquitylating enzymes

Even though CBK77 does not affect overall DUB activity, it may nevertheless inhibit deubiquitylation by rendering the ubiquitin chains less susceptible to DUB activity. To directly address this question, we performed an *in vitro* experiment in which Usp2 was incubated with K48-linked tetraubiquitin chains after the chains had been exposed to NQO1-activated CBK77. Inhibition of the disassembly of K48-linked tetraubiquitin chains by Usp2 was observed when tetraubiquitin had been incubated with activated CBK77, which could be prevented by blocking NQO1 activity with dicoumarol (**Fig. 7F**). To assess whether this reflects a more general phenomenon, we performed a similar assay with the deubiquitylating enzyme AMSH, a metalloprotease representing a different class of deubiquitylating enzymes with a preference for K63-linked ubiquitin chains^36^. This revealed a similar inhibitory activity of CBK77 towards disassembly of these ubiquitin chains (**Fig. 7G**). Our data show that CBK77-modified ubiquitin chains are less susceptible to deubiquitylating activities *in vitro*, providing a possible explanation for the observed accumulation of polyubiquitylated proteins and selective impairment of ubiquitin-dependent proteasomal degradation in cells exposed to CBK77.

### CBK77 reduces growth of NQO1-proficient human cancer cells and xenograft tumors in mice

To further explore the importance of NQO1 for potential anti-tumor activity of CBK77, we took advantage of a naturally occurring NQO1 polymorphism. Individuals that express the unstable NQO1^C609T^ variant have undetectable or trace levels of the NQO1 protein and therefore very low NQO1 activity^37^. As expected, a breast cancer cell line derived from an individual expressing this NQO1 variant (MDA-MB-231) were largely resistant to CBK77 with an IC_50_ of > 50 µM, which was very similar to the sensitivity of the cell line to the poorly active CBK07 analogue (**Fig. 8A**). However, ectopic expression of stable NQO1 in MDA-MB-231 cells resulted in a 10-fold increased sensitivity to CBK77, while having little effect on CBK07 sensitivity (**Fig. 8B**). In line with this toxicity profile, we observed that ectopic expression of stable NQO1 in MDA-MB-231 cells resulted in the accumulation of ubiquitin conjugates and an increase in Hsp70 levels, indicative of a global UPS impairment (**Fig. 8C**). The effects of ectopic NQO1 expression were abrogated by administration of the NQO1 inhibitor ES936, showing that the proposed anti-cancer effect of CBK77 is dependent on NQO1 activity.

**Figure 8.**
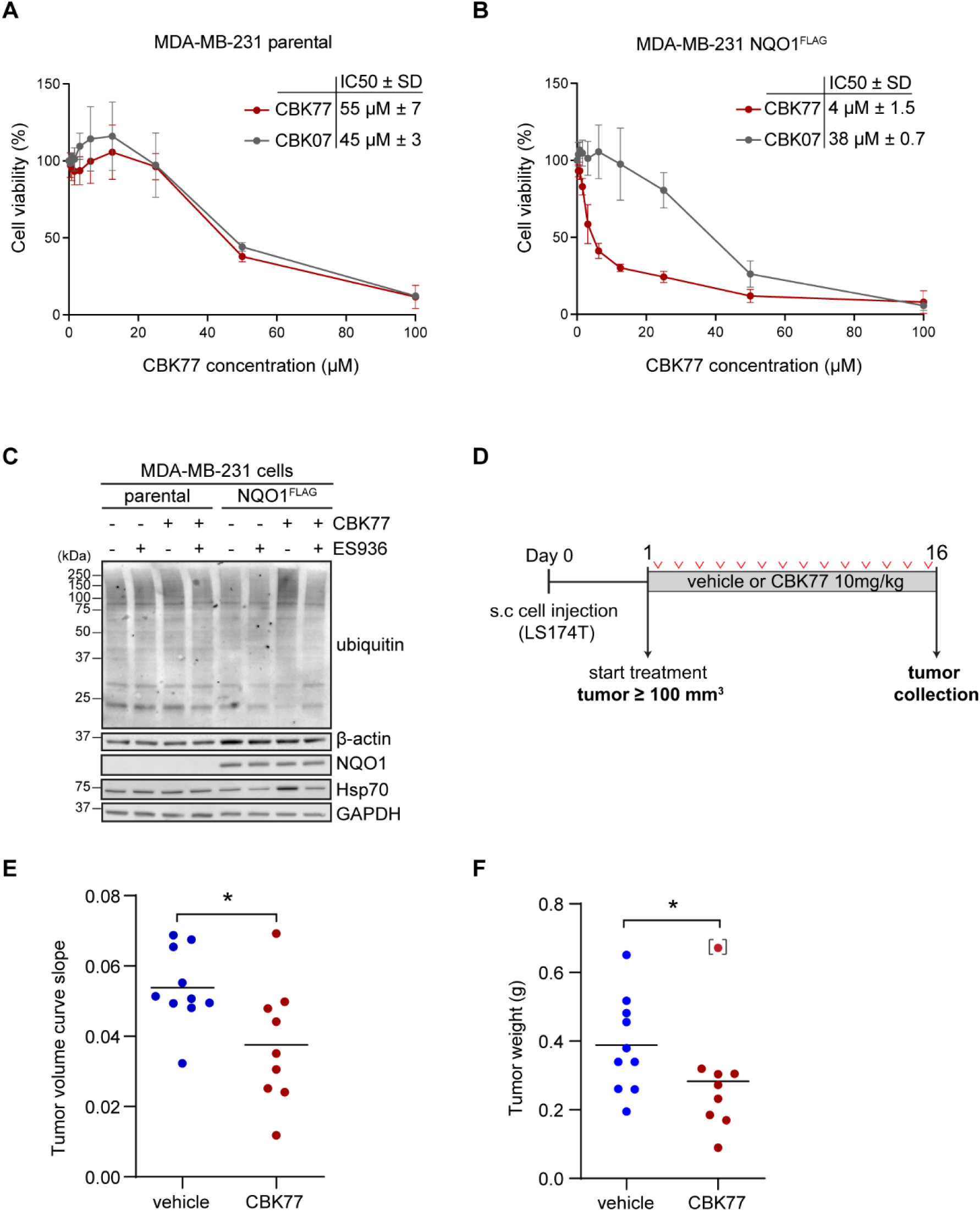
CBK77 reduces growth of NQO1-proficient human cancer cells and xenograft tumors in mice. **A)** Concentration-response experiments performed with MDA-MB-231 parental cells. Cell viability was assessed after 72 hours using the WST-1 proliferation reagent. Data are represented as mean ± SEM of three independent experiments. The half-maximal inhibitory concentrations (IC_50_) ± SD are shown. **B)** Concentration-response experiments performed with MDA-MB-231 cells stably expressing NQO1^FLAG^. Cell viability was assessed after 72 hours using the WST-1 proliferation reagent. Data are represented as mean ± SEM of three independent experiments. The half-maximal inhibitory concentrations (IC_50_) ± SD are shown. **C)** MDA-MB-231 parental and NQO1^FLAG^-expressing cells were incubated with the indicated compounds for 6 hours. Lysates were probed with the indicated antibodies. **D)** Experimental timeline for the xenotransplant experiment. **E)** Scatter plot of the tumor volume curve slopes obtained in a simple linear regression model (according to ^55^). * p-value = 0.024 (two-sided, unpaired t-test). **F)** Scatter plot of the tumor weights (grams) at necropsy (day 16). The median value is depicted as a solid black line. A Grubbs’ outlier test identified one tumor as a significant outlier in the CBK77 group (marked in brackets). Note that all tumors in the treated group, besides the one identified as an outlier, are clustered below the median of the vehicle group. * p-value = 0.014 (two-sided, unpaired t-test, excluding outlier datapoint).

For an *in vivo* assessment of the efficacy of CBK77 against tumors, we selected the human colon adenocarcinoma cell line LS174T, which expresses functional NQO1 at a comparable level to the MelJuSo cell line used in the high-content screen (**Suppl. Fig. 5A**). Accordingly, LS174T cells were found to be sensitive to CBK77 (**Suppl. Fig. 5B**). LS174T cells were injected subcutaneously into the flanks of immunocompromised NMRI nu/nu mice. When the tumors had reached a minimum size of 0.1 mL, mice were randomly assigned to either daily intraperitoneal administration of vehicle (solvent only) or 10 mg/kg CBK77 (**Fig. 8D**). During treatment, we did not observe weight loss or obvious behavioural signs of toxicity (**Suppl. Fig. 5C**). Consistent with its *in vitro* activity, we found that CBK77 administration resulted in a significant reduction in tumor growth rates (**Fig. 8E**) and tumor weights at necropsy (**Fig. 8F**). These data show that CBK77 inhibits growth of an NQO1-proficient cell line in a xenotransplant tumor model without overt signs of toxicity.

## Discussion

Despite its richness in potentially druggable molecular targets, the few clinically approved drugs targeting the UPS are all directed against the proteolytic activity of the proteasome^8^. Our high-content phenotypic screen resulted in the discovery of a compound that effectively blocked proteasomal degradation by an unanticipated mechanism that differs from the mode of action of proteasome inhibitors. Our study revealed that CBK77 does not interfere with global ubiquitylation, deubiquitylation or proteasome activity. Moreover, its inhibitory activity is confined to ubiquitin-dependent degradation and strictly depends on NQO1. To the best of our knowledge, CBK77 is the first UPS inhibitor that relies for its inhibitory activity on activation by an endogenous enzyme. While the first non-peptidic proteasome inhibitor, the natural product lactacystin, also behaves as a prodrug, its active component β-lactone is generated by a spontaneous chemical reaction (i.e. lactonization) without involvement of cellular enzymes^38^.

An attractive strategy for enhancing tumor specificity is the use of prodrugs that are activated in the altered intracellular environment of cancer cells. It has been previously proposed that NQO1 can be exploited for this purpose since it is upregulated in many solid tumors as part of the anti-oxidative response governed by the transcription factor Nrf2^39^. It has been shown that the elevated levels of this oxidoreductase can provide a mean to restrict the effect of redox-activated prodrugs primarily to cancer cells^40^. A notable example of an NQO1-activatable anti-tumorigenic drug is the naturally occurring quinone Mitomycin C, which upon reduction by NQO1 behaves as a DNA alkylating agent^41^. While quinones are its primary natural substrates, NQO1 can also reduce certain nitroaromatic compounds as we confirmed for CBK77, which was found to be an efficient substrate on par with the classic NQO1 substrate quinone menadione.

A few studies have linked NQO1 to processes that involve the UPS. *In vitro* data showed that apo-NQO1 – i.e. in the absence of its coenzyme FAD – interacts with the 20S proteasome and is degraded in a ubiquitin-independent fashion, which has been suggested to play a regulatory role in controlling levels of active NQO1 complex in cells^42^. Moreover, NQO1 has also been shown to inhibit proteasomal degradation of ODC^43^, p53^44^ and HIF1α^45^. Given the specific nature of this NQO1-mediated stabilization and the fact that, at least for ODC and HIF1α, stabilization depends on a direct interaction of NQO1 with intrinsically disordered structures in these proteins^43, 45^, we consider it unlikely that the general UPS impairment caused by CBK77 is mechanistically linked to this specific stabilizing activity of NQO1. However, the tight link between NQO1 and the UPS and its ability to interact with the proteasome may be of relevance for the inhibitory effect of CBK77 as this may position the presumably short-lived, reactive CBK77 metabolites in close proximity to key players of the UPS, such as ubiquitin.

Our data suggest that CBK77 targets a protein centrally involved in ubiquitin-dependent proteasomal degradation. In line with this notion, we identified ubiquitin as a protein that is covalently modified by the activity probe. Moreover, CBK77 impaired the cleavage of ubiquitin chains *in vitro* of DUB enzymes of different families, suggesting a broad effect that would be difficult to explain by inhibition of the catalytic activity of these enzymes. The irreversible collapse of the UPS in the absence of global inhibition of ubiquitylation, DUB or proteasomal activity support a model in which impaired or inefficient disassembly of polyubiquitin chains leads to an accumulation of UPS substrates that ultimately causes cell death. Blocking of the UPS by ubiquitin-interacting compounds is not unprecedented. Ubistatins are a group of large, symmetrically organized molecules that non-covalently interact with two ubiquitin monomers simultaneously, thereby interfering with the targeting of polyubiquitylated proteins to the proteasome^13^. Future work focusing on the interaction between CBK77 and ubiquitin conjugates, and to what extent CBK77-modified ubiquitin conjugates contribute to UPS impairment and cell death, will be required to clarify CBK77’s mechanism of action.

Although CBK77 passed PAINS filters^46^, it raised some medicinal chemistry concerns as the structural alert moiety nitrofuran was found to be critical for its biological effect. It should be noted, however, that the 5-nitrofuran compound Nitrofurantoin has been used for more than half of a century for treatment of urinary tract infection with relatively mild adverse effects^47^. Moreover, another nitrofuran-based compound, Nifurtimox, is used as a trypanocidal drug for treatment of Chagas disease^48^. In our testing of CBK77 in a xenotransplant mouse model, we did not observe acute signs of toxicity caused by CBK77 while the tumor growth was significantly reduced, which further illustrates its potential as a lead compound. It is noteworthy that in an earlier high-throughput screen, related nitrofuran compounds were found to activate the heat-shock response by an unknown mechanism^49^. Our data provide an explanation for this earlier observation, as it is well established that induction of the heat-shock response can be a direct consequence of a disturbed protein homeostasis^17^.

In summary, we have identified and characterized a novel, bioactivatable inhibitor of the UPS that causes an irreversible collapse of the UPS, induces caspase-dependent cell death and displays anti-tumor activity. Our data show that CBK77 blocks the UPS in an NQO1-dependent fashion. The fact that NQO1 is often upregulated in solid tumors opens possibilities in treating tumors with poor sensitivity towards proteasome inhibitors. We propose that CBK77 can provide a starting point for the development of a novel class of UPS inhibitors.

## Acknowledgements

We thank Jelena Strenger, Simona Walker, Anne Beskow and Kristian Björk Grimberg for performing experiments in the early phase of the project. We also thank Galina Selivanova, Marcela Franco, Janne Lethiö, Sonia Laín, Elias Arnér, Stig Linder and Andreas Fälting and Mauricio Barrientos-Somarribas for help and critical feedback. Part of this work was carried out in the High Throughput Genome Engineering Facility (HTGE) and the Clinical Proteomics facility, both funded by Science for Life Laboratory. We acknowledge support from the National Genomics Infrastructure. Computations were performed on resources provided by SNIC (project SNIC 2017-7-265) on the Uppsala Multidisciplinary Center for Advanced Computational Science (UPPMAX). We also acknowledge support from SciLifeLab’s Drug Discovery and Development (DDD) platform, particularly from André Mateus and Richard Svensson from the UDOPP facility in Uppsala. This work was supported by the Swedish Research Council (N.P.D. 2016-02479; R.J. 2017-01653), the Swedish Cancer Society (N.P.D. CAN 2018/693; R.J. 174182) and the Karolinska Institute.

## Author contributions

FAS, PY performed the initial screen; MH, TL, ALG assisted with screen logistics, hit follow-up and designed and synthesized the compounds; TAG, FAS characterized the compound and its mode of action; TAG, LHME, MW, JIJ tested the compound in mice; TAG, MA performed the DUB assays and analyzed the data; TAG, RJ, JE, performed the mass spectrometry and data analysis; TAG, FAS, MH, NPD wrote the draft of the manuscript; TAG prepared the figures; FAS, NPD coordinated the project;. All authors edited and approved the final manuscript.

## Materials & Methods

### Compound screen

Details of the primary compound screen can be found in **Supplementary Table S1**.

### Cell culture

The human melanoma MelJuSo cells, human cervical cancer HeLa and the human breast carcinoma MDA-MB-231 cells were maintained in DMEM. LS174T cells were maintained in RPMI 1640 medium. All media contained high glucose, GlutaMAX and sodium pyruvate, and were supplemented with 10% fetal bovine serum. All cells were maintained at 37°C in a humidified 5% CO_2_ atmosphere, and tested periodically for mycoplasma infection by PCR. MelJuSo cells stably expressing Ub-YFP, Ub-R-GFP, YFP-CL1 and CD3δ-YFP were previously generated^50^. The ZsGreen-ODC (ornithine decarboxylase) cells were previously generated by transfection of MelJuSo cells with the ZsProsensor-1 plasmid (BD Bioscience Clontech)^20^. MDA-MB-231 cells stably expressing NQO1^FLAG^ were generated by transfection of MDA-MB-231 cells with the NQO1-FLAG plasmid. After 16 hours, selection was started (1 mg/ml geneticin). Clones expressing NQO1^FLAG^ were isolated and validated by western blotting. For the CRISPR/Cas9 screen, the MelJuSo cell line was made to stably express the Cas9 nuclease. In brief, a construct coding for Cas9 and blasticidin under the control of the EF1α promoter was introduced by lentiviral transduction. After two weeks of blasticidin selection, Cas9 expression was confirmed by western blot. Plasmid DNA and siRNA transfections were performed using lipofectamine 2000 (Thermo Fisher Scientific) according to the manufacturer’s instructions.

### Cell proliferation assay

Cells were seeded into 96-well plates at 1500-5000 cells/well. Sixteen hours after seeding (ca 60% confluency), cells were treated with the indicated compound concentrations in a serial dilution. DMSO at 1% was used as control. After 48 or 72 hours of incubation, WST-1 tetrazolium salt (Roche) was added and incubated for 1 hour at 37°C. The formation of formazan was assessed by measuring the absorbance at 480 nm with the plate reader FLUOStar OPTIMA (BMG Labtech).

### Western blotting

Equal amounts of cells were lysed in 1X SDS sample buffer (Tris-HCl 0.3 M pH 6.8, 2% SDS, 17.5% glycerol, bromophenol blue) containing 10% NuPAGE reducing agent (Thermo Fisher Scientific) and lysates were boiled at 95°C for 5 minutes. Cell protein extracts were resolved by Bis-Tris polyacrylamide gel electrophoresis gels (Thermo Fisher Scientific) and run in either MOPS or MES buffer (Thermo Fisher Scientific). Proteins were transferred onto PVDF 0.45 µm or nitrocellulose membranes (GE Healthcare) in a Tris-glycine transfer buffer containing 20% methanol. After blocking in Tris-buffered saline (TBS)/non-fat milk 5% containing 0.1% Tween-20, membranes were incubated with primary antibodies, washed with TBS-Tween 20 0.1% and incubated with secondary HRP-linked antibodies. Detection was performed by enhanced chemiluminiscence (Amersham ECL reagents, GE Healthcare) on X-ray films (Fujifilm). Alternatively, secondary antibodies coupled to near-infrared fluorescent dyes (LI-COR) were used, and membranes scanned with an Odyssey scanner (LI-COR) and analyzed with Image Studio Lite analysis software version 5.2 (LI-COR).

### CRISPR/Cas9 interference screen

#### Brunello-UMI Library

The genome-wide Brunello sgRNA library^51^ was resynthesized to include Unique Molecular Identifiers^52^. Guides were cloned in pool and packaged into lentivirus. Lentiviral backbone was based on lentiGuide-Puro (Addgene # 52963), with AU-flip as in^53^.

#### Screen

Cas9-expressing cells were transduced with the Brunello-UMI library at an approximate MOI of 0.4 and 1000 cells/guide in 2 µg/ml polybrene. Transduced cells were selected with puromycin (2 µg/ml) from post-transduction day two to seven and then split into two replicates cultured independently until 16 days after transduction. Cells were treated with either DMSO or CBK77 at 25 µM. Two replicates of CBK77-treated cells were performed in parallel with a common DMSO-treated sample (the number of CBK77-treated cells and DMSO-treated cells corresponded to approximately 200-fold and 500-fold library coverage, respectively). When less than 10% of the cells were left, cells were harvested and genomic DNA was isolated using the QIAamp DNA Blood Maxi kit (Qiagen), and guide and UMI sequences were amplified by PCR. Sequencing was performed on an Illumina HiSeq instrument. Next Generation Sequencing (NGS) data were analyzed with MaGeCK^54^ and by UMI lineage dropout analysis^52^. We considered interesting genes those which guides had an average log fold change (LFC) higher than 1 and had a false discovery rate (FDR) lower than 0.1. A summary of the results can be found in **Supplementary Table S3**.

### Protein purification

BL21 *E. coli* bacteria were transformed with the pGEX-6P-1 plasmid encoding GST-NQO1. Overnight cultures of individual colonies were grown at 37°C in the presence of 100 μg/ml ampicillin. Bacteria were grown at 37°C in 500 ml LB until reaching OD600 = 0.6, when protein expression was induced by 1 mM IPTG for 3.5 hours at 30°C. Bacteria were harvested and resuspended in GST lysis buffer (50 mM HEPES pH 7.5, 500 mM NaCl, 10% glycerol, 1 mM MgCl2, 10 mM DTT) and lysed by sonication. The lysate was cleared and incubated with equilibrated glutathione sepharose 4B beads (GE Healthcare) overnight at 4°C. Beads were washed in GST lysis buffer and GST-NQO1 was eluted for 4 hours at 4°C in elution buffer (50 mM Tris pH 7.5, 10 mM reduced glutathione, 10 mM DTT). To obtain untagged NQO1, beads were washed in PreScission cleavage buffer (50 mM Tris pH 7.5, 150 mM NaCl, 1 mM EDTA, 1 mM DTT) and protein was eluted by ca 40 units PreScission protease (GE Healthcare) for 4 hours at 4°C under rotation. The purity of the resulting protein was analyzed on a Coomassie gel.

### NQO1 *in vitro* assay

NQO1 activity was assessed by measuring the levels of NAD(P)H via absorbance at 340 nm in a Versamax plate reader (Molecular Devices) at 37°C. The reaction mixture in a final volume of 200 µl contained 50 mM ammonium bicarbonate, 0.1 mg/ml BSA, 2 mM EDTA, 200 µM NAD(P)H as electron donor (N7505, Sigma-Aldrich) and 2 µg of recombinant NQO1 (for final enzyme concentration of ca 0.5 µM). Positive and negative controls were included by adding 400 µM of the NQO1 known substrate menadione (M5625, Sigma-Aldrich) in the presence or absence of the NQO1 inhibitor dicoumarol at 100 µM, respectively. To assess whether the compounds were substrates of NQO1, 100 µM of the indicated compounds were added instead of menadione in the absence or presence of dicoumarol to measure specific NQO1 activity. Measurements were made every 10 seconds for 20 minutes. For analytical high-performance liquid chromatography with diode array detection (DAD) and electrospray ionization mass spectrometry (HPLC-MS), the *in vitro* reactions were filtered using Amicon Ultra-0.5 centrifugal filter devices with a 30kD molecular weight cut-off (Merck-Millipore) for 10 minutes at 13,000× rpm. 100 µl of flow-through samples were injected on an Agilent/HP 1200 system 6110 mass spectrometer with electrospray ionization (ESI+). Absorbance was monitored at 305 ± 90.

### Proteasomal activity

After treatments, cells were trypsinized and lysed (25 mM HEPES pH 7.2, 50 mM NaCl, 1 mM MgCl_2_, 1 mM ATP, 1 mM DTT, 10% glycerol, 1% Triton X-100) for 30 minutes at 4°C. 10 μg protein in 100μl reaction buffer were mixed with 100 μM Suc-Leu-Leu-Val-Tyr-AMC (Affiniti, P802). In one well, 500 nM epoxomicin was added additionally. Samples were analyzed in a microplate reader (FLUOStar OPTIMA, BMG Labtech) at 380 nm/440 nm every minute for 1 hour.

### Cellular DUB activity

After treatments, cells were trypsinized and lysed (25 mM HEPES pH 7.2, 50 mM NaCl, mM MgCl_2_, 1 mM ATP, 1 mM DTT, 10% glycerol, 1% Triton X-100). Protein concentrations were measured with the Bradford assay (Bio-Rad). 10 μg protein were mixed with 80 μl reaction buffer (lysis buffer without Triton X-100) and 10 μl ubiquitin-AMC (Enzo Life Sciences, BML-SE211-0025) for a final concentration of 1 μM. Samples were analyzed in a microplate reader (FLUOStar OPTIMA, BMG Labtech) at 355 nm/460 nm every 30 seconds for 1 hour.

### DUB profiling

Labelling of DUBs with activity-based probe was carried out as previously described^19^. In brief, after treatments, cells were lysed (50 mM Tris buffer pH 7.4, 150 mM NaCl, 5 mM EDTA, 0.5% NP-40). Lysates were incubated on ice for 45 minutes and cleared by centrifugation (10,000× g for 20 minutes at 4°C). Protein concentrations were measured by BCA. 100 μg of protein were incubated with 4 µg of HA-tagged ubiquitin glycine vinyl methylester (HA-UbVME) or HA-tagged ubiquitin bromoethylamine (HA-UbBr2) for 2 hours at 37°C. A DMSO-treated lysate without probe was included as a background control. After incubating, samples were denatured by the addition of LDS-sample buffer and boiled at 95°C for 5 minutes. Samples were analyzed by SDS-PAGE and immunoblotting with anti-HA and specific DUB antibodies.

### *CLICK* chemistry and analytical labelling

Cells were washed with PBS and the pellets lysed in lysis buffer (1% NP-40, 100 mM NaCl, 50 mM HEPES pH 7.2). Following centrifugation (10 min, 16,000× g, 4°C), supernatants were supplemented with 50 µM TAMRA-azide (Jena Bioscience), 5 mM sodium ascorbate (Sigma-Aldrich), 5 mM aminoguanidine-HCl (Sigma-Aldrich), 0.1 mM CuSO4 (Sigma-Aldrich) and 0.5 mM THPTA (Jena Bioscience), with gentle vortexing. The reaction mixtures were incubated at RT for 30 min, light protected. For gel electrophoresis, 4x LDS sample buffer was added. Samples were resolved by SDS-PAGE. In-gel fluorescence was recorded using a ChemiDoc MP Imaging System (Bio-Rad) equipped with 695/55 (Cy5, for visualization of the Precision Plus Protein Standard, Bio-Rad) and 605/50 (Cy3/TAMRA) filters and coupled to Image Lab v 5.0 analysis software (Bio-Rad).

### Streptavidin beads pulldown using the TFL TAMRA-azide-biotin

Cells previously seeded into 10 cm dishes were treated with either CBK77^CLICK^ at 20 µM or with DMSO 0.2% for 1.5 hours. Cells were then lysed in the aforementioned lysis buffer. Following centrifugation (16,000× g, 15 min, 4 °C), supernatants were supplemented with 50 µM trifunctional linker TAMRA-azide-PEG3-biotin (TFL, Jena Bioscience), and *click* chemistry performed as described. Proteins were precipitated by the addition of a 5-fold volume excess of acetone and incubated overnight at −20 °C. Following centrifugation (16,000× g, 30 min, 4 °C), the supernatant was discarded, and pellets washed two times with prechilled methanol. Subsequently, pellets were dissolved in 0.2% SDS in 25 mM sodium bicarbonate by sonication and incubated under gentle mixing with 100 μL of Dynabeads M-280 streptavidin (Sigma-Aldrich) for hours at 4°C. The beads were washed three times with 25 mM ammonium bicarbonate/0.2% SDS, twice with 6 M urea, and three times with 25 mM ammonium bicarbonate. For gel analysis, 15 μL 2x SDS loading buffer were added and the proteins released for SDS-PAGE by 10 min incubation at 95°C.

### Proteomics

The bead-bound proteins after streptavidin-beads pulldown were reduced in 1 mM DTT at 37°C for 1 hour. Next, samples were alkylated by incubation in the dark at RT in 5 mM chloroacetamide for 10 minutes. After, any remaining chloroacetamide was quenched by the addition of 5 mM DTT. Digestion was carried out by the addition of 0.4 µg Trypsin (sequencing grade modified, Pierce) overnight at 37°C. Peptides were collected and dried using a speedvac. Peptides were then dissolved in 30 µl 3% acetonitrile + 0.1% formic acid and transferred to an MS-vial.

#### LC-ESI-MS/MS

LC-MS analysis was performed using a Dionex UltiMate 3000 RSLCnano System coupled to a Q-Exactive mass spectrometer (Thermo Scientific). 5 µl per sample were injected. Samples were trapped on a C_18_ guard desalting column (Acclaim PepMap 100, 75 µm x 2 cm, nanoViper, C_18_, 5 µm, 100 Å), and separated on a 50 cm long C_18_ column (Easy spray PepMap RSLC, C_18_, 2 µm, 100Å, 75 µm x 15 cm). The nano capillary solvent A was 95% water, 5% DMSO, 0.1% formic acid; and solvent B was 5% water, 5% DMSO, 95% acetonitrile, 0.1% formic acid. At a constant flow of 0.25 μl min^−1^, the curved gradient went from 6% B up to 43% B in 180 min, followed by a steep increase to 100% B in 5 min.

FTMS master scans with 60 000 resolution (and mass range 300-1500 m/z) were followed by data-dependent MS/MS (30 000 resolution) on the top 5 ions using higher energy collision dissociation (HCD) at 30% normalized collision energy. Precursors were isolated with a 2 m/z window. Automatic gain control (AGC) targets were 1e6 for MS1 and 1e5 for MS2. Maximum injection times were 100 ms for MS1 and MS2. The entire duty cycle lasted ∼2.5 s. Dynamic exclusion was used with 60 s duration. Precursors with unassigned charge state or charge state 1 were excluded. An underfill ratio of 1% was used.

#### Peptide and protein identification

The MS raw files were searched using Sequest HT and validated with Percolator with Proteome Discoverer 1.4 (Thermo Scientific) against human UniProt database (downloaded 2015-04-06) and filtered to a 1% FDR cut off. We used a precursor ion mass tolerance of 15 ppm, and product ion mass tolerances of 0.02 Da for HCD-FTMS. The algorithm considered tryptic peptides with maximum 2 missed cleavages; carbamidomethylation (C) as fixed modifications; oxidation (M) as variable modifications. The mass spectrometry proteomics data have been deposited to the ProteomeXchange Consortium via the PRIDE partner repository with the dataset identifier **PXD019519**.

### Gene ontology

Enriched molecular functions in the proteomics dataset were calculated using String (https://string-db.org/). The minimum required interaction score was selected to high confidence (0.7) and only ‘Experiment’ and ‘Databases’ were considered. Results are displayed in **Supplementary Table S4**.

### Microscopy

Widefield fluorescent images were taken with the ImageXpress Micro XLS microscope (Molecular Devices) equipped with 10 or 20X air objectives (NA 0.45 Nikon ELWD) and a dichroic mirror set for DAPI, FITC, Cy3, Cy5 and Texas Red. Live-cell imaging experiments for 24 hours were performed in Leibovitz’s L-15 medium (Thermo Fisher Scientific) at 37°C. Imaging was performed using the MetaXpress software (Molecular Devices). For confocal images, the Zeiss LSM880 microscope equipped with a 63x Plan-A (1.4 NA) oil-immersion lens was used. Images were recorded using the ZEN 2012 software. Images were analyzed with either in-built pipelines in the MetaXpress software (Molecular Devices), custom-made pipelines in CellProfiler (www.cellprofiler.org) or ImageJ (https://imagej.nih.gov/ij/). ImageJ was also used for the processing of representative images.

### *In vivo* experiments

Nineteen NMRI nu/nu female mice 6-8 weeks of age (Taconic) were anesthetized and 10 million LS174T cells were subcutaneously injected into a flank of the mouse. When the tumors reached a minimum volume of 0.1 mL, mice were randomly assigned to either vehicle (n= 10) or compound (n= 9) treatment (day 1). The mice were treated daily for a maximum of 16 days with intra peritoneal injection of either vehicle (solvent only) or compound at 10 mg/kg concentration. At day 16, animals were sacrificed, and tumors weighted and cut into two parts, which were fixed in 4% formalin (for standard FFPE processing) or were frozen at – 80°C. Tumor size was assessed every two days using a caliper and tumor volumes were calculated using the formula length × width^2^ × 0.44. For calculating the tumor growth slopes, tumor volumes were turned into logarithmic scale and fitted into a simple linear regression model^55^. Animals were maintained at a maximum of six per cage and given sterile water and food ad libitum. All animal experiments were approved by the Stockholm ethics committee for animal research (no. N231/14), appointed and under the control of the Swedish Board of Agriculture and the Swedish Court. The animal experiments presented herein were in accordance with national regulations (SFS no. 2018:1192 and SFS no. 2019:66).

### Statistical analysis

Statistical analyses were performed using RStudio, Python or GraphPad Prism version 8.3. To test for Gaussian distribution, the D’Agostino & Pearson normality test was used. If the normality test was passed, data were analyzed by Student’s unpaired t-test (two groups) or by ANOVA (more than two groups). If the data were not normally distributed, statistical analysis was performed using the nonparametric Mann-Whitney test (two groups) or Kruskal-Wallis test for multiple comparisons, with Dunnett’s or Tukey’s test to adjust for multiple comparisons. Grubbs’ test was used for the detection of outliers. Adjusted p-values are shown. For the *in vivo* experiment, no blinding was performed. No sample size calculation was performed. Sample size was based on previous experiments with the LS174T cell line on testing drug effects. Data are shown as mean from at least 2-3 independent experiments. Error bars represent ± SD (standard deviation) or SEM (standard error of the mean), as indicated in each figure legend. The following p-values were considered significant: * P≤0.05; ** P≤0.01; *** P≤0.001; **** P≤0.0001.

### Synthetic procedures for chemical compounds

#### Abbreviations and Acronyms

Aq: aqueous; br: broad; d:doublet; dd, doublet of doublets; m/z: mass-to-charge ratio; m: multiplet; NMR: nuclear magnetic resonance; RT: room temperature; s: singlet; sat. : saturated; t: triplet.

#### General methods for chemistry

All evaporations were carried out *in vacuo* with a rotary evaporator at 10−30 mmHg. Analytical samples were dried under high vacuum (1−5 mmHg) at RT. Thin layer chromatography (TLC) was performed on silica gel plates (Merck) with fluorescent indicator. Spots were visualized by UV light (214 and 254 nm). Flash chromatography was performed using silica gel 60 (Merck). 1H NMR spectra were recorded using a Bruker DPX400 spectrometer (400 MHz) using deuterated solvents. All compounds evaluated in biological tests were purified to ≥ 95% as determined by HPLC−MS on an Agilent/HP 1200 system 6110 mass spectrometer with electrospray ionization (ESI+). Absorbance was monitored at 305 ± 90 and 254 nm.

#### Analytical HPLC-MS

System A: Column ACE 3 C8, 3 µm, 50 x 3.0 mm maintained at 40°C. 0.1% (v/v) trifluoroacetic (TFA) in H_2_O and acetonitrile were used as mobile phases at a flow rate of 1 mL/min, with a gradient time of 3 minutes.

System B: Column Xterra MSC18, 3.5 µm, 50 x 3.0 mm maintained at 40°C. H_2_O

(containing 10 mM ammonium bicarbonate (NH_4_HCO_3_); pH = 10) and acetonitrile were used as mobile phases at a flow rate of 1 mL/min, with a gradient time of 3 minutes.

*Preparative HPLC* was performed on a Gilson HPLC system.

Column ACE C8, 5 µm (150 x 30 mm); 0.1% TFA (v/v) in H_2_O and acetonitrile were used as mobile phases at a flow rate of 35 mL/min with a gradient time of 7 minutes.

The nomenclature of structures was determined using the “Structure to Name” function of MarvinSketch Carbon v.5 and choosing “preferred IUPAC name”.

**N-(6-ethoxy-1,3-benzothiazol-2-yl)-5-nitrofuran-2-carboxamide** (CBK006377, “CBK77” in the main text)

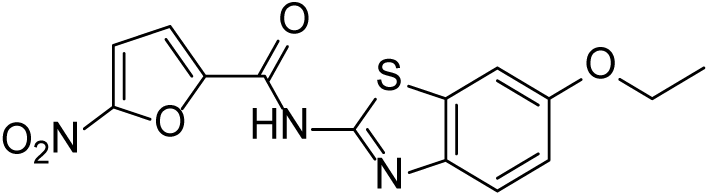

1-Propanephosphonic acid cyclic anhydride (2.2 mL, 3.7 mmol) was added to a solution of 5-nitro-2-furanoic acid (124 mg, 0.64 mmol) in dimethylformamide (DMF). After 10 minutes, 2.2 mL trimethylamine (Et_3_N) (15.9 mmol) and 6-ethoxy-1,3-benzothiazol-2-amine (100 mg, 0.64 mmol) and the mixture left stirring for 1 hour were after ethyla acetate (EtOAc) and H_2_O were added. The organic phase was washed with sodium carbonate (Na_2_CO_3_) (aq, sat.) whereby a solid formed that was filtered off and dissolved in H_2_O and methanol (MeOH). Hydrochloric acid (HCl) (1M) was added dropwise and the final product precipitated as a beige solid. Yield 60 mg. ^1^H NMR (400MHz, DMSO-d_6_) d = 13.51 - 13.08 (m, 1 H), 7.82 (br. s., 2 H), 7.60 (br. s., 2 H), 7.13 - 6.99 (m, 1 H), 4.08 (q, *J* = 6.8 Hz, 4 H), 1.36 (t, *J* = 6.8 Hz, 3 H). ^13^C NMR (101 MHz, DMSO-*d*_6_) δ ppm 14.61 (s, 1 C) 63.64 (s, 1 C) 105.73 (s, 1 C) 113.31 (s, 1 C) 115.71 (s, 1 C) 117.81 (s, 1 C) 129.57 (s, 1 C) 133.41 (s, 1 C) 152.19 (s, 1 C) 155.69 (s, 1 C). LC-MS: m/z calculated for C14H12N3O5S: [M + H]^+^, 334.05; found, 334.1

**N-(4-methoxy-1,3-benzothiazol-2-yl)-5-nitrofuran-2-carboxamide** (CBK085907, “CBK07” in the main text)

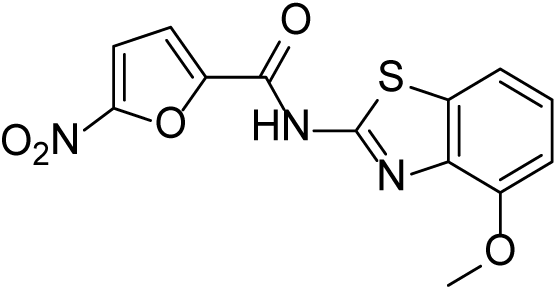

To a solution of 5-nitrofuran-2-carboxylic acid (62 mg, 0.40 mmol) in dry DMF (2mL) were added successively N-methylmorpholine (96 µL, 0.88 mmol) and isobutyl chloroformate (58 µL, 0.44 mmol). The solution was stirred for 15 minutes and then 4-methoxy-1,3-benzothiazol-2-amine (72 mg, 0.40 mmol) was added and the reaction was left at RT stirring for 2 hours. H_2_O (2 mL) was added and the product was filtered off and washed with hot water to of a yellow solid. Yield 65 mg. ^1^H NMR (400 MHz, DMSO-d6) δ ppm 3.93 (s, 3 H) 7.04 (d, J=8.06 Hz, 1 H) 7.30 (t, J=7.98 Hz, 1 H) 7.57 (d, J=7.90 Hz, 1 H) 7.82 (d, J=3.95 Hz, 1 H) 7.88 (br. s., 1 H) 13.57 (br. s., 1 H) ^13^C NMR (101 MHz, DMSO-d6) δ ppm 55.78 (s, 1 C) 107.76 (s, 1 C) 113.35 (s, 1 C) 113.63 (s, 1 C) 117.80 (s, 1 C) 124.83 (s, 1 C) 132.62 (br. s, 1 C) 146.94 (br. s, 1 C) 151.44 (br. s, 1 C) 152.34 (s, 1 C).) LC-MS: m/z calculated for C13H10N2O5S: [M + H]^+^, 320.03; found, 320.0

**Prop-2-yn-1-yl N-[2-(5-nitrofuran-2-amido)-1,3-benzothiazol-6-yl]carbamate** (CBK293384, “CBK77^CLICK^” in the main text)

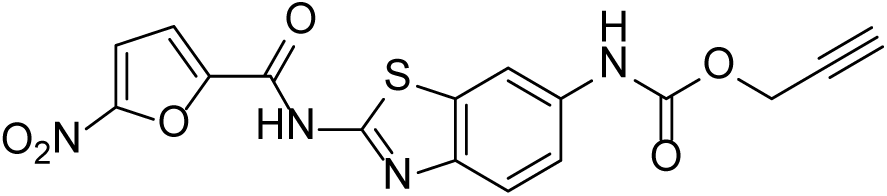

Propargyl chloroformate (30µL, 0.31 µmol) was added at RT to a solution of N-(6-aminobenzo[d]thiazol-2-yl)-5-nitrofuran-2-carboxamide hydrochloride (60mg, 0.18mmol) and methyl morpholine (90µL, 0.65mmol) in DMF (1mL) and the mixture was stirred overnight. Aqueous MeOH was added and the product was filtered off. Washing with aqueous MeOH and drying in vacuum gave the title compound as an orange solid. Yield 38mg. ^1^H NMR (400 MHz, DMSO-*d*_6_) δ ppm 3.58 (t, *J*=2.45 Hz, 1 H) 4.78 (d, *J*=2.37 Hz, 2 H) 7.48 (dd, *J*=8.77, 2.13 Hz, 1 H) 7.65 (d, *J*=8.69 Hz, 1 H) 7.77 (d, *J*=3.47 Hz, 1 H) 7.81 (dd, *J*=3.95, 1.00 Hz, 1 H) 8.09 - 8.12 (m, 1 H) 10.05 (s, 1 H) 13.42 (br. s., 1 H). ^13^C NMR (101 MHz, DMSO-*d*_6_) δ ppm 52.06 (s, 1 C) 77.67 (s, 1 C) 78.92 (s, 1 C) 110.87 (br. s., 1 C) 113.36 (s, 1 C) 117.94 (br. s., 1 C) 118.38 (br. s., 1 C) 135.35 (s, 1 C) 152.31 (s, 1 C) 152.73 (s, 1 C)LC-MS: m/z calculated for C16H11N4O6S: [M + H]^+^, 387.04; found, 387.0

#### Synthesized compounds for SAR analysis

**5-nitro-N-(1,3-thiazol-2-yl)furan-2-carboxamide** (CBK260866)

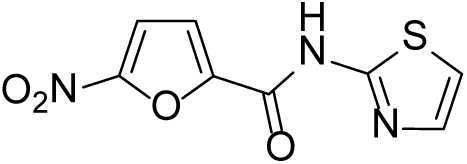

5-Nitrofuran-2-carboxylic acid (70 mg, 0.13 mmol) was treated with thionyl chloride (2 mL) at reflux for 5 hours and then the volatile was evaporated, and the residue was dried in vacuum. The residue was dissolved in dichloromethane (21 mL) and 1/3 of this solution (7 mL) was added to 1,3-thiazol-2-amine (10 mg, 0.10 mmol) and then pyridine (100 µL) was added and the mixture was stirred overnight at RT. The resulting solid was filtered off and washed with MeOH to afford the title compound as a yellow solid. Yield 10 mg. ^1^H NMR (400 MHz, DMSO-d6) δ ppm 7.30 (d, J=3.76 Hz, 1 H) 7.59 (d, J=3.76 Hz, 1 H) 7.73 (br. s., 1 H) 7.80 (d, J=3.76 Hz, 1 H) 13.31 (br. s., 1 H) LC-MS: m/z calculated for C8H6N3O4S: [M + H]^+^, 240.01; found, 240.1

**N-(6-methoxy-1,3-benzothiazol-2-yl)furan-2-carboxamide** (CBK260867)

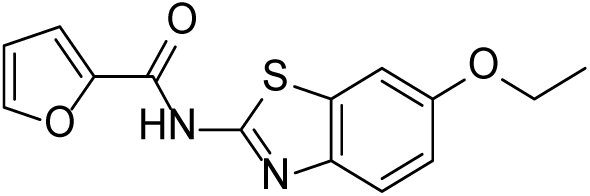

Diisopropylethylamine (21 µL, 0.12 mmol) was added to a mixture of 2-(1H-Benzotriazole-1-yl)-1,1,3,3-tetramethyluronium hexafluorophosphate (HBTU) (45 mg, 0.12 mmol) and furan-2-carboxylic acid (11 mg. 0.10 mmol) in DMF (0.2 mL). After stirring for 15 minutes at RT, 6-methoxy-1,3-benzothiazol-2-amine (19 mg, 0.10 mmol) was added stirring continued over 72 hours. The reaction product was purified by preparative HPLC to give the title compound as a white solid. Yield 19 mg. ^1^H NMR (400 MHz, DMSO-*d*_6_) δ ppm 1.36 (t, *J*=6.95 Hz, 3 H) 4.08 (q, *J*=6.95 Hz, 2 H) 6.73 - 6.78 (m, 1 H) 7.04 (dd, *J*=8.77, 2.61 Hz, 1 H) 7.58 (d, *J*=2.53 Hz, 1 H) 7.65 (d, *J*=8.85 Hz, 1 H) 7.70 (d, *J*=3.48 Hz, 1 H) 8.04 (dd, *J*=1.74, 0.79 Hz, 1 H) 12.12 - 13.24 (m, 1 H). LC-MS: m/z calculated for C14H13N2O3S: [M + H]^+^, 289.06; found, 289.1

**N-(1,3-benzothiazol-2-yl)-5-nitrofuran-2-carboxamide** (CBK260868)

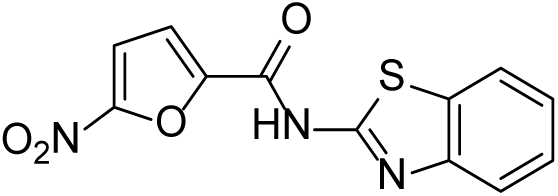

Diisopropylethylamine (21 µL, 0.12 mmol) was added to a mixture of HBTU (45 mg, 0.12 mmol) and 5-nitrofuran-2-carboxylic acid (16 mg. 0.10 mmol) in DMF (0.2 mL). After stirring for 15 minutes at RT, 1,3-benzothiazol-2-amine (15 mg, 0.10 mmol) was added stirring continued over the weekend. Acetonitrile (0.6 mL) and H_2_O (0.2 mL) was added and the mixture was stirred for 30 minutes and the product was filtered off and washed with acetonitrile: H_2_O (1:1) to give the title compound as a brown solid. Yield 14 mg. ^1^H NMR (400 MHz, DMSO-*d*_6_) δ ppm 7.32 - 7.41 (m, 1 H) 7.50 (td, *J*=7.65, 1.25 Hz, 1 H) 7.64 - 7.89 (m, 3 H) 8.02 (d, *J*=7.78 Hz, 1 H) 13.56 (br. s., 1 H). LC-MS: m/z calculated for C12H8N3O4S: [M + H]^+^, 290.02; found, 290.1

**N-(4-methyl-1,3-benzothiazol-2-yl)-5-nitrofuran-2-carboxamide** (CBK260869)

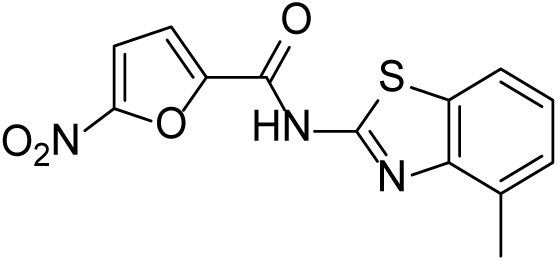

Diisopropylethylamine (21 µL, 0.12 mmol) was added to a mixture of HBTU (45 mg, 0.12 mmol) and 5-nitrofuran-2-carboxylic acid (16 mg. 0.10 mmol) in DMF (0.2 mL). After stirring for 15 minutes at RT, 4-methyl-1,3-benzothiazol-2-amine (16 mg, 0.10 mmol) was added stirring continued for 3 hours. Acetonitrile (0.4 mL) and H_2_O (1 mL) was added and the mixture was stirred for 30 minutes and the product was filtered off and washed with acetonitrile:H_2_O (1:1) to give the title compound as a beige solid. Yield 16 mg. ^1^H NMR (400 MHz, DMSO-*d*_6_) δ ppm 2.61 (s, 3 H) 7.22 - 7.34 (m, 2 H) 7.75 - 7.87 (m, 2 H) 7.95 (br. s., 1 H) 13.44 (br. s., 1 H). LC-MS: m/z calculated for C13H10N3O4S: [M + H]^+^, 304.03; found, 304.1

**N-(6-methyl-1,3-benzothiazol-2-yl)-5-nitrofuran-2-carboxamide** (CBK260870)

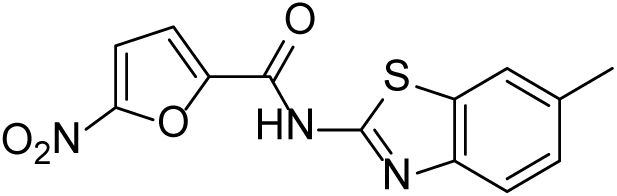

Diisopropylethylamine (21 µL, 0.12 mmol) was added to a mixture of HBTU (45 mg, 0.12 mmol) and 5-nitrofuran-2-carboxylic acid (16 mg. 0.10 mmol) in DMF (0.2 mL). After stirring for 15 minutes at RT, 6-methyl-1,3-benzothiazol-2-amine (16 mg, 0.10 mmol) was added stirring continued for 3 hours. Acetonitrile (0.4 mL) and H_2_O (1 mL) was added and the mixture was stirred for 30 minutes and the product was filtered off and washed with acetonitrile:H_2_O (1:1) to give the title compound as a beige solid. Yield 12mg. ^1^H NMR (400 MHz, DMSO-*d*_6_) δ ppm 2.42 (s, 3 H) 7.27 - 7.33 (m, 1 H) 7.59 (br. s., 1 H) 7.73 - 7.84 (m, 3 H) 13.14 - 13.95 (m, 1 H)

LC-MS: m/z calculated for C13H10N3O4S: [M + H]+, 304.03; found, 304.1

**N-(6-acetamido-1,3-benzothiazol-2-yl)-5-nitrofuran-2-carboxamide** (CBK277750)

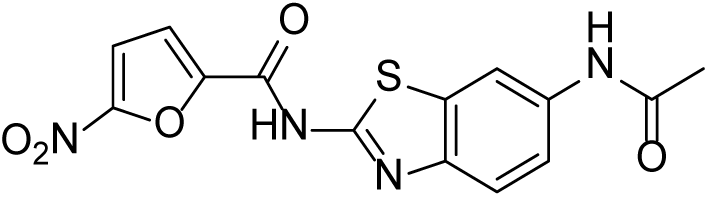

Acetic anhydride (10 µL, 0.11 mmol) was added at RT to a solution of N-(6-amino-1,3-benzothiazol-2-yl)-5-nitrofuran-2-carboxamide hydrochloride (10 mg, 0.03 mmol) in pyridine (0.2 mL) and dichloromethane (0.5 mL). After 3 hours the solvents were evaporated, and the residue was dried in vacuum over the weekend. The residue was washed with H_2_O and a small amount of aqueous MeOH to give the title compound as a red solid. Yield 3 mg. ^1^H NMR (400 MHz, DMSO-*d*_6_) δ ppm 2.07 (s, 3 H) 7.52 - 7.60 (m, 1 H) 7.65 (br. s., 1 H) 7.74 - 7.95 (m, 2 H) 8.31 (s, 1 H) 10.15 (s, 1 H) 13.49 (br. s, 1 H). LC-MS: m/z calculated for C14H11N4O5S: [M + H]^+^, 347.03, 347.1

**N-(6-chloro-1,3-benzothiazol-2-yl)-5-nitrofuran-2-carboxamide** (CBK277751)

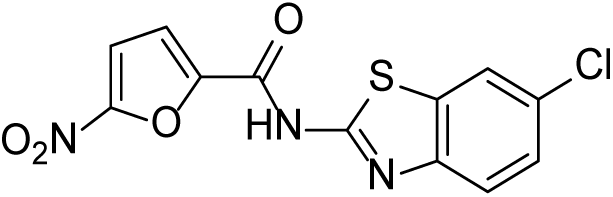

To a solution of 5-nitrofuran-2-carboxylic acid (31 mg, 0.20 mmol) in dry DMF (1 mL) cooled at 0°C under N_2_, were added successively N-methylmorpholine (48 µL, 0.44 mmol) and isobutyl chloroformate (29µL, 0.22mmol). The solution was stirred for 15 minutes at 0°C and then 15 minutes at RT. 6-chloro-1,3-benzothiazol-2-amine (37 mg, 0.20 mmol) was added and the reaction was left at RT stirring. The product was filtered off. Recrystallization from aqueous MeOH gave the title compound as a yellow solid. Yield 19 mg. ^1^H NMR (400 MHz, DMSO-*d*_6_) δ ppm 7.52 (dd, *J*=8.53, 2.26 Hz, 1 H) 7.76 (br. s., 1 H) 7.84 (d, *J*=3.76 Hz, 1 H) 7.85 - 7.96 (br. s, 1 H) 8.18 (d, *J*=1.51 Hz, 1 H) 13.53 (br. s., 1 H). LC-MS: m/z calculated for C12H7ClN3O4S: [M + H]^+^, 323.98, found, 324.0

**N-(4-chloro-1,3-benzothiazol-2-yl)-5-nitrofuran-2-carboxamide** (CBK277752)

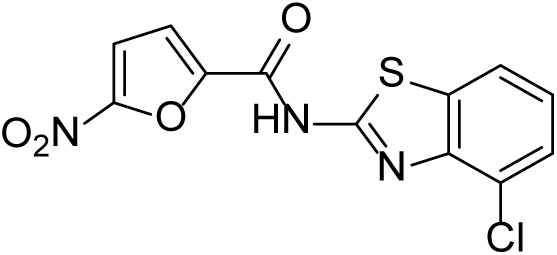

To a solution (0°C) of 5-nitrofuran-2-carboxylic acid (31 mg, 0.20 mmol) in dry DMF (1 mL), were added successively N-methylmorpholine (48 µL, 0.44 mmol) and isobutyl chloroformate (29 µL, 0.22 mmol). The solution was stirred for 15 minutes at 0°C and then 15 minutes at RT. 4-chloro-1,3-benzothiazol-2-amine hydrobromide (53 mg, 0.20 mmol) was added and the reaction was left at RT stirring. The product was filtered off. Recrystallization from aqueous MeOH gave the title compound as a yellow solid. Yield 21 mg. ^1^H NMR (400 MHz, DMSO-*d*_6_) δ ppm 2.61 (s, 3 H) 7.22 - 7.34 (m, 2 H) 7.75 - 7.87 (m, 2 H) 7.95 (br. s., 1 H) 13.44 (br. s., 1 H). LC-MS: m/z calculated for C12H7ClN3O4S: [M + H]^+^, 324.00, found, 324.0

**N-(5-chloro-1,3-benzoxazol-2-yl)-5-nitrofuran-2-carboxamide** (CBK277754)

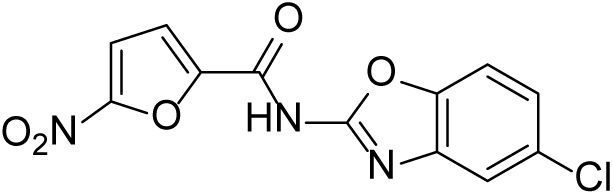

To a solution (0°C) of 5-nitrofuran-2-carboxylic acid (31 mg, 0.20 mmol) in dry DMF (1 mL), were added successively N-methylmorpholine (48 µL, 0.44 mmol) and isobutyl chloroformate (29 µL, 0.22 mmol). The solution was stirred for 15 minutes at 0°C and then 15 minutes at RT. 5-chloro-1,3-benzothiazol-2-amine (34 mg, 0.20 mmol) was added and the reaction was left at RT stirring. The product was filtered off. Purification by preparative HPLC gave the title compound as a brown solid. Yield 5 mg ^1^H NMR (400 MHz, DMSO-*d*_6_) δ ppm 7.37 (dd, *J*=8.69, 2.05 Hz, 1 H) 7.63 (d, *J*=18.17 Hz, 2 H) 7.70 (d, *J*=8.69 Hz, 1 H) 7.79 (d, *J*=3.79 Hz, 1 H) 13.03 (br. s, 1 H). ^13^C NMR (101 MHz, DMSO-*d*_6_) δ ppm 109.44 (s, 1 C) 113.93 (s, 1 C) 114.62 (s, 1 C) 116.22 (s, 1 C) 120.62 (s, 1 C) 126.75 (s, 1 C) 145.04 (br. s, 1 C) 145.61 (br. s, 1 C) 150.94 (s, 1 C) 161.51 (s, 1 C) LC-MS: m/z calculated for C12H7ClN3O5: [M + H]^+^, 308.00, found, 308.0

**N-(1H-1,3-benzodiazol-2-yl)-5-nitrofuran-2-carboxamide** (CBK277755)

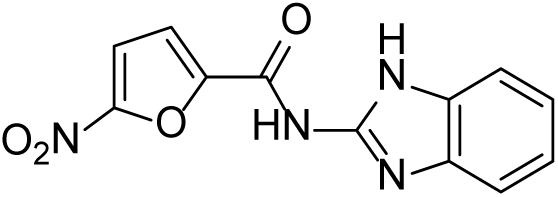

To a solution (0°C) of 5-nitrofuran-2-carboxylic acid (31 mg, 0.20 mmol) in dry DMF (1 mL), were added successively N-methylmorpholine (48 µL, 0.44 mmol) and isobutyl chloroformate (29 µL, 0.22 mmol). The solution was stirred for 15 minutes at 0°C and then 15 minutes at RT. 1H-1,3-benzodiazol-2-amine (27 mg, 0.20 mmol) was added and the reaction was left at RT stirring. The product was filtered off. Purification by preparative HPLC gave the title compound as a yellow solid. Yield 4 mg. ^1^H NMR (400 MHz, DMSO-*d*_6_) δ ppm 7.21 - 7.24 (m, 2 H) 7.32 (d, *J*=3.79 Hz, 1 H) 7.41 - 7.45 (m, 2 H) 7.73 (d, *J*=3.79 Hz, 1 H) 12.72 (br. s., 2 H)LC-MS: m/z calculated for C12H9N4O4: [M + H]^+^, 273.05, found, 273.1

**N-(6-amino-1,3-benzothiazol-2-yl)-5-nitrofuran-2-carboxamide hydro chloride** (CBK277756)

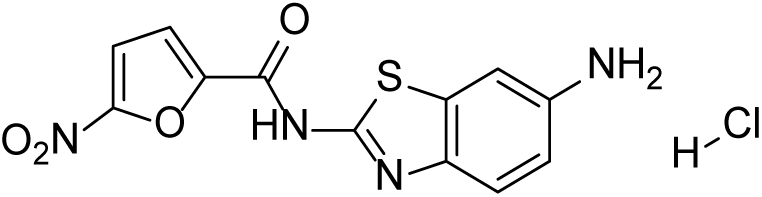

Methylmorpholine (0.22 mL, 2 mmol) and isobutyl chloroformate (0.26 mL, 2 mmol) were added to a solution of 5-nitrofuran-2-carboxylic acid (0.31 g, 2 mmol) in DMF (4 mL) at 0°C. The temperature was allowed to reach ca 10°C during 1 hour and tert-butyl N-(2-amino-1,3-benzothiazol-6-yl)carbamate (0.14 g, 0.52 mmol) was added. After 1 hour, more tert-butyl N-(2-amino-1,3-benzothiazol-6-yl)carbamate (0.39 g, 1.47 mmol) was added and the reaction was left stirring on at RT. MeOH (1 mL) and H_2_O (1 mL) was added. The mixture was cooled in ice H_2_O and filtered, the solid was washed with 70% cold MeOH. The crystals were dried in vacuum to yield 0.22 g. The material was treated with concentrated HCl (2 mL) at 100°C for 10 minutes. After cooling down to RT, the light-yellow solid was filtered and carefully washed with a small amount of cold H_2_O. The solid was dried in vacuum. The color quickly becomes dark brown. Yield 70 mg. ^1^H NMR (400 MHz, DMSO-*d*_6_) δ ppm 7.45 (dd, *J*=8.53, 2.21 Hz, 1 H) 7.81 (d, *J*=8.69 Hz, 1 H) 7.84 (d, *J*=3.95 Hz, 1 H) 7.85 - 7.89 (m, 1 H) 8.01 (d, *J*=2.05 Hz, 1 H) 10.38 (br. s., 1 H). ^13^C NMR (101 MHz, DMSO-*d*_6_) δ ppm 113.29 (s, 1 C) 116.12 (s, 1 C) 118.31 (s, 1 C) 121.46 (s, 1 C) 129.06 (s, 1 C) 152.38 (s, 1 C). LC-MS: m/z calculated for C12H9N4O4S: [M + H]^+^, 305.03, found, 305.1

**N-(4,5-dichloro-1,3-benzothiazol-2-yl)-5-nitrofuran-2-carboxamide** (CBK277757)

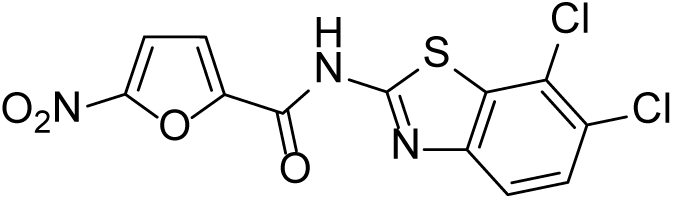

DMF (2.5 µL) was added to a stirred mixture of 5-nitrofuran-2-carboxylic acid (50 mg, 0.32 mmol) and oxalyl chloride (50 µL) in dichloromethane (1 mL) at RT. After 30 minutes the solvents were evaporated, and the residue dried in vacuum. The solid material was dissolved in dichloromethane (4 mL) and treated with 4,5-dichloro-1,3-benzothiazol-2-amine (70 mg, 0.32 mmol). Triethylamine (50 µL) was added and the mixture was stirred at RT overnight. Water and EtOAc was added and the organic phase were evaporated, and the residue purified by preparative HPLC to give the target compound. Yield 3 mg. ^1^H NMR (400 MHz, DMSO-*d*_6_) δ ppm 7.67 (d, *J*=3.76 Hz, 1 H) 7.76 (d, *J*=8.78 Hz, 1 H) 7.83 (d, *J*=4.02 Hz, 1 H) 7.87 (d, *J*=8.53 Hz, 1 H) 10.84 (s, 1 H) LC-MS: m/z calculated for C12H6Cl2N3O4SxNH3: [M + H]^+^, 374.97; found, 375.1

**N-(6-methyl-1,3-benzothiazol-2-yl)-5-nitrothiophene-2-carboxamide** (CBK277772)

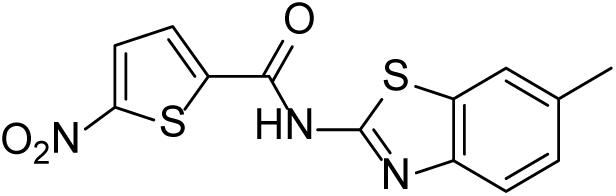

1-Propanephosphonic acid cyclic anhydride (227 µL, 0.38 mmol) was added to a solution of 5-nitro-2-thiophenecarboxylic acid (25 mg, 0.16 mmol) in DMF (0.5 mL). After 30 minutes triethylamine (0.11 mL, 0.8 mmol) was added. After another 30 minutes 6-methyl-1,3-benzothiazol-2-amine (20 mg, 0.12 µmol) was added and stirred overnight. MeOH (0.5 mL) was added and the volatile solvents were evaporated. Addition of 0.5-1 mL of MeOH and H_2_O (1-2 mL) and filtration and drying in vacuum gave the title compound as a brown solid. Yield 15 mg. ^1^H NMR (400 MHz, DMSO-*d*_6_) δ ppm 2.41 (s, 3 H) 7.29 (dd, *J*=8.29, 1.18 Hz, 1 H) 7.57 (d, *J*=8.21 Hz, 1 H) 7.77 (s, 1 H) 8.03 (br. s., 1 H) 8.17 (d, *J*=4.26 Hz, 1 H) 12.79 - 14.19 (m, 1 H). LC-MS: m/z calculated for C13H10N3O3S2: [M + H]^+^, 320.01; found, 320.1

**N-(5-chloro-4-methyl-1,3-benzothiazol-2-yl)-5- nitrothiophene-2-carboxamide** (CBK277775)

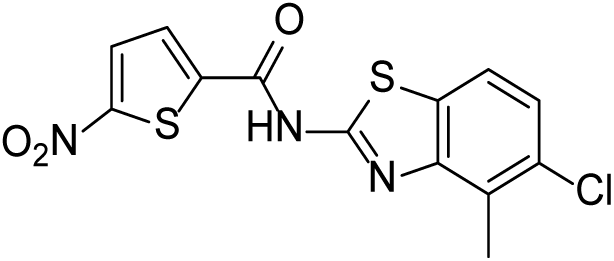

1-Propanephosphonic acid cyclic anhydride (227 µL, 0.38 mmol) was added to a solution of 5-nitro-2-thiophenecarboxylic acid (25 mg, 0.16 mmol) in DMF (0.5 mL). After 30 minutes triethylamine (0.11 mL, 0.8 mmol) was added. After another 30 minutes 5-chloro-4-methyl-1,3-benzothiazol-2-amine (25 mg, 0.12 µmol) was added and stirred overnight. MeOH (0.5 mL) was added and the volatile solvents were evaporated. Addition of 0.5-1 mL of MeOH and H_2_O (1-2 mL) and filtration and drying in vacuum gave the title compound as a brown solid. Yield 12 mg. ^1^H NMR (400 MHz, DMSO-*d*_6_) δ ppm 2.31 (s, 3 H) 7.54 (d, *J*=8.53 Hz, 1 H) 7.71 (d, *J*=8.78 Hz, 1 H) 8.03 (d, *J*=4.27 Hz, 1 H) 8.23 (d, *J*=4.27 Hz, 1 H) 10.77 (s, 1 H). LC-MS: m/z calculated for C13H8ClN3O3S: [M + NH4]+, 371.00; found, 371.0

**N-(4,6-dichloro-1,3-benzothiazol-2-yl)-5- nitrothiophene-2-carboxamide** (CBK277776)

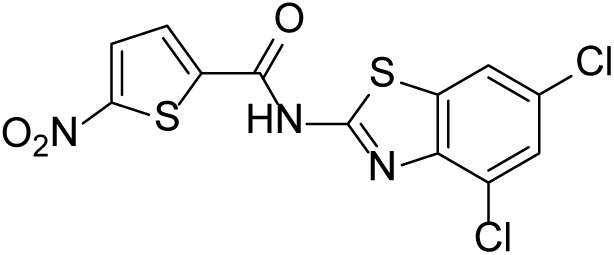

1-Propanephosphonic acid cyclic anhydride (227 µL, 0.38 mmol) was added to a solution of 5-nitro-2-thiophenecarboxylic acid (25 mg, 0.16mmol) in DMF (0.5 mL). After 30 minutes triethylamine (0.11 mL, 0.8 mmol) was added. After another 30 minutes 4,6-dichloro-4-1,3-benzothiazol-2-amine (25 mg, 0.11 µmol) was added and stirred overnight. MeOH (0.5 mL) was added and the volatile solvents were evaporated. Addition of 0.5-1 mL of MeOH and H_2_O (1-2 mL) and filtration and drying in vacuum gave the title compound as a brown solid. Yield 13 mg. ^1^H NMR (400 MHz, DMSO-*d*_6_) δ ppm 7.72 (d, *J*=2.01 Hz, 1 H) 8.22 (d, *J*=2.01 Hz, 1 H) 8.23 (d, *J*=4.27 Hz, 1 H) 8.34 (d, *J*=4.52 Hz, 1 H) 13.82 (br. s., 1 H). LC-MS: m/z calculated for C12H6Cl2N3O3S2: [M + H]^+^, 373.91; found, 374.0

**N-(6-bromo-1,3-benzothiazol-2-yl)-5-nitrofuran-2-carboxamide** (CBK277778)

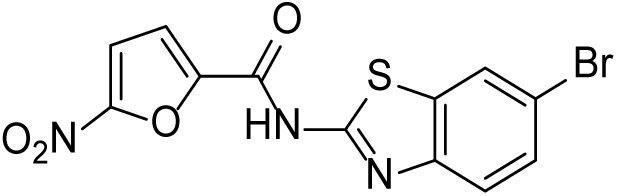

1-Propanephosphonic acid cyclic anhydride (250 µL, 0.4 mmol) was added to a solution of 5-nitro-2-furanoic acid (32 mg, 0.2 mmol)) in DMF (1mL). After 30 minutes triethylamine (125 µL, 0.9 mmol) was added. After another 30 minutes 6-bromo-1,3- benzothiazol-2-amine (29 mg, 0.13 µmol) was added and stirred overnight. MeOH (1mL) was added and the volatile solvents were evaporated. H_2_O was added and a solid was filtered off and dried in vacuum to afford the title compound as a brown solid. Yield 30 mg. ^1^H NMR (400 MHz, DMSO-*d*_6_) δ ppm 7.64 (dd, *J*=8.53, 2.01 Hz, 1 H) 7.69 (br. s., 1 H) 7.84 (d, *J*=3.76 Hz, 1 H) 7.88 (br. s., 1 H) 8.31 (s, 1 H) 13.56 (br. s., 1 H). LC-MS: m/z calculated for C12H7BrN3O4S: [M + H]^+^, 367.93; found, 368.0

**N-(5-chloro-4-methyl-1,3-benzothiazol-2-yl)-5-nitrofuran-2-carboxamide** (CBK277779)

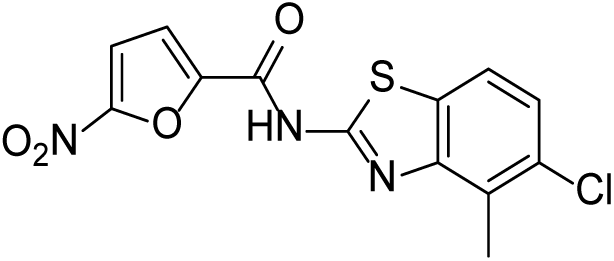

1-Propanephosphonic acid cyclic anhydride (250 µL, 0.4 mmol) was added to a solution of 5-nitro-2-furanoic acid (25 mg, 0.16 mmol) in DMF (1 mL). After 30 minutes triethylamine (125 µL, 0.9 mmol) was added. After another 30 minutes 5-chloro-4-methyl-1,3-benzothiazol-2-amine (26 mg, 0.13 µmol) was added and stirred overnight. MeOH (1 mL) was added and the volatile solvents were evaporated. H_2_O was added and a solid was filtered off and dried in vacuum to afford the title compound as a beige solid. Yield 9 mg. ^1^H NMR (400 MHz, DMSO-*d*_6_) δ ppm 7.52 (d, *J*=8.53 Hz, 1 H) 7.63 (d, *J*=4.02 Hz, 1 H) 7.71 (d, *J*=8.53 Hz, 1 H) 7.84 (d, *J*=4.02 Hz, 1 H) 10.74 (s, 1 H). LC-MS: m/z calculated for C13H8ClN3O4S: [M + NH4]+, 355.03; found, 355.1

**5-nitro-N-(6-nitro-1,3-benzothiazol-2-yl)furan-2-carboxamide** (CBK277781)

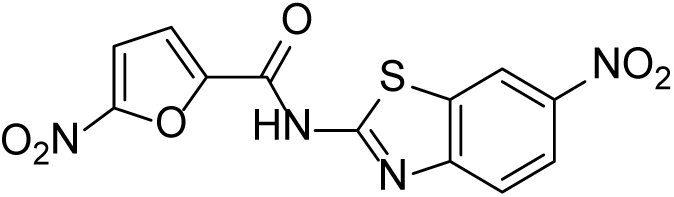

1-Propanephosphonic acid cyclic anhydride (250 µL, 0.4 mmol) was added to a solution of 5-nitro-2-furanoic acid (25 mg, 0.16 mmol) in DMF (1 mL). After 30 minutes triethylamine (125 µL, 0.9 mmol) was added. After another 30 minutes 6-nitro-1,3- benzothiazol-2-amine (25 mg, 0.13 µmol) was added and stirred overnight. MeOH (1 mL) was added and the volatile solvents were evaporated. H_2_O was added and a solid was filtered off and dried in vacuum to afford the title compound as a brown solid. Yield 25 mg. ^1^H NMR (400 MHz, DMSO-*d*_6_) δ ppm 7.85 (d, *J*=4.02 Hz, 1 H) 7.88 - 7.98 (m, 2 H) 8.33 (dd, *J*=8.91, 2.38 Hz, 1 H) 9.11 (d, *J*=2.51 Hz, 1 H) 13.88 (br. s., 1 H). LC-MS: m/z calculated for C12H7N4O6S: [M + H]^+^, 335.00; found, 335.1

**N-(4,6-dichloro-1,3-benzothiazol-2-yl)-5-nitrofuran-2-carboxamide** (CBK277782)

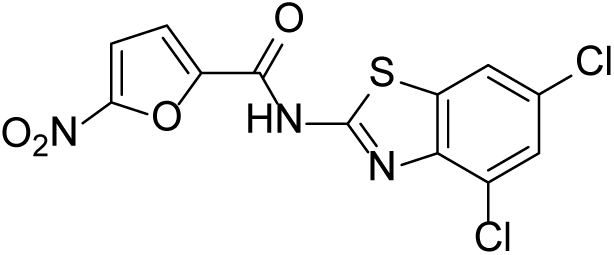

1-Propanephosphonic acid cyclic anhydride (250 µL, 0.4 mmol) was added to a solution of 5-nitro-2-furanoic acid (32 mg, 0.2 mmol) in DMF (1mL). After 30 minutes triethylamine (125 µL, 0.9 mmol) was added. After another 30 minutes 4,6-dichloro- 1,3-benzothiazol-2-amine (28 mg, 0.13 µmol) was added and stirred overnight. MeOH (1 mL) was added and the volatile solvents were evaporated. H_2_O was added and a solid was filtered off and dried in vacuum to afford the title compound as a brown solid. Yield 7 mg. ^1^H NMR (400 MHz, DMSO-*d*_6_) δ ppm 7.72 (d, *J*=2.05 Hz, 1 H) 7.86 (d, *J*=3.95 Hz, 1 H) 8.01 (d, *J*=3.95 Hz, 1 H) 8.21 (d, *J*=1.90 Hz, 1 H) 13.79 (br. s., 1 H). LC-MS: m/z calculated for C12H6Cl2N3O4S: [M + H]^+^, 357.94; found, 357.9

**Preparation of *in vivo* solutions**

A Captisol solution was made by dissolving 4 g Captisol in H_2_O up to a final volume of 10 mL during gentle stirring for 30 minutes. Sodium 6-ethoxy-N-(5-nitrofuran-2- carbonyl)-1,3-benzothiazol-2-aminide (25 mg) was added and gentle stirring continued for 1 hour. Tris buffer (2.5 mL, 25 mM, pH 8.3) was added and the solution was pH adjusted to ca pH = 7 with 1M HCl. The solution was filtered through a 45 micron filter and kept at 4°C.

Another solution (vehicle) was made as above with the exception that no Sodium 6- ethoxy-N-(5-nitrofuran-2-carbonyl)-1,3-benzothiazol-2-aminide was added.

**Sodium 6-ethoxy-N-(5-nitrofuran-2-carbonyl)-1,3-benzothiazol-2-aminide** (CBK291422N, sodium salt of CBK77)

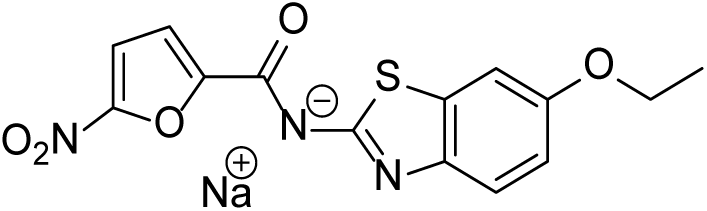

20 mg of the solid was partly dissolved in EtOH and sodium hydroxide solution (5x10µL, 2M) was added to the mixture until the pH was basic and the color had changed to deep red-brown. By then the sodium salt has precipitated. The mixture was filtered and the solid washed with a little EtOH. Gave a dark violet solid. ^1^H NMR (400 MHz, DMSO-*d*_6_) δ ppm 1.30 - 1.37 (m, 3 H) 3.97 - 4.07 (m, 2 H) 6.83 (dd, *J*=8.69, 2.69 Hz, 1 H) 7.16 (d, *J*=3.79 Hz, 1 H) 7.28 (d, *J*=2.53 Hz, 1 H) 7.35 - 7.41 (m, 1 H) 7.71 (d, *J*=3.79 Hz, 1 H). ^13^C NMR (101 MHz, DMSO-*d*_6_) δ ppm 14.86 (s, 1 C) 63.41 (s, 1 C) 105.08 (s, 1 C) 113.40 (s, 1 C) 113.67 (s, 1 C) 114.31 (s, 1 C) 119.38 (s, 1 C) 134.31 (s, 1 C) 144.59 (s, 1 C) 150.77 (s, 1 C) 153.72 (s, 1 C) 156.32 (s, 1 C) 160.98 (s, 1 C) 166.75 (s, 1 C) LC-MS: m/z calculated for C14H12N3O5S: [M + H]^+^, 334.05; found, 334.1

## Supplementary Material

**Figure S1.**
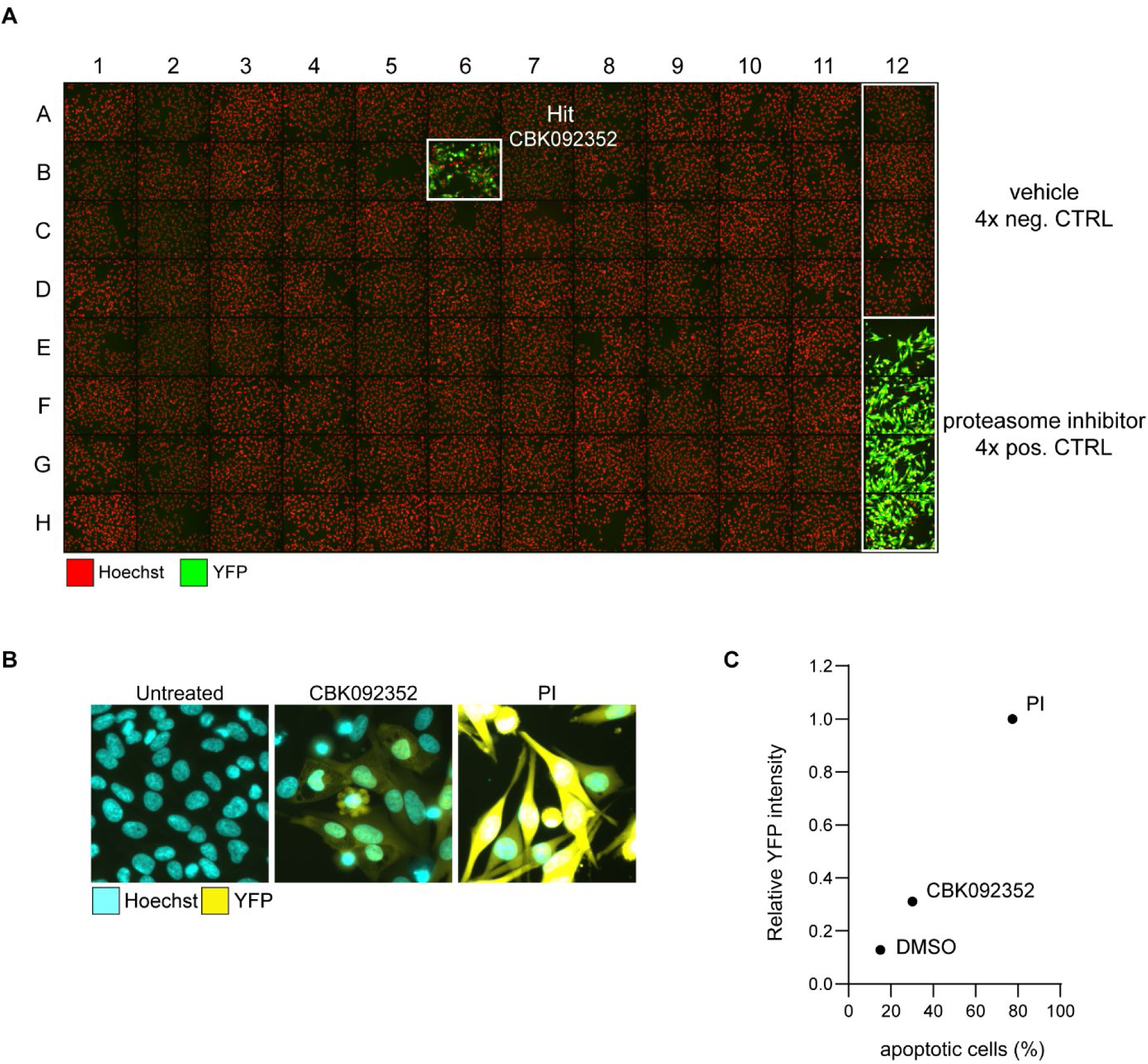
Screen for UPS inhibitors. **A)** Overview of plate #35 from the screen, in which the active compound was found in position B6 (hit compound CBK092352). **B)** Representative images of MelJuSo Ub-YFP cells from the indicated conditions in the screen plate #35. **C)** Scatter plot showing the percentage of apoptotic cells (identified as fragmented nuclei based on the Hoechst staining) and the average YFP-intensity in the nucleus relative to the intensity in epoxomicin (proteasome inhibitor, PI) treated cells (relative YFP-intensity).

**Figure S2.**
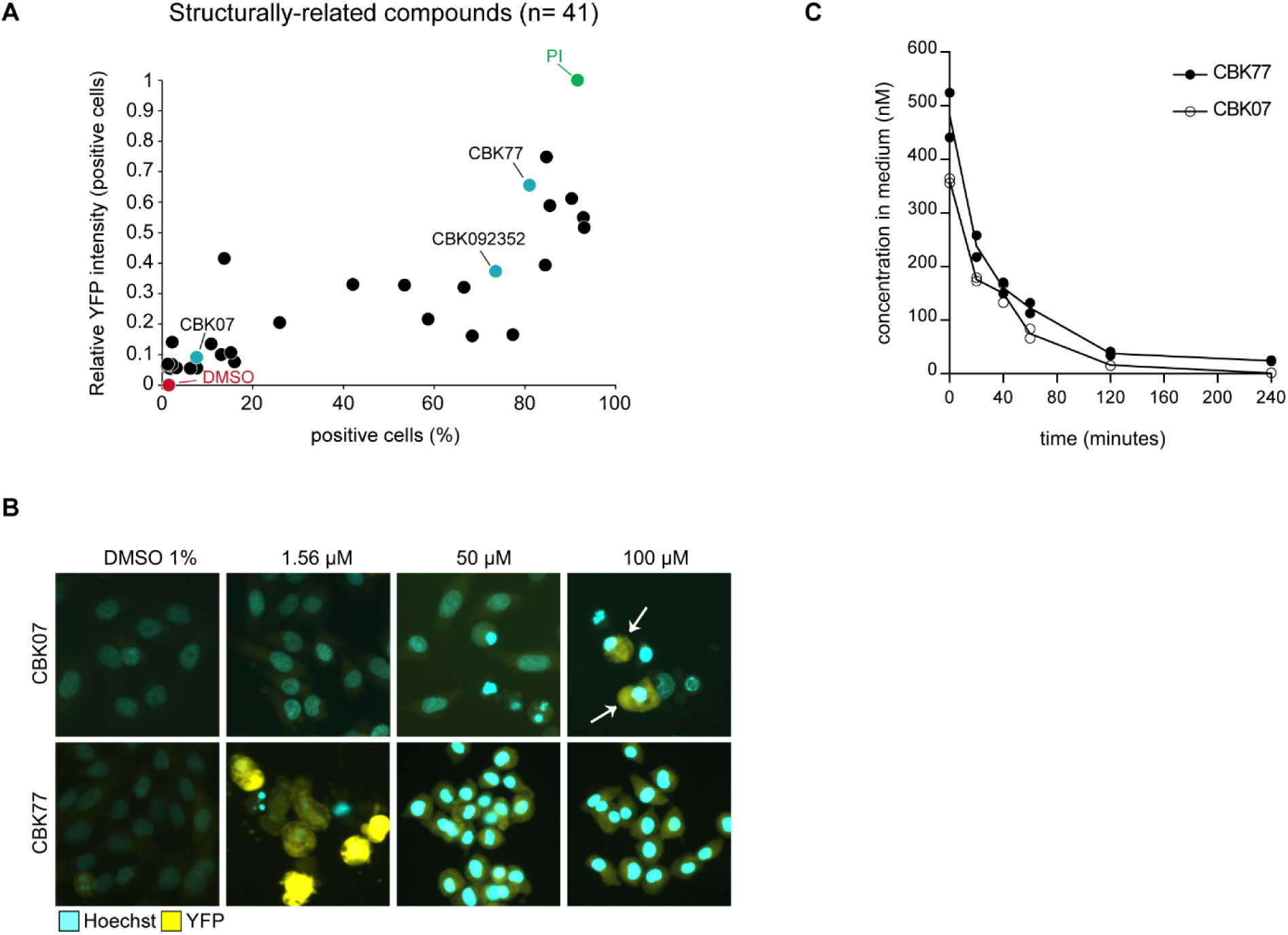
CBK77 and CBK07 are both taken up by cells while only CBK77 efficiently blocks UPS activity. **A)** MelJuSo Ub-YFP cells were treated for 16 hours with selected compounds related to the benzothiazole hit molecule (10 µM). Epoxomicin (proteasome inhibitor = PI, 100 nM) was included as a positive control. After the treatment, nuclei were counterstained with Hoechst and cells were imaged live with a widefield automated microscope. The nuclear YFP intensity per cell was quantified using MetaXpress. Data are shown as the normalized YFP intensity to epoxomicin versus the percentage of positive cells. Data can be found in Supplementary Table S2. **B)** Representative images of MelJuSo Ub-YFP cells treated for 24 hours with CBK77 or CBK07 at the indicated concentrations. Nuclei were counterstained with Hoechst and after fixation cells imaged with an automated widefield microscope. **C)** MelJuSo parental cells were treated with 1 µM of CBK77 for 20, 40, 60, 120 and 240 minutes. After incubation, the medium was removed an analyzed by mass spectrometry to calculate the amount of compound left in the medium. In parallel, the content of the cells was solubilized with a 60:40 acetonitrile:H_2_O mixture (50 mM acetonitrile final concentration) and analyzed by mass spectrometry to calculate the amount of compound inside the cells. The compound could not be detected in this latter analysis, probably due to a fast metabolism-related reaction, or due to covalent binding to a target.

**Figure S3.**
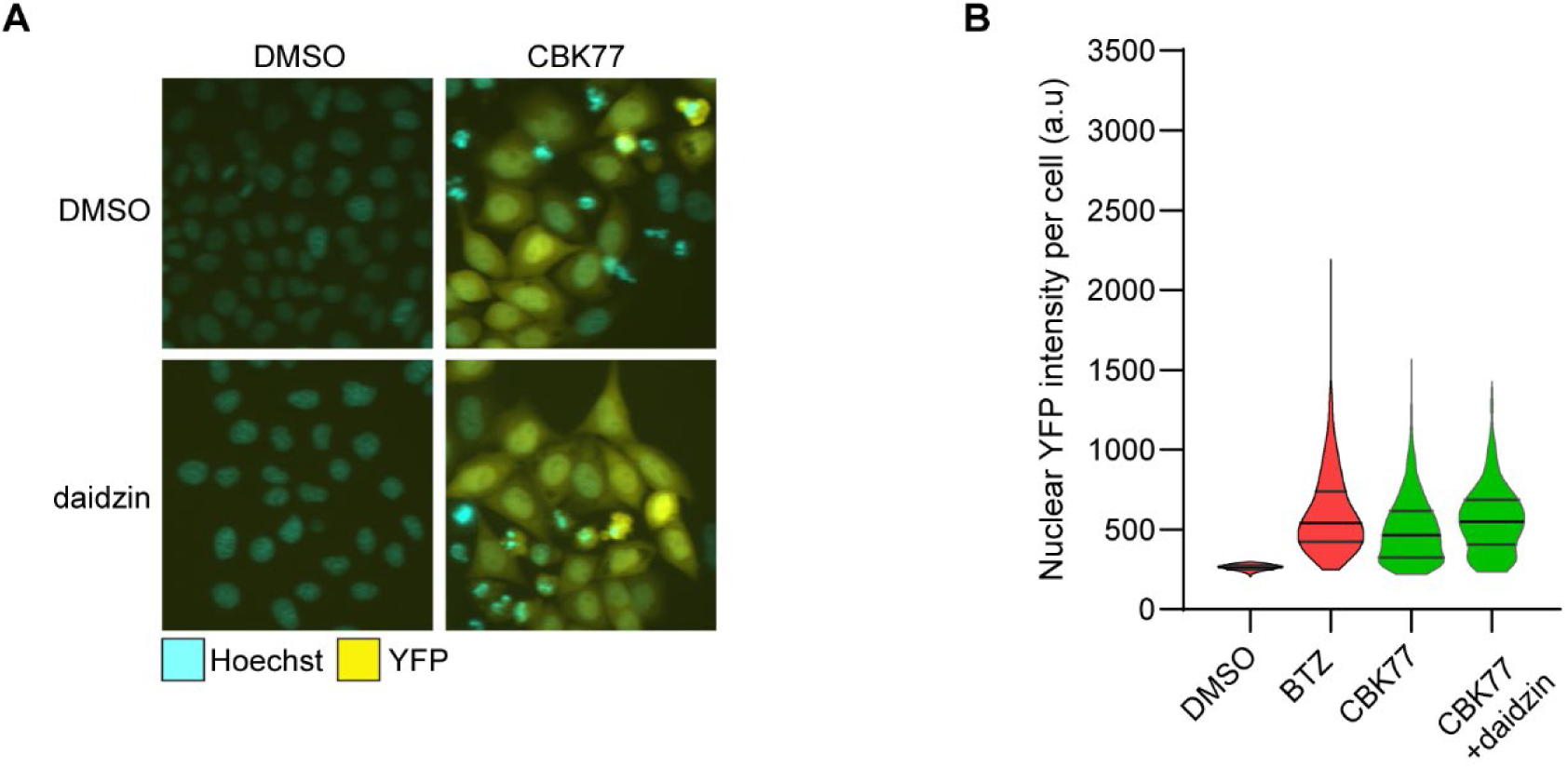
ALDH-2 is not involved in bioactivation of CBK77. **A)** Representative images of HeLa Ub-YFP cells treated for 24 hours with DMSO 0.2% or daidzin 100 µM with or without co-treatment with CBK77 20 µM. Nuclei were counterstained with Hoechst and cells were imaged with an automated widefield microscope. **B)** Quantification of (**A**). The nuclear YFP intensity per cell was quantified using the MetaXpress software. Frequency and distribution of the measured YFP intensity per cell are shown as violin plots. n= 1798 cells (DMSO); n = 509 cells (bortezomib); n = 912 cells (CBK77) and n= 493 cells (CBK77+daidzin) from a representative experiment. Black lines within each distribution represent the median and the upper and lower interquartile range limits.

**Figure S4.**
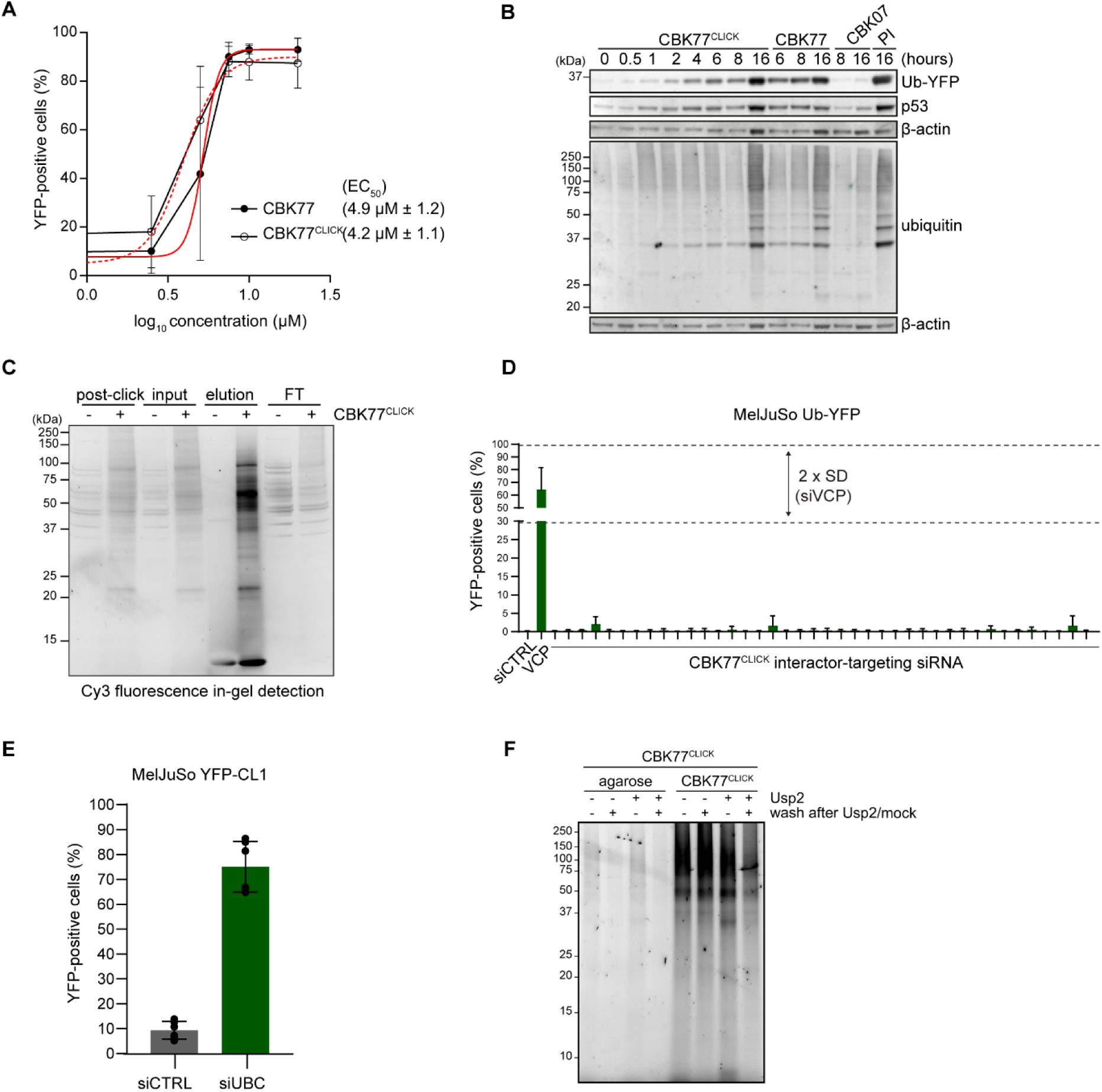
Characterization of the activity probe CBK77^CLICK^ and related experiments to Fig. 7. **A)** Concentration-response experiments performed with MelJuSo Ub-YFP cells. Cells were treated for 6 hours with a range of compound concentrations. Nuclei were stained with Hoechst 33342 and cells directly imaged live with an automated widefield microscope. The number of cells accumulating the UFD reporter was quantified using the MetaXpress software. Data are represented as mean ± SD of three independent experiments. Non-linear curve fitting is depicted in red. The effective concentration (EC) at which 50% of the cells accumulate the UFD reporter above the basal threshold (EC_50_) upon CBK77^CLICK^ treatment is shown (4.2 µM). **B)** MelJuSo Ub-YFP cells were treated with either CBK77^CLICK^, CBK77, CBK07 (10 µM) or epoxomicin (proteasome inhibitor = PI, 100 nM) and harvested at the indicated timepoints. Cell lysates were analyzed by immunoblotting with the indicated antibodies. **C)** MelJuSo parental cells were treated with CBK77^CLICK^ (20 µM) for 1 hour. After labelling of the compound with the trifunctional linker TAMRA-azide-biotin, the labelled compound was pulled down with streptavidin beads. **D)** siRNA subscreen with target genes selected from the CBK77^CLICK^ interactome mass spectrometry dataset. MelJuSo Ub-YFP cells were transfected with the indicated siRNAs for 48 hours. Nuclei were counterstained with Hoechst and after fixation cells imaged with an automated widefield microscope. Data are shown as mean percentage of YFP-positive cells ± SD of three independent experiments. siCTRL and the ubiquitin-selective segregase siVCP/p97, which is critical for the degradation of Ub-YFP, were used as negative and positive controls, respectively. **E)** MelJuSo YFP-CL1 cells were depleted of ubiquitin for 24 hours with the indicated siRNAs (siUBC, 30 nM). Non-targeting siRNA (siCTRL) was used as control. After the treatment, nuclei were counterstained with Hoechst and cells were imaged live in a widefield automated microscope. The average percentage of YFP-positive cells in the data pooled from two independent experiments (with three technical replicates each, shown as aligned dots) ± SD is shown as a bar plot. **F)** MelJuSo Ub-YFP cells were treated with CBK77^CLICK^ (10 µM) for 1 hour. After cell lysis and labelling of the compound with a TAMRA-azide fluorophore, ubiquitin was pulled down with either control agarose beads or TUBE-agarose beads. After incubation, the beads were treated with or without Usp2 for 1 hour. A set of samples were stopped directly with addition of LDS-sample buffer, while a second set was further washed to remove unbound proteins. Binding of the compound to ubiquitin was assessed via TAMRA detection and immunoblotting with a ubiquitin antibody.

**Figure S5.**
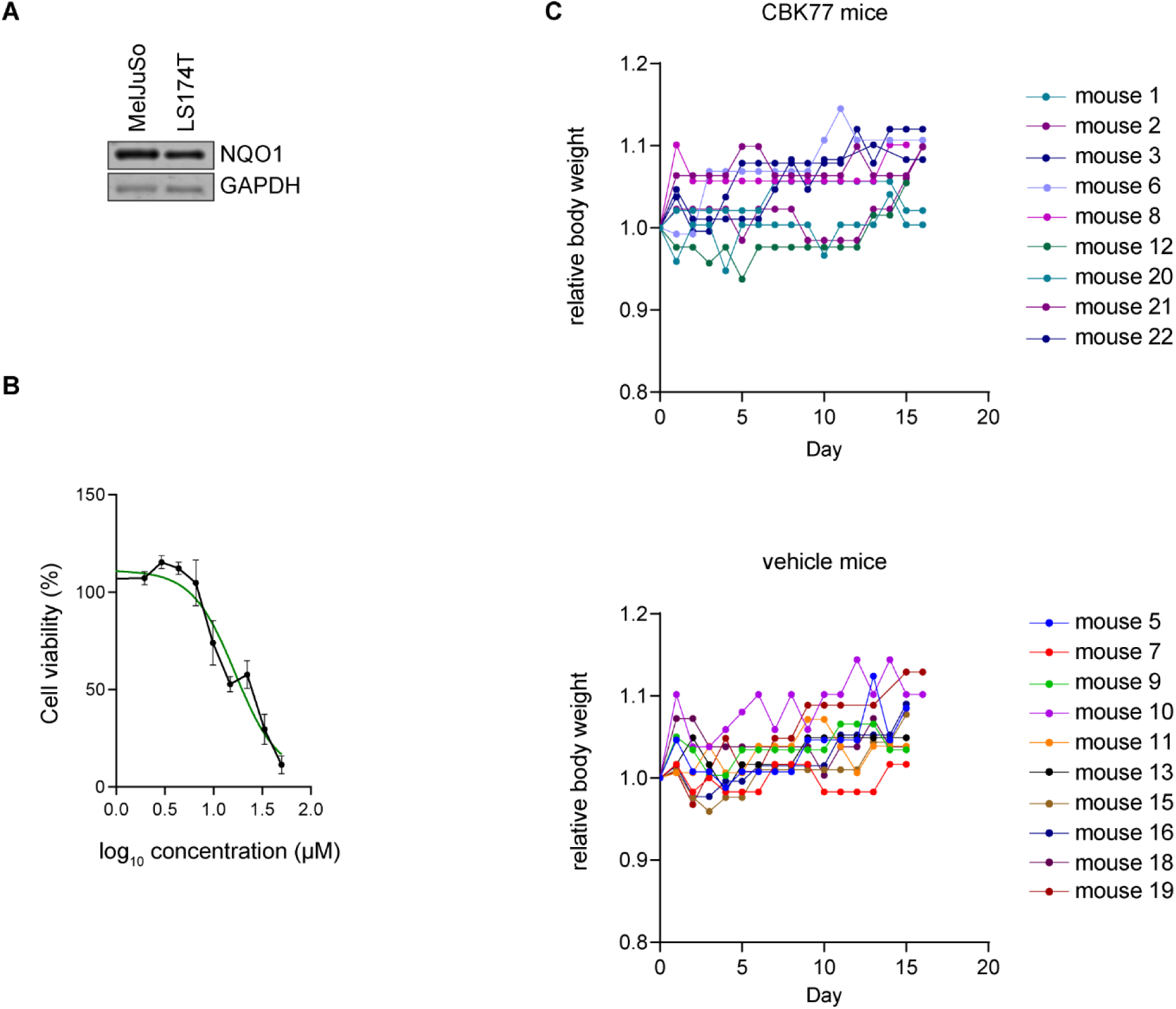
*In vivo* treatment with CBK77. **A)** Western blot analysis of cell lysates from MelJuSo and LS174T cells comparing the levels of NQO1. GAPDH is shown as a loading control. **B)** Concentration-response experiments performed with LS174T cells. Cell viability was assessed after 72 hours using the WST-1 proliferation reagent. Data are represented as mean ± SEM of five independent experiments. A curve-fit is shown in green. **C)** Mice were weighed every second day after subcutaneous injection of LS174T cells until necropsy. Weights were normalized to the weight on the day of subcutaneous cell injection.

**Supplementary Table S1.**
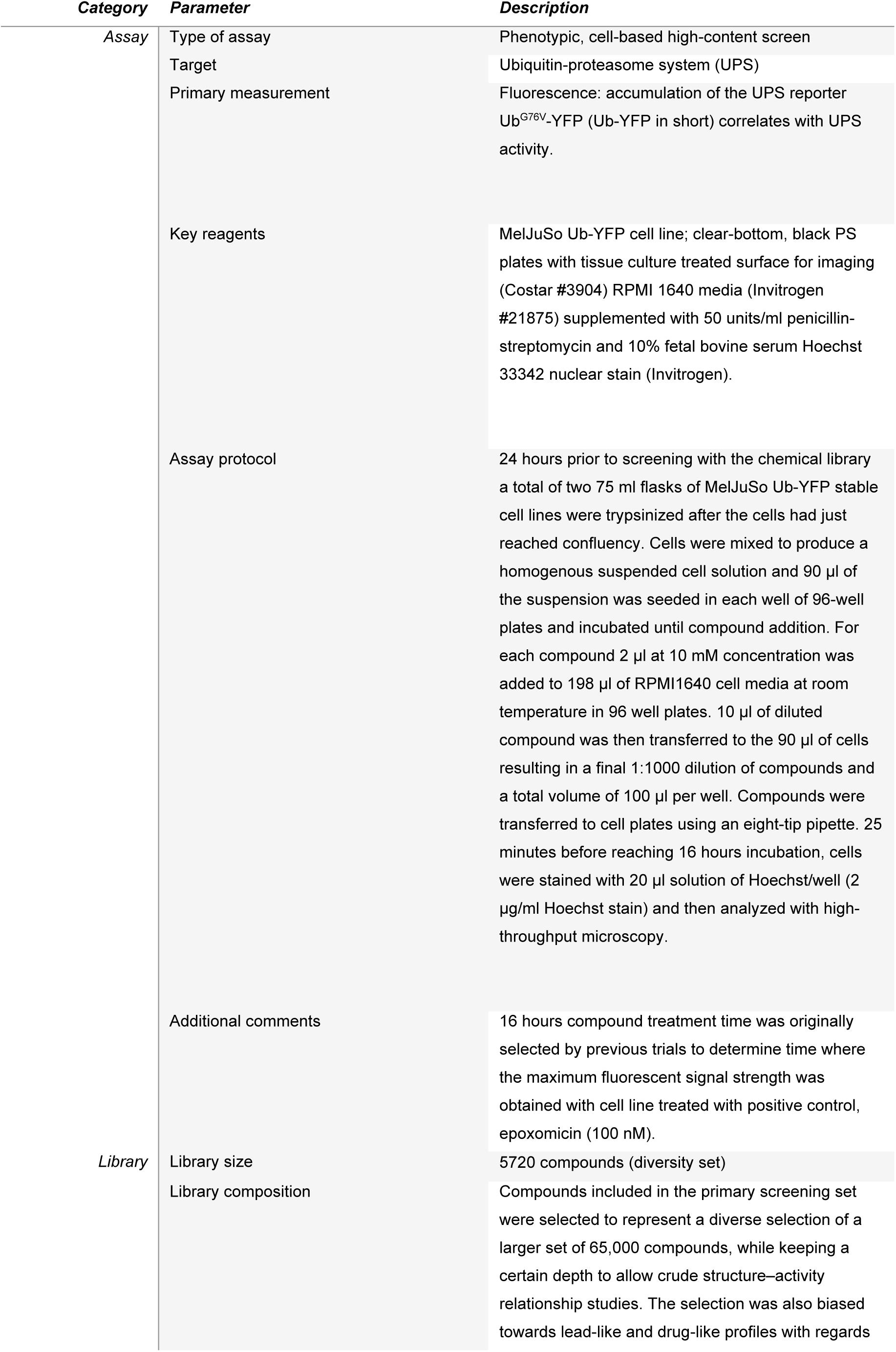
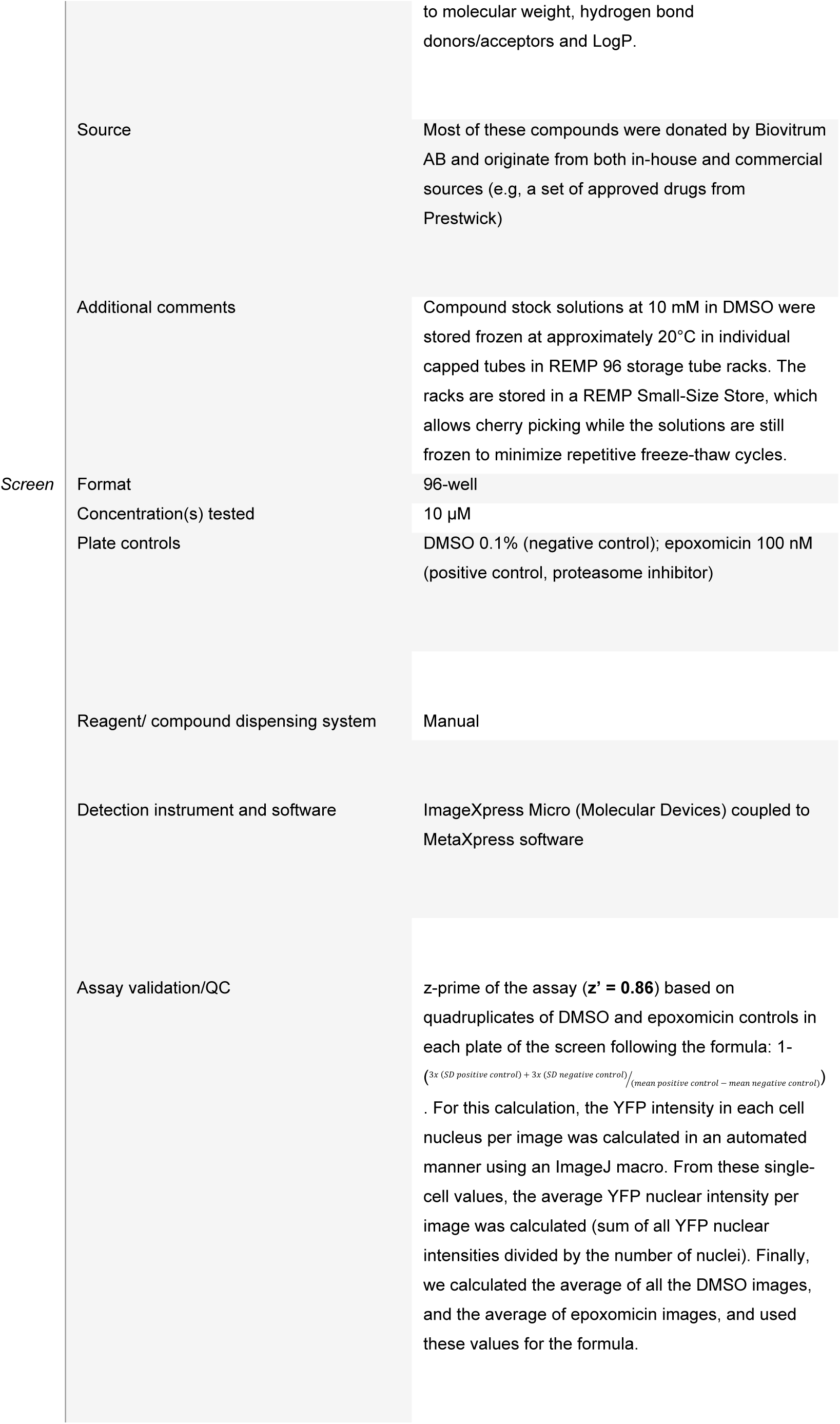
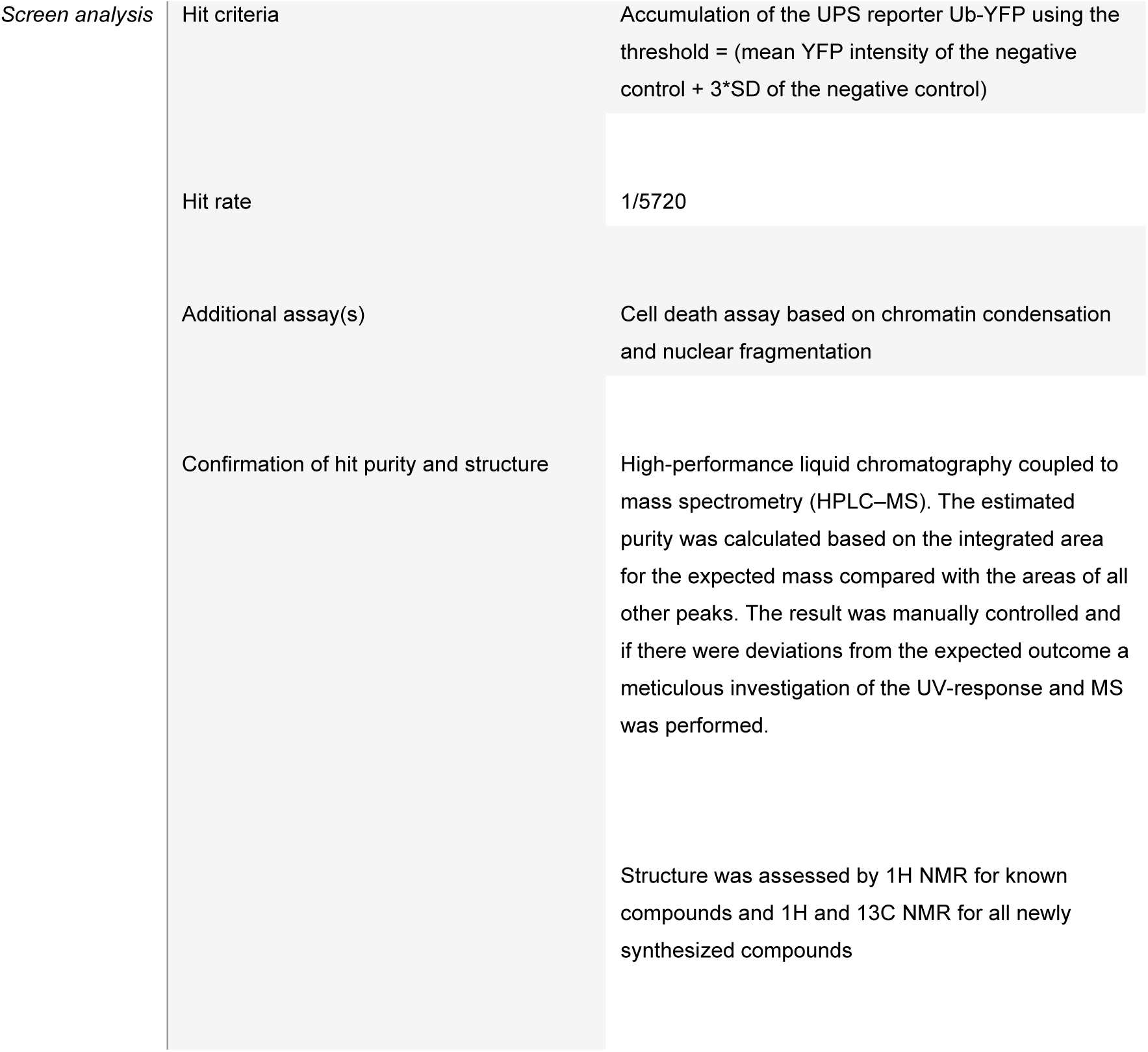
High-content screen for small molecule inhibitors of the UPS.

**Supplementary Table S2.1.**
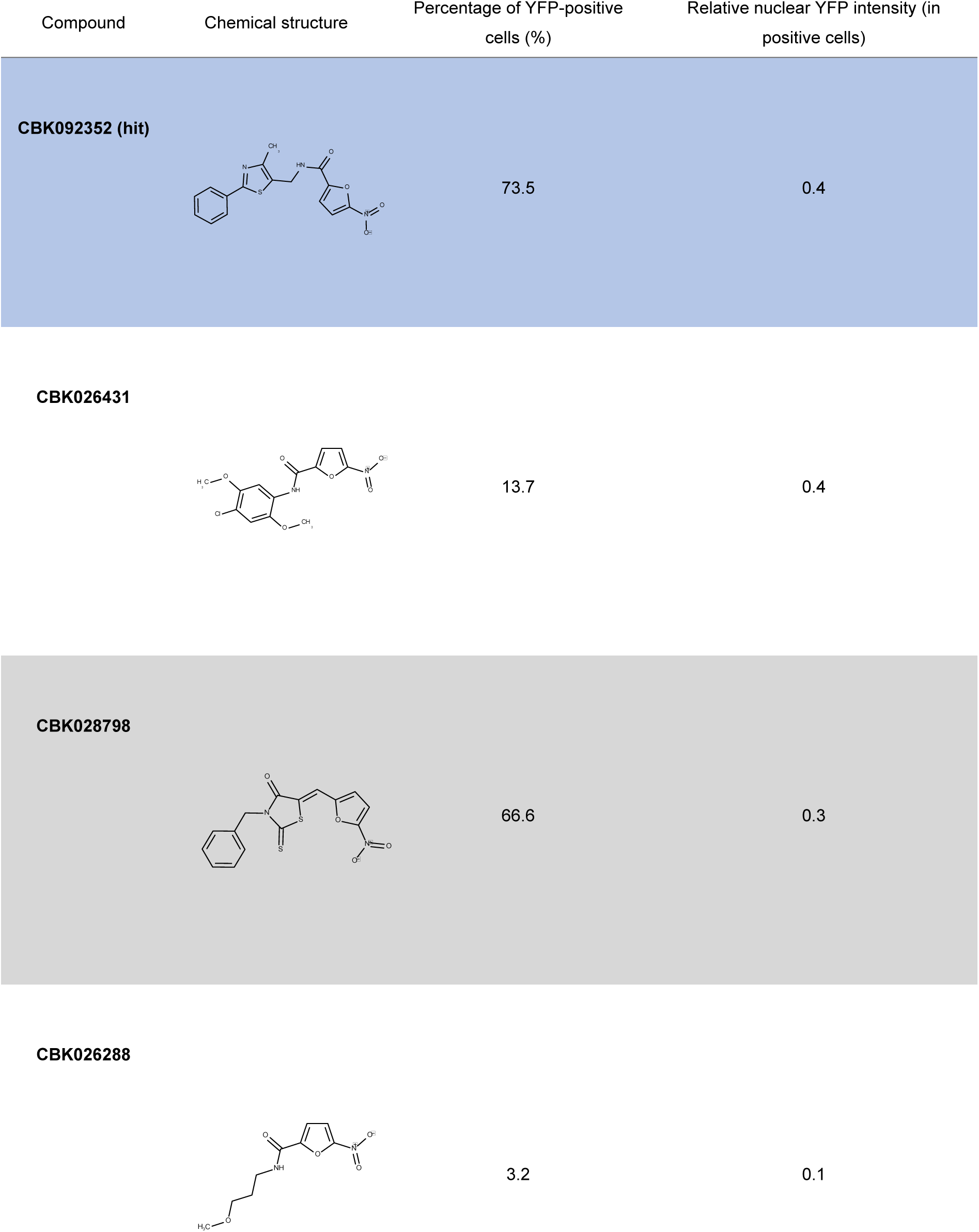
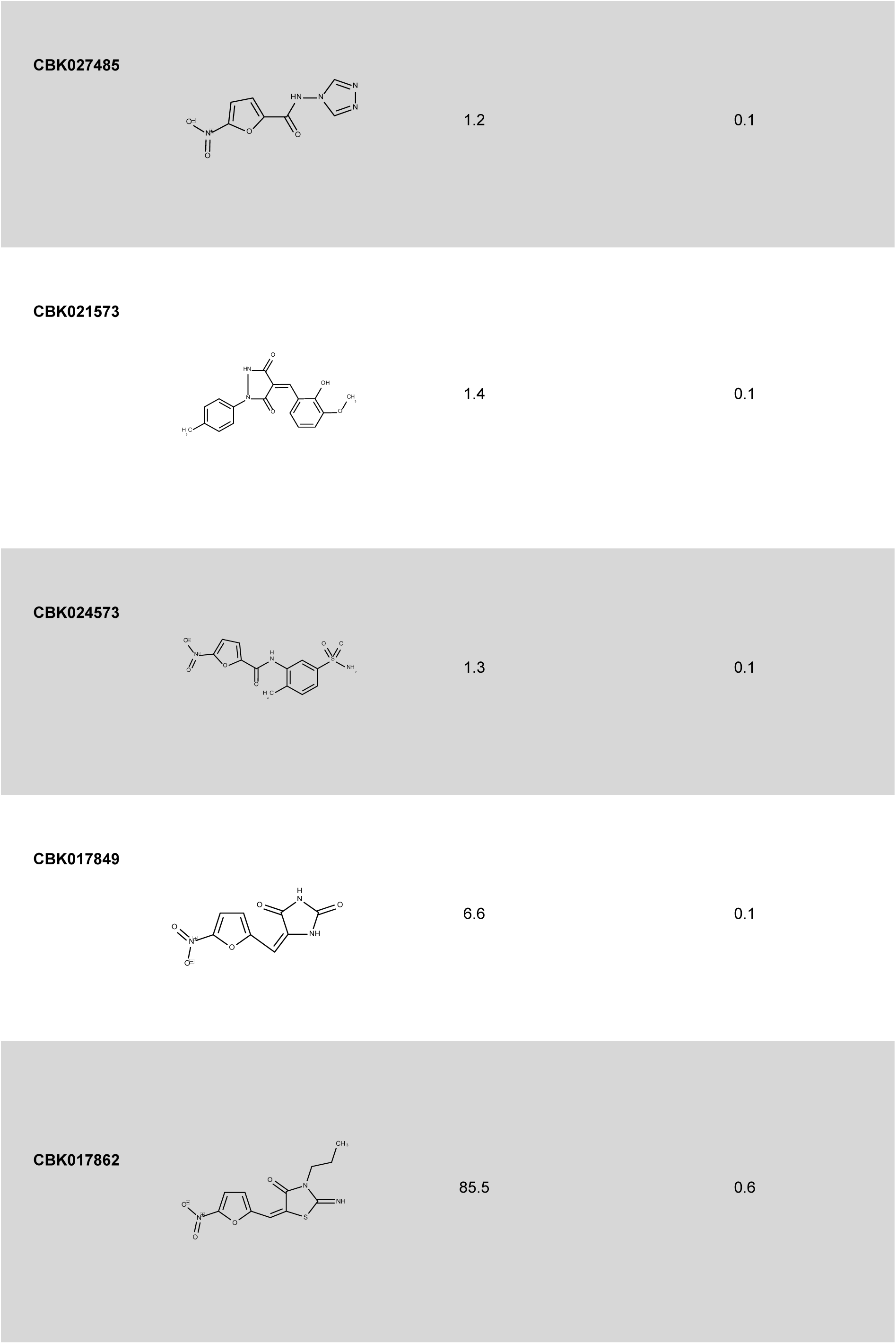
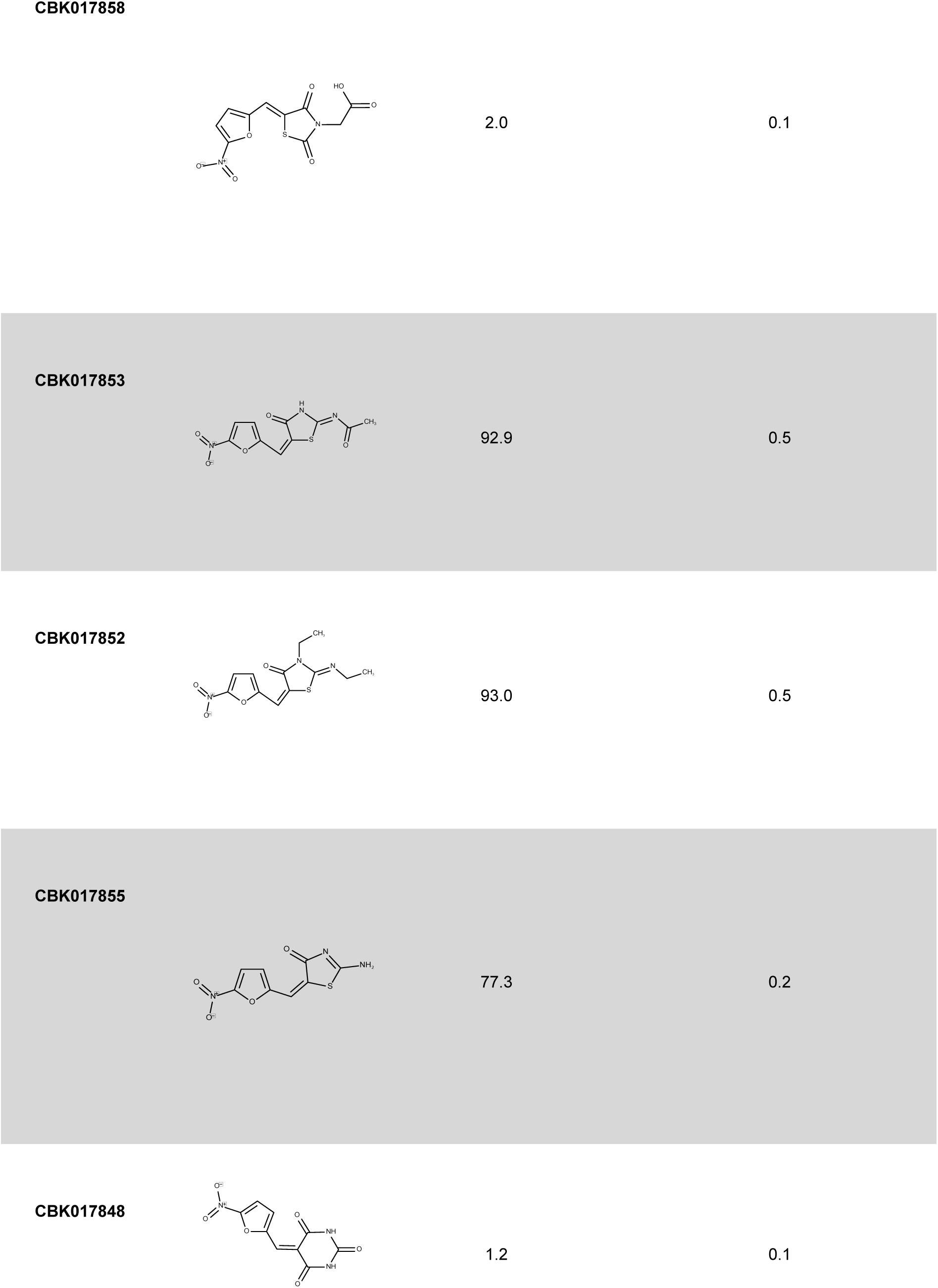
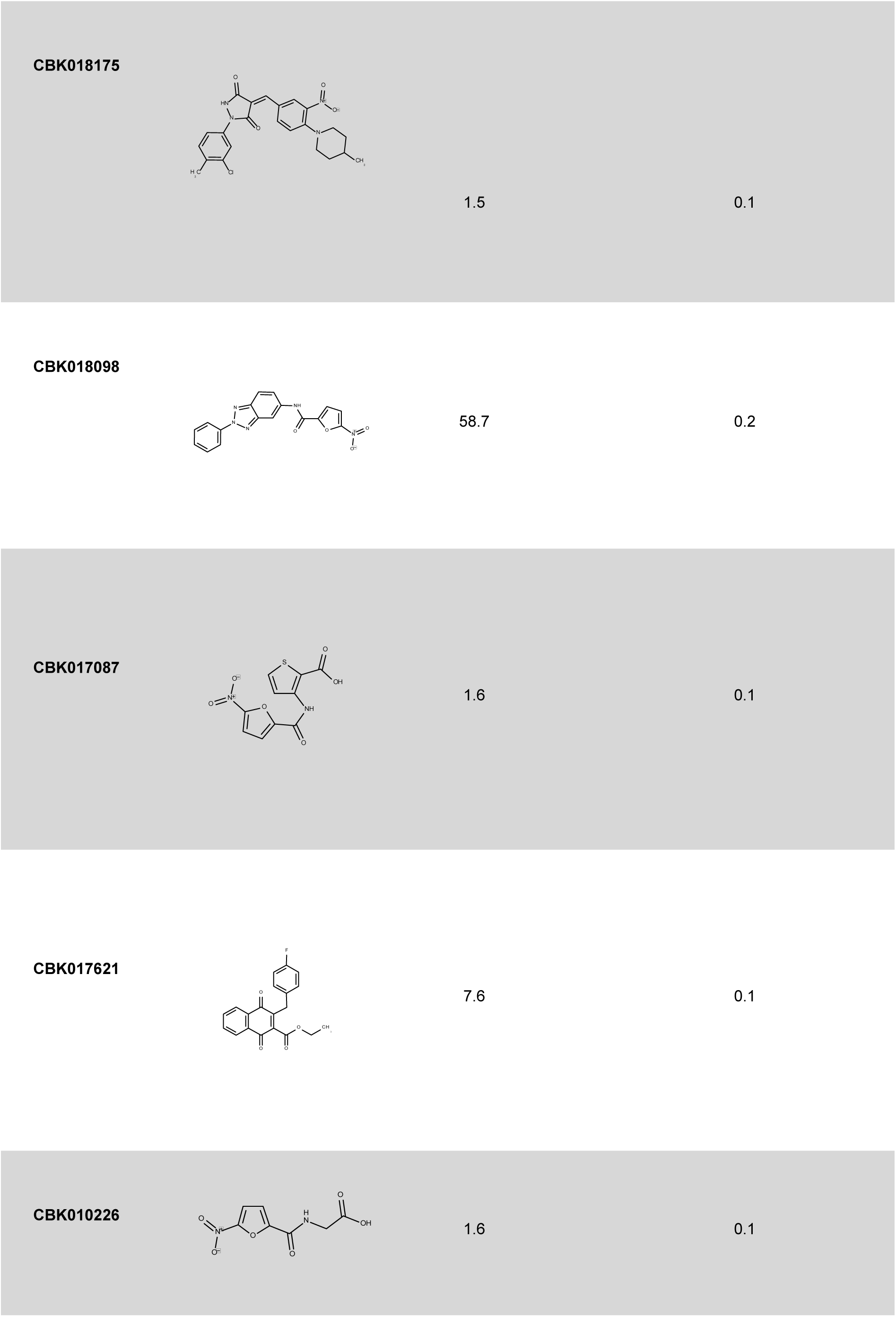
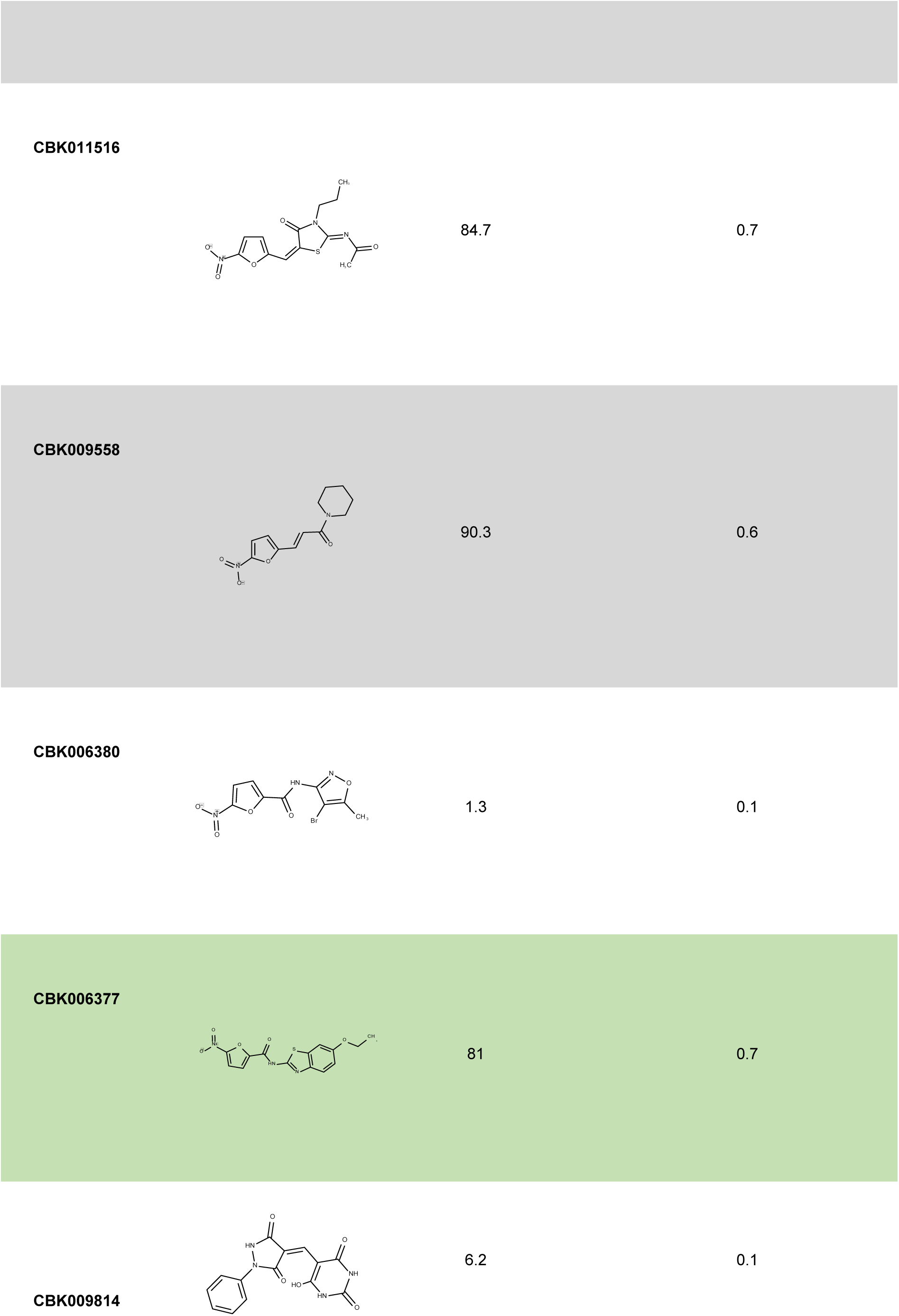
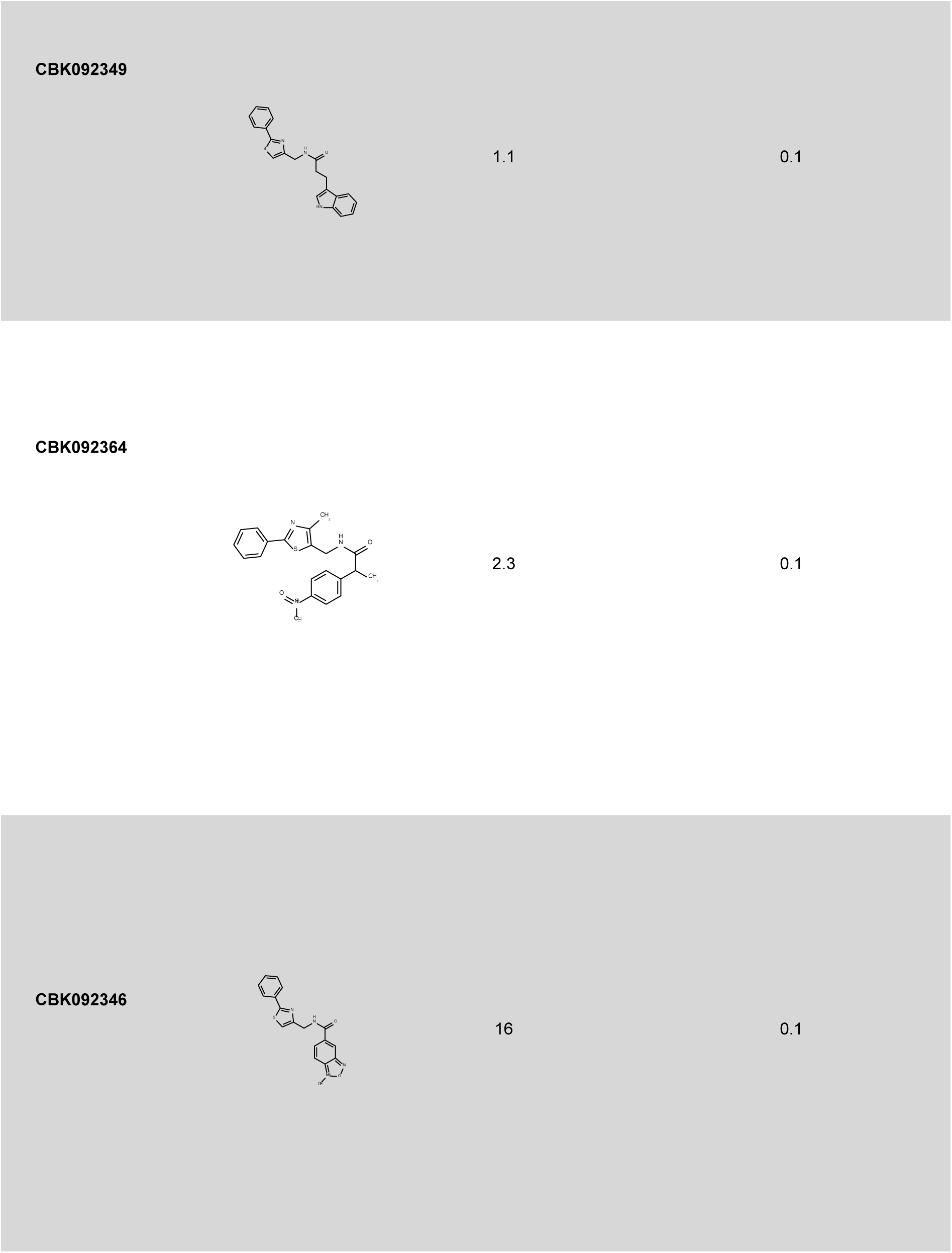
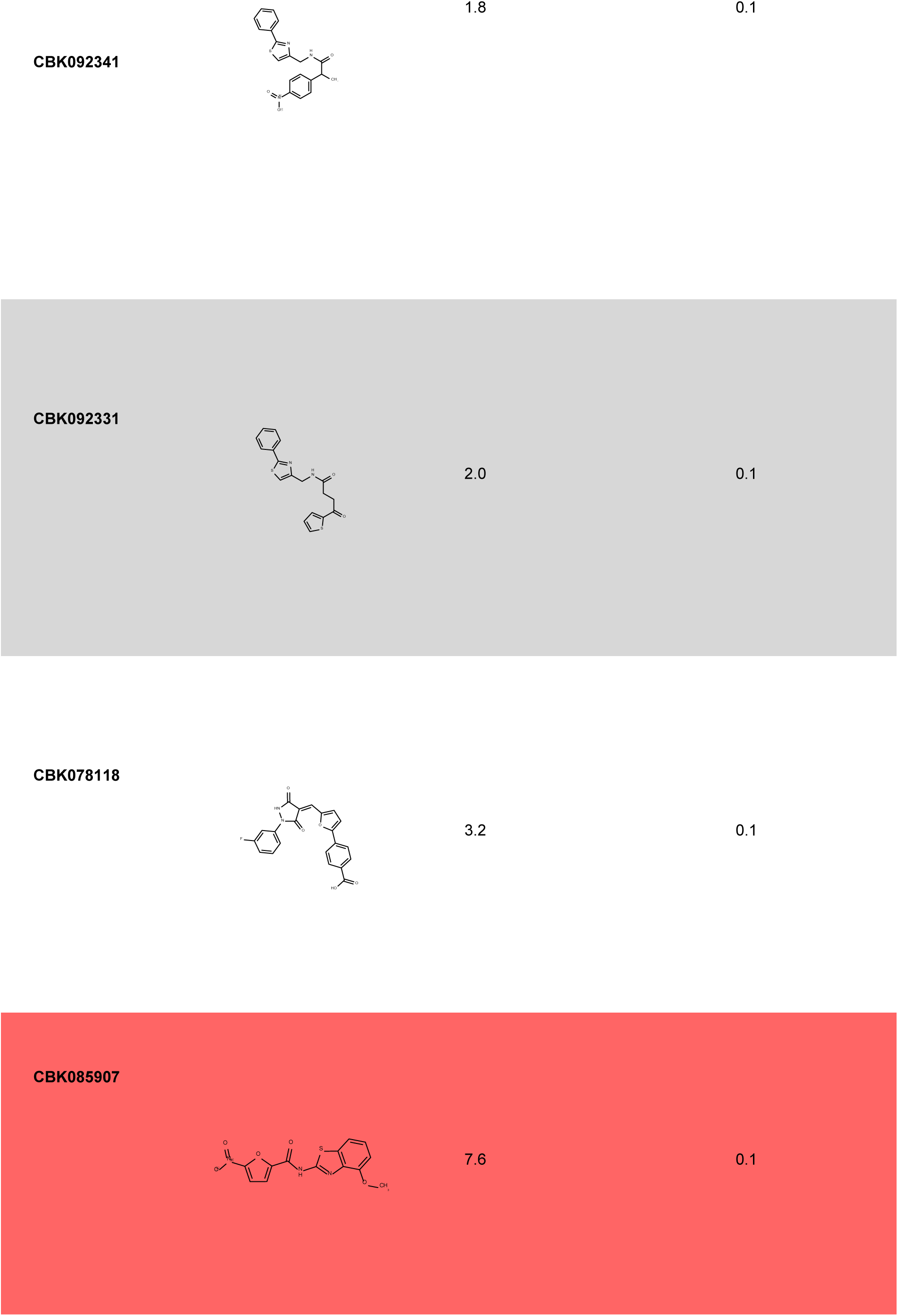
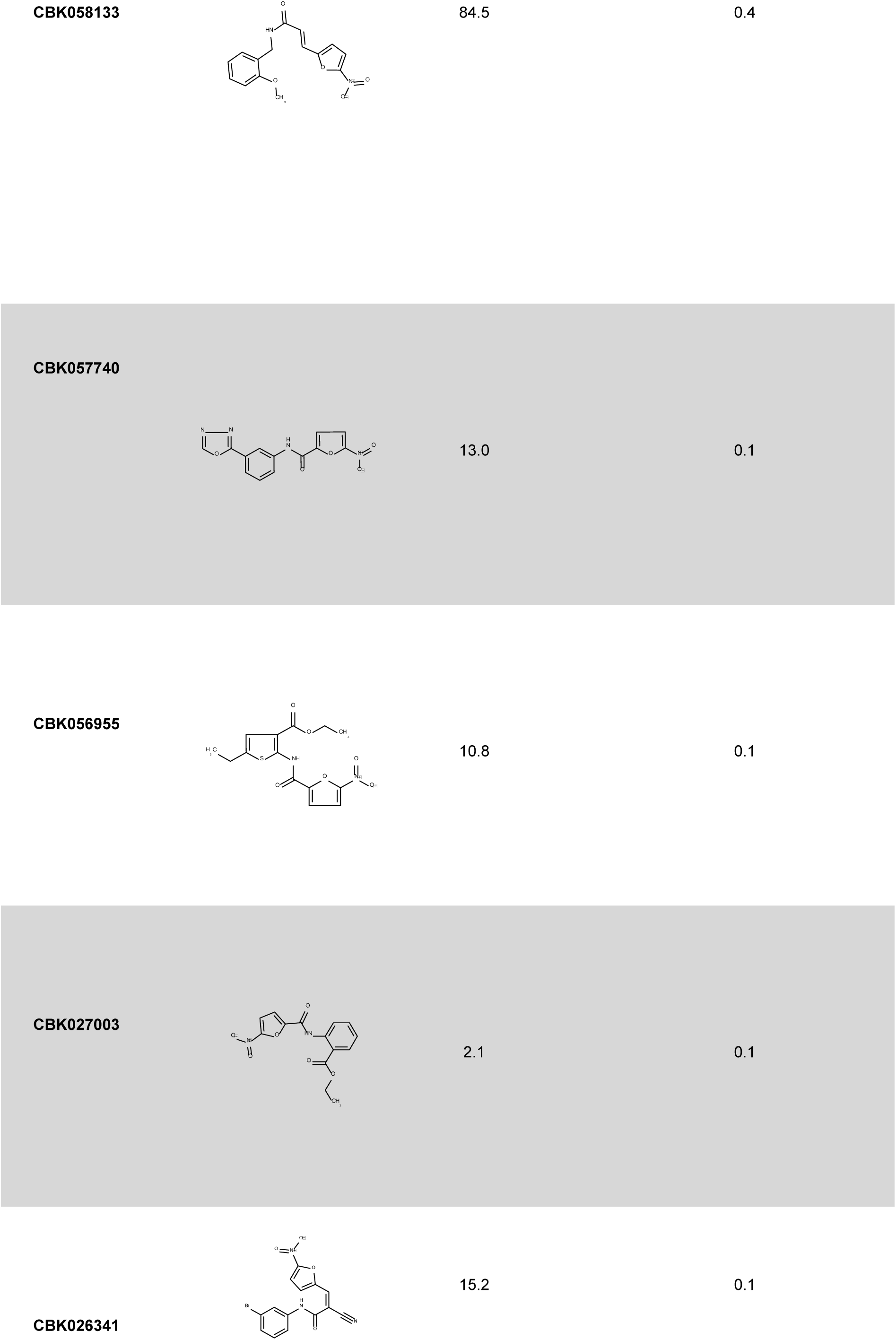
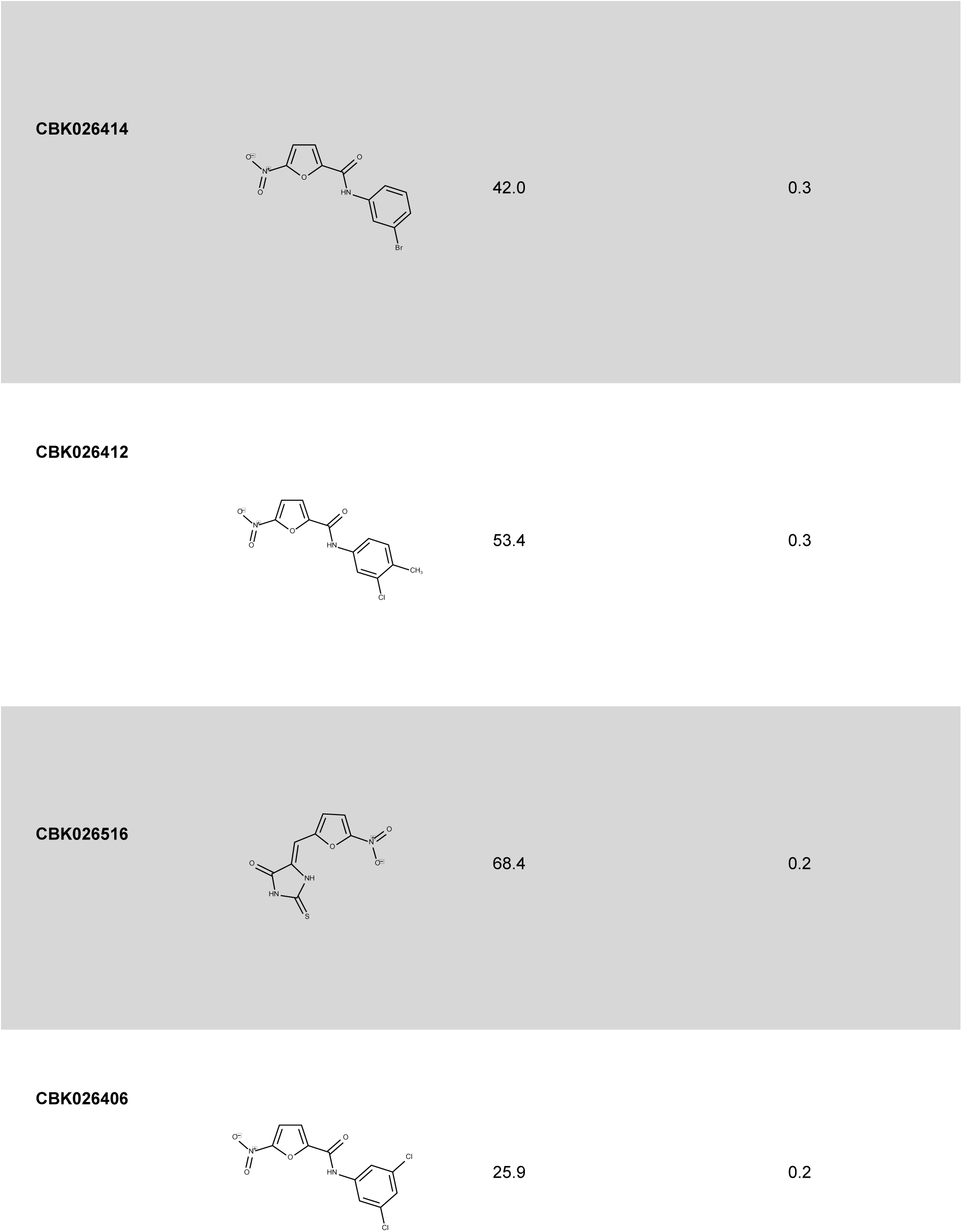
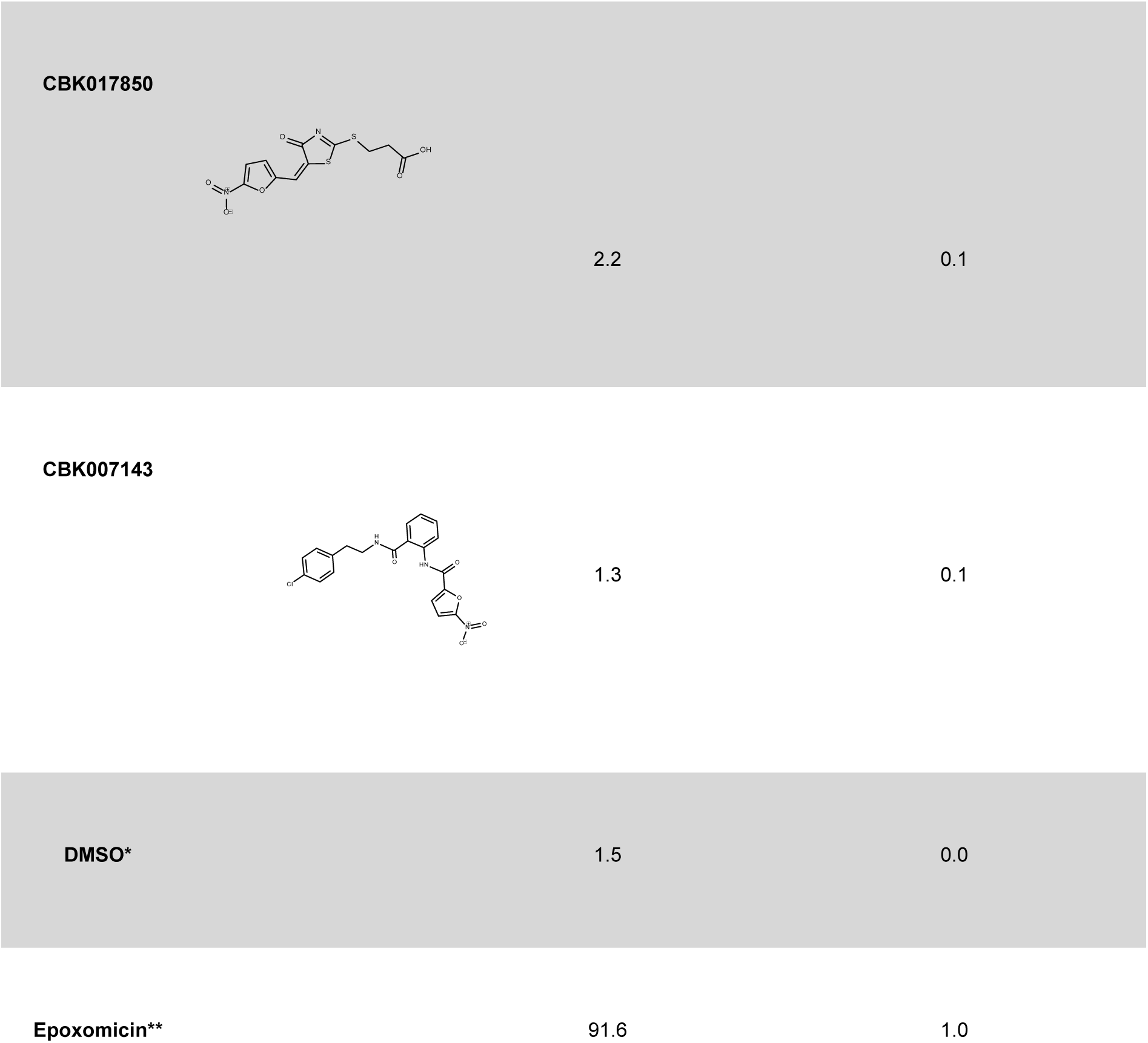
Secondary screen with structurally related compounds to the hit compound CBK092352. (as shown in Suppl. Fig. S2A). * = negative control. ** = positive control. The percentage (%) of positive cells was defined as cells with YFP levels in the nucleus above a predefined threshold (based on background fluorescence in DMSO-treated cells). The average YFP intensity in the nuclei of those cells classified as positive for YFP accumulation are shown as relative values to the positive control (epoxomicin-treated cells), set as 1.

**Supplementary Table S2.2.**
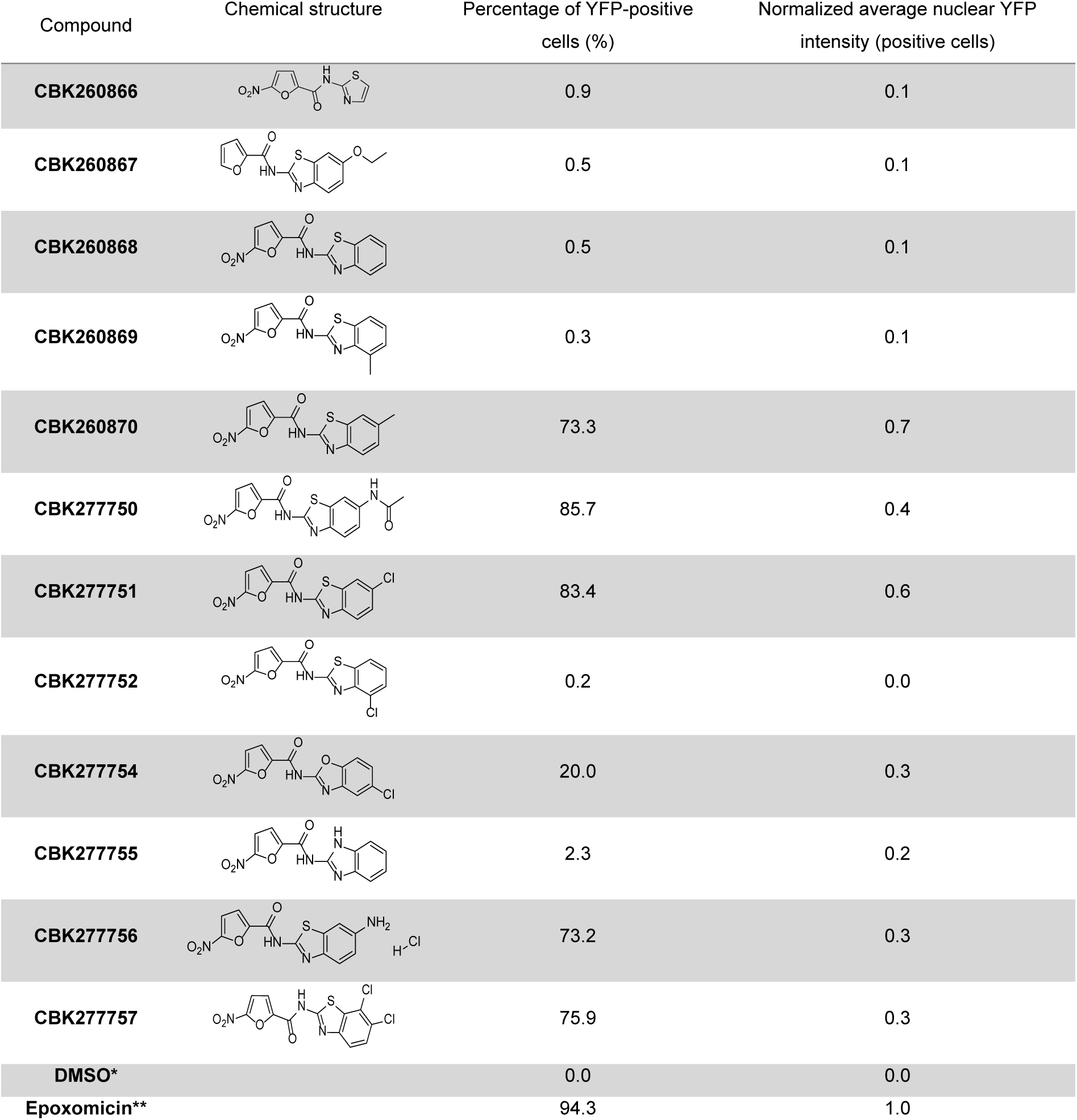
Structure-activity relationship studies. * = negative control. ** = positive control. The percentage (%) of positive cells was defined as cells with YFP levels in the nucleus above a predefined threshold (based on background fluorescence in DMSO-treated cells). The average YFP intensity in the nuclei of those cells classified as positive for YFP accumulation are shown as relative values to the positive control (epoxomicin-treated cells), set as 1.

| Compound | Chemical structure | Percentage of YFP-positive cells (%) | Normalized average nuclear YFP intensity (positive cells) |
| --- | --- | --- | --- |
| CBK260866 |  | 0.9 | 0.1 |
| CBK260867 |  | 0.5 | 0.1 |
| CBK260868 |  | 0.5 | 0.1 |
| CBK260869 |  | 0.3 | 0.1 |
| CBK260870 |  | 73.3 | 0.7 |
| CBK277750 |  | 85.7 | 0.4 |
| CBK277751 |  | 83.4 | 0.6 |
| CBK277752 |  | 0.2 | 0.0 |
| CBK277754 |  | 20.0 | 0.3 |
| CBK277755 |  | 2.3 | 0.2 |
| CBK277756 |  | 73.2 | 0.3 |
| CBK277757 |  | 75.9 | 0.3 |
| DMSO* |  | 0.0 | 0.0 |
| Epoxomicin** |  | 94.3 | 1.0 |

**Supplementary Table S2.3.**
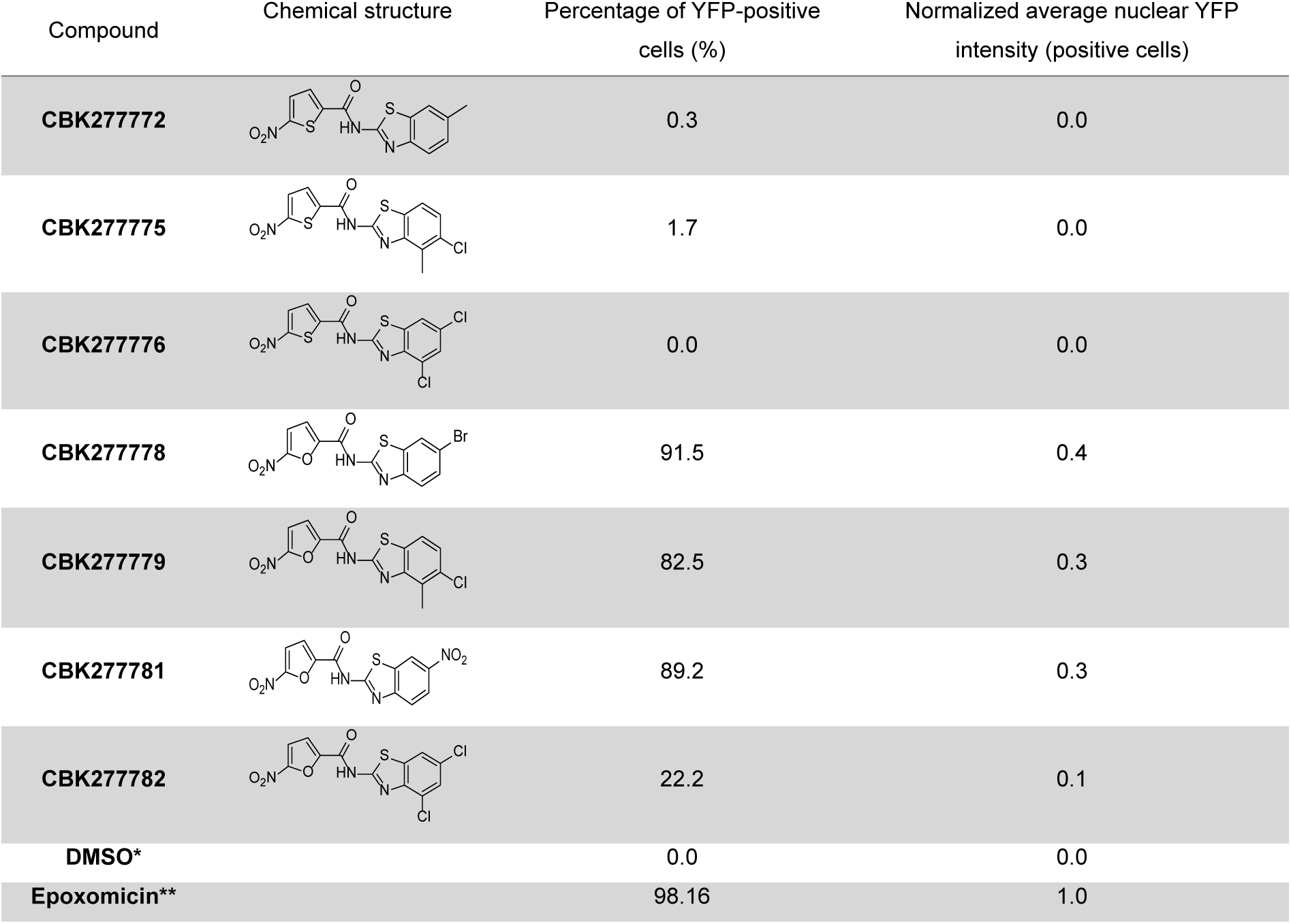
Structure-activity relationship studies. * = negative control. ** = positive control. The percentage (%) of positive cells was defined as cells with YFP levels in the nucleus above a predefined threshold (based on background fluorescence in DMSO-treated cells). The average YFP intensity in the nuclei of those cells classified as positive for YFP accumulation are shown as relative values to the positive control (epoxomicin-treated cells), set as 1.

| Compound | Chemical structure | Percentage of YFP-positive cells (%) | Normalized average nuclear YFP intensity (positive cells) |
| --- | --- | --- | --- |
| CBK277772 |  | 0.3 | 0.0 |
| CBK277775 |  | 1.7 | 0.0 |
| CBK277776 |  | 0.0 | 0.0 |
| CBK277778 |  | 91.5 | 0.4 |
| CBK277779 |  | 82.5 | 0.3 |
| CBK277781 |  | 89.2 | 0.3 |
| CBK277782 |  | 22.2 | 0.1 |
| DMSO* |  | 0.0 | 0.0 |
| Epoxomicin** |  | 98.16 | 1.0 |

**Supplementary Table S3.** Excelsheet containing the results of the two replicates performed in the CRISPR/Cas9 screen, summarized per gene.

**Supplementary Table S4.** Enriched molecular function terms in the network of proteins enriched upon CBK77 treatment in the proteomics dataset.

| Term ID | term description | Observed gene count | Background gene count | False discovery rate | Matching proteins in your network (labels) |
| --- | --- | --- | --- | --- | --- |
| GO:0015037 | peptide disulfide oxidoreductase activity | 4 | 13 | 6.10e-06 | P4HB, PDIA3, PDIA4, TXN |
| GO:0003756 | protein disulfide isomerase activity | 3 | 20 | 0.00065 | P4HB, PDIA3, PDIA4 |
| GO: 0016209 | antioxidant activity | 4 | 79 | 0.00072 | GSTP1, PRDX1, PRDX4, TXN |
| GO: 0004601 | peroxidase activity | 3 | 40 | 0.0025 | GSTP1, PRDX1, PRDX4 |
| GO: 0008379 | thioredoxin peroxidase activity | 2 | 5 | 0.0025 | PRDX1, PRDX4 |
| GO: 0043022 | ribosome binding | 3 | 50 | 0.0030 | EEF2, EIF6, RPSA |
| GO:0016853 | isomerase activity | 4 | 147 | 0.0034 | P4HB, PDIA3, PDIA4, PPIA |
| GO: 0008092 | cytoskeletal binding | 8 | 882 | 0.0036 | ANXA2, CNN3, DSTN, EEF2, HSP90AA1, MAP4, MARCKS, TPM3 |
| GO: 0051015 | actin filament binding | 4 | 158 | 0.0039 | DSTN, EEF2, MARCKS, TPM3 |
| GO: 0016491 | oxidoreductase activity | 7 | 716 | 0.0054 | GSTP1, P4HB, PDIA3, PDIA4, PRDX1, PRDX4, TXN |
| GO: 0003746 | translation elongation factor activity | 2 | 16 | 0.0074 | EEF1G, EEF2 |
| GO:0008135 | translation factor activity, RNA binding | 3 | 84 | 0.0079 | EEF1G, EEF2, EIF6 |
| GO:0003779 | actin binding | 5 | 413 | 0.0124 | CNN3, DSTN, EEF2, MARCKS, TPM3 |
| GO:000436 | glutathione transferase activity | 2 | 25 | 0.0130 | EEF1G, GSTP1 |
| GO:0005198 | structural molecule activity | 6 | 679 | 0.0160 | FLG2, MAP4, MYL6, RPSA, TPM3, TUBA1B |
| GO:0044877 | protein-containing complex binding | 7 | 968 | 0.0184 | DSTN, EEF2, EIF6, MARCKS, P4HB, RPSA, TPM3 |
| GO:0008307 | structural constituent of muscle | 2 | 45 | 0.0349 | MYL6, TPM3 |
| GO:0019205 | nucleobase-containing compound kinase activity | 2 | 49 | 0.0349 | NME2, TK1 |
| GO:0042802 | identical protein binding | 9 | 1754 | 0.0364 | ANXA2, HSP90AA1, LGALS1, MARCKS, PDIA3, PRDX1, PRDX4, S100A10, TK1 |
| GO:0050840 | extracellular matrix binding | 2 | 51 | 0.0364 | ANXA2, LGALS1 |
| GO:0019901 | protein kinase binding | 5 | 599 | 0.0394 | EEF2, GSTP1, HSP90AA1, MARCKS, PEBP1 |
| GO:0019899 | enzyme binding | 10 | 2197 | 0.0457 | ANXA2, ATOX1, EEF2, GSTP1, HSP90AA1, MARCKS, P4HB, PEBP1, TUBA1B, UBC |
| GO:0005488 | binding | 32 | 11878 | 0.0079 | ANXA2, ATOX1, CNN3, DSTN, DUT, EEF1G, EEF2, EIF6, FLG2, GSTP1, HCCS, HSP90AA1, LGALS1, MAP4, MARCKS, MYL6, NME2, P4HB, PCBP1, PDIA3, PEBP1, PRDX1, PRDX4, RPSA, S100A10, STRAP, TK1, TPM3, TUBA1B, UBC, VDAC2 |
| GO:0005515 | protein binding | 22 | 6605 | 0.0130 | ANXA2, ATOX1, CNN3, DSTN, EEF2, GSTP1, HSP90AA1, LGALS1, MAP4, MARCKS, P4HB, PDIA3, PEBP1, PPIA, PRDX1, PRDX4, S100A10, STRAP, TK1, TPM3, TUBA1B, UBC |

